# Genome wide association analysis in dilated cardiomyopathy reveals two new key players in systolic heart failure on chromosome 3p25.1 and 22q11.23

**DOI:** 10.1101/2020.02.28.969147

**Authors:** Sophie Garnier, Magdalena Harakalova, Stefan Weiss, Michal Mokry, Vera Regitz-Zagrosek, Christian Hengstenberg, Thomas P Cappola, Richard Isnard, Eloisa Arbustini, Stuart A. Cook, Jessica van Setten, Jörg Callis, Hakon Hakonarson, Michael P Morley, Klaus Stark, Sanjay K. Prasad, Jin Li, Declan P O’Regan, Maurizia Grasso, Martina Müller-Nurasyid, Thomas Meitinger, Jean-Philippe Empana, Konstantin Strauch, Mélanie Waldenberger, Kenneth B Marguiles, Christine E. Seidman, Benjamin Meder, Pierre Boutouyrie, Patrick Lacolley, Xavier Jouven, Jeanette Erdman, Stefan Blankenberg, Thomas Wichter, Volker Ruppert, Luigi Tavazzi, Olivier Dubourg, Gerard Roizes, Richard Dorent, Pascal DeGroote, Laurent Fauchier, Jean-Noël Trochu, Jean-François Aupetit, Marine Germain, Uwe Völker, Hemerich Daiane, Ibticem Raji, Delphine Bacq-Daian, Carole Proust, Kristin Lehnert, Renee Maas, Robert Olaso, Ganapathivarma Saripella, Stephan B. Felix, Steven Mc Ginn, Laëtitia Duboscq-Bidot, Alain van Mil, Céline Besse, Vincent Fontaine, Hélène Blanché, Brendan Keating, Pablo Garcia-Pavia, Angélique Curjol, Anne Boland, Michel Komajda, François Cambien, Jean-François Deleuze, Marcus Dörr, Folkert W Asselbergs, Eric Villard, David-Alexandre Trégouët, Philippe Charron, On behalf of GENMED consortium

## Abstract

We present the results of the largest genome wide association study (GWAS) performed so far in dilated cardiomyopathy (DCM), a leading cause of systolic heart failure and cardiovascular death, with 2,719 cases and 4,440 controls in the discovery population. We identified and replicated two new DCM-associated loci, one on chromosome 3p25.1 (lead SNP rs62232870, p = 8.7 × 10^−11^ and 7.7 × 10^−4^ in the discovery and replication step, respectively) and the second on chromosome 22q11.23 (lead SNP rs7284877, p = 3.3 × 10^−8^ and 1.4 × 10^−3^ in the discovery and replication step, respectively) while confirming two previously identified DCM loci on chromosome 10 and 1, *BAG3* and *HSPB7*. The genetic risk score constructed from the number of lead risk-alleles at these four DCM loci revealed that individuals with 8 risk-alleles were at a 27% increased risk of DCM compared to individuals with 5 risk alleles (median of the referral population). We estimated the genome wide heritability at 31% ± 8%.

*In silico* annotation and functional 4C-sequencing analysis on iPSC-derived cardiomyocytes strongly suggest *SLC6A6* as the most likely DCM gene at the 3p25.1 locus. This gene encodes a taurine and beta-alanine transporter whose involvement in myocardial dysfunction and DCM is supported by recent observations in humans and mice. Although less easy to discriminate the better candidate at the 22q11.23 locus, *SMARCB1* appears as the strongest one.

This study provides both a better understanding of the genetic architecture of DCM and new knowledge on novel biological pathways underlying heart failure, with the potential for a therapeutic perspective.

## Introduction

Dilated cardiomyopathy (DCM) is a heart muscle disease characterized by left ventricular dilatation and systolic dysfunction in the absence of abnormal loading conditions or coronary artery disease (CAD)(1,2). It is a major cause of systolic heart failure, the leading indication for heart transplantation, and therefore a major public health problem due to the important cardiovascular morbidity and mortality(1,2). The understanding of the genetic basis of DCM has improved during the past years with the role of both rare and common variants resulting in a complex genetic architecture of the disease(3,4). More than 50 genes(4–6) with rare pathogenic mutations have been reported as causing DCM, mainly inherited as dominant with variable penetrance. Several large scale association studies in sporadic cases have been performed to identify common DCM-associated alleles(7–13) including four genome wide association studies (GWAS)(9,10,12,13). Altogether, these genetic investigations have so far robustly identified 2 loci harboring common susceptibility alleles: a locus on chromosome 1, encompassing multiple candidate genes in strong linkage disequilibrium (LD), including *ZBZTB17/MIZ-1, HSPB7* and *CLCNKA*(7–12); and a second locus on chromosome 10 for which the culprit gene, *BAG3*, is also involved in familial forms of DCM(9,14). The analysis of 116,855 common coding variants in an exome wide association study (EWAS) has also suggested the existence of six potential additional DCM loci(11) but, in absence of proper replication, their true contribution to DCM has not been definitively established. Here, we report the results of a new GWAS for sporadic DCM performed on 2,719 cases and 4,440 controls. Individual genotype data were imputed for the 1000Genomes reference panel and the main findings replicated in two independent case-control samples totaling 584 cases and 986 controls. Then, *in silico* annotation and functional analyses were performed in candidate loci to identify the better candidates. We also estimated the global genetic heritability, and built a genetic risk score (GRS) for DCM.

## Material and Methods

### Populations inclusion and samples collection

DCM patients and controls from five populations (France, Germany, USA, Italy and UK) were included in the GWAS (a detailed description of the cohorts is provided in **Supplemental Material**). Sporadic DCM was diagnosed according to standard criteria(1,4,15,16) by reduced ejection fraction (EF, echocardiography: <45% or MRI: <2 standard deviations (SDs) below the age- and sex-adjusted mean (16)) and an enlarged left ventricle end-diastolic volume/diameter (LVEDD >117% of value predicted from age and body surface area on echocardiography, or >2 SDs from the age- and sex-adjusted mean by MRI(16)) in the absence of significant coronary artery disease or intrinsic valvular disease, documented myocarditis, systemic disease, sustained arterial hypertension, or congenital malformation. All patients signed informed consent and the study protocol was approved by local ethics committees. In total, 2,719 cases and 4,440 controls were included in the discovery GWAS analysis.

Two case-control samples were available for replication of the main discovery GWAS findings, a Dutch population composed of 145 DCM cases and 527 controls(17) and a German collection of 439 patients with left ventricle dilation and/or hypokinetic non-dilated cardiomyopathy (HNDC) and 439 controls(18,19) (detailed descriptions are provided in **Supplemental Material**).

### Genotyping, genotype calling, and imputation

Genotyping was performed with high-density arrays for all samples and was further imputed with the 1,000 Genomes reference dataset. Summary descriptions of specific genotyping arrays, QC filtering, and imputation methods are given in **Supplemental Material** and **Supplementary Table 1**.

### Association analysis in the discovery phase

Association of imputed SNPs with DCM was performed using the Mach2dat software(20,21) implementing a logistic regression model adjusted for sex and genome-wide genotype-derived principal components under the assumption of additive allele effects. Only bi-allelic SNPs with imputation quality, r^2^, greater than 0.5 and minor allele frequency (MAF) higher than 0.005 were kept for association analysis. A statistical threshold of 5 × 10^−8^ was used to declare genome-wide significance.

To check the existence of several independent DCM-associated SNPs at loci with genome-wide statistical significance, we conducted conditional analyses adjusting for the lead SNP at each identified locus. In the case of more than one significant SNP at a given locus, additional haplotype analyses were conducted using the THESIAS software(22).

### Association analysis in the replication phase

SNPs selected for replication were tested using the same statistical model as that used in the discovery stage, separately in each of the two replication cohorts, adopting a one-tailed hypothesis and applying a Bonferroni correction procedure to declare statistical replication while controlling for the number of tested SNPs. Results of the replication studies were subsequently meta-analyzed using a fixed-effects model based on the inverse-weighting method as implemented in the Metal software(23). A similar meta-analysis framework was applied to combine the results of the discovery and replication phase. Heterogeneity across studies (between the two replication studies or between the discovery and replication stage) was tested using the Cochran’s Q statistic and the I^2^ index was used to express its magnitude.

### Regional association plot

At each of the replicated associated genetic loci, a regional association plot was performed using the LocusZoom online tool (http://locuszoom.sph.umich.edu/).

### Genetic risk score analysis

A genetic risk score (GRS) was built upon the two SNPs already known to associate with DCM, *HSPB7-*rs10927886(7–12) and *BAG3*-rs2234962(9,14) and the independent replicated genome-wide significant DCM associated SNPs. That score was tested, using logistic regression analysis, for association with DCM on a continuous scale and with quintile repartition. For each individual i, the GRS was defined as 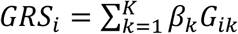, where K is the number of variants composing the score, *β*_*k*_ is the log-odds ratio for DCM associated with variant k obtained in the replication phase and *G*_*ik*_ is the allele dosage at variant k for individual i.

### Genetic heritability

The linkage disequilibrium (LD) score regression approach(24) was used to get an estimate of the genome-wide genetic heritability underlying DCM. We also used this methodology to calculate the genetic correlation between DCM and several cardiovascular traits capitalizing on the GWAS results available at the LD Hub (http://ldsc.broadinstitute.org/ldhub/).

### Candidates selection strategy at associated loci

For each newly identified DCM locus, a downstream fine-mapping strategy was deployed using *in silico* and experimental functional data to select the best candidates at each locus.

#### Cis-regulation of associated SNPs

At each locus, DCM associated SNPs (p-value ≤ 5 × 10^−8^ and/or presenting a high LD (r^2^>0.7) with the lead SNP) were used to define the associated “LD block”. We checked whether those blocks could overlap with DNA regulatory elements by using the UCSC Genome Browser(25) in human assembly hg19 (http://genome.ucsc.edu/) by visualizing the ENCODE3 DNase hypersensitivity sites (DHS)(26) and transcription factor (TF) chromatin immunoprecipitation sequencing (ChIP-seq)(27) tracks produced on 125 and 130 cell lines, respectively. To detect left ventricle (LV)-specific putative regulatory regions we enriched those tracks with H3K27ac (active chromatin), H3K4me1 (enhancer) and H3K4me3 (promotor) histone marks of ENCODE heart LV samples (GSM910575, GSM910580, GSM908951)(28). To support the potential regulatory/promotor role of those regions, we looked at regulatory elements predicted from ORegAnno(29) and checked sequence conservation in a subset of the vertebrates(30).

#### Topologically associating domains (TAD) and intra-TAD chromatin interactions

TADs are domains of preferential chromatin interaction separated by insulators where genes are accessible to intra-TAD regulatory elements, such as enhancers(31). Taking advantage of public resources describing those TADs, we delimited the region comprising the LD block, thus the subset of candidate genes, that are jointly regulated. Regional candidate genes were defined based on published LV TADs described in Leung *et al.*(28), as well as preferential chromatin interaction measured *via* promoter chromatin Hi-C (PCHi-C) on iPSC-derived cardiomyocytes (iPSC-CM) by Montefiori *et al.*(32).

Those published TAD boundaries were confirmed by in-house circular chromatin conformation capture (4C)-sequencing data produced on an iPSC-derived cardiomyocyte line from a donor (a full description is enclosed in the **Supplemental Material**).

#### Candidate genes’ biological insights

The heart expression level of each candidate gene in an associated region was evaluated *via* RNA-sequencing cardiac expression data issued from the Genotype-Tissue Expression (GTEx) project database22 (http://www.gtexportal.org/home/) (LV and atrial appendage) and from LV explants of DCM cases and controls produced by Henig *et al*.(33). That later work also enabled us to check whether differential expression existed between the 97 DCM patients and the 108 healthy donors. For the subset of gene displaying interesting expression features, we then scrutinized publicly available resources for gene annotation and functions.

#### Annotation of associated SNPs

The annotation of SNPs’ blocks was realized with Annovar software(34) and bioinformatics prediction of effects using RegulomeDB(35), Regulatory Mendelian mutation (ReMM)(36) and the GRCh37-v1.4 Combined Annotation Dependent Depletion (CADD) model(37) (https://cadd.gs.washington.edu/snv). Each SNP of an association block being prone to be implicated in an alteration of local regulatory function, we went through various *in silico* data in an effort to identify regulatory ones. We thus screened GTEx database for expression and splicing quantitative trait loci (eQTL and sQTL) in human cardiac and skeletal muscle tissues, checked whether some of those SNPs could be associated with blood DNA methylation levels (mQTL)(38) and looked at their location in putative enhancer or promotor region (H3K27ac, H3K4me1, and H3K4me3 histone marks).

## Results

### Main statistical findings

After quality controls, 9,152,885 SNPs (including 8,945,131 autosomal and 207,754 X chromosome SNPs) were tested for association with DCM in a total of 2,651 cases and 4,329 controls. Results of the discovery GWAS are summarized in a Manhattan and a Quantile-Quantile plot (**Figure 1** and **Supplementary Figure 1**). The main findings are summarized in **Table 1**. Five loci reached genome-wide significance. Two were already known to associate with DCM, *BAG3* (p = 4.7 × 10^−14^ for rs61869036) and *HSPB7* (p-value = 2.12 10^−13^ for rs10927886). Of note, the *BAG3* rs61869036 is in complete LD (r^2^∼1) with the nonsynonymous rs2234962 already reported to associate with DCM(9,11) that was thus further used as *BAG3* lead SNP (p = 5.6 × 10^−14^). Three new loci reached genome-wide significance on chr3p25.1 (rs62232870, p = 8.7 × 10^−11^) downstream *LSM3*, on chr16p13.3 (*PKD1* rs2519236, p = 3.0 10^−8^) and on chr22q11.23 (*SMARCB1* rs7284877, p = 3.3 10^−8^). Regional association plots at these 5 loci are shown in **Supplementary Figures 2-6**.

**Table 1.**
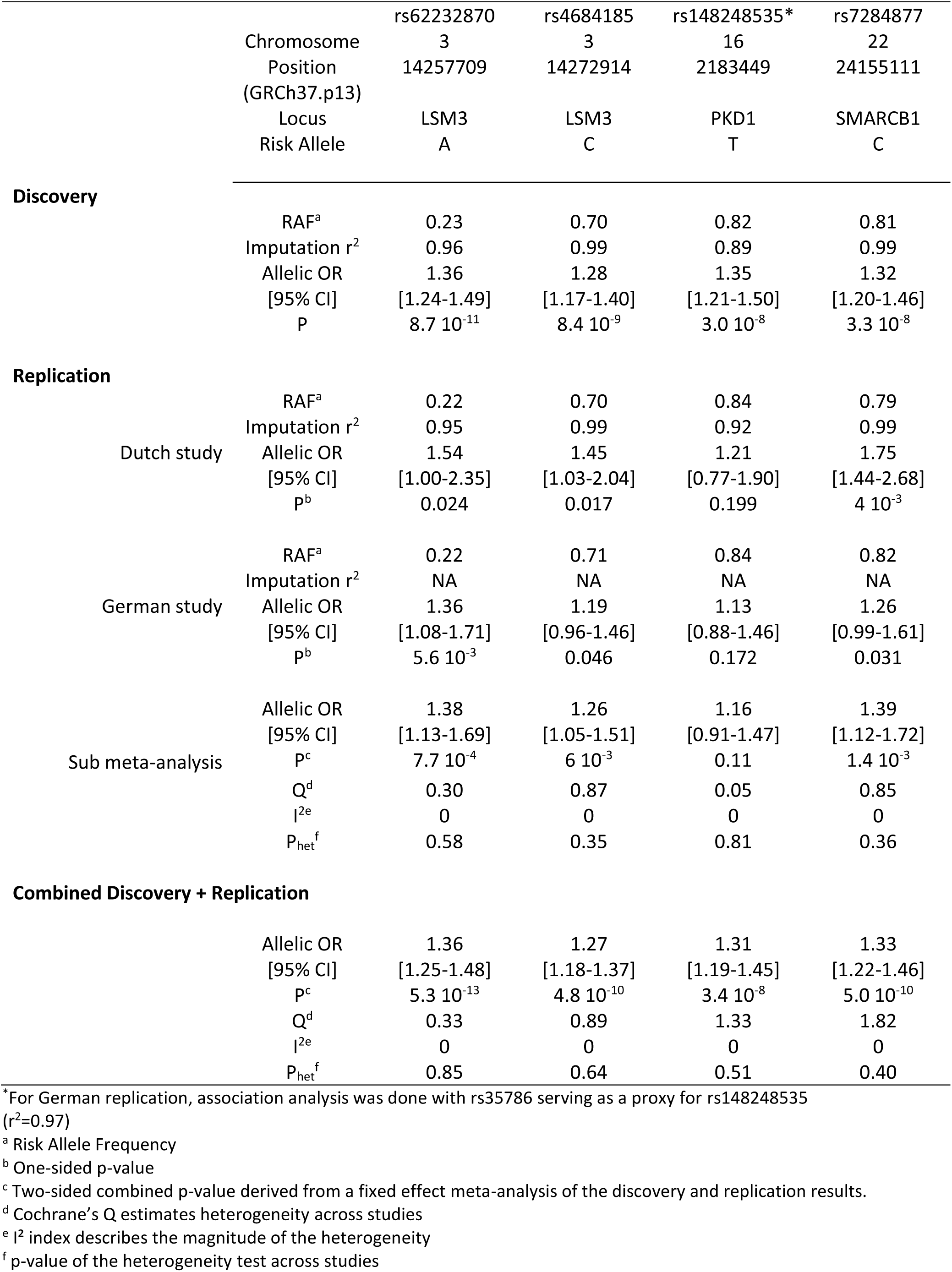
Main association findings of the DCM GWAS results.

|  |  | rs62232870 | rs4684185 | rs148248535* | rs7284877 |
| --- | --- | --- | --- | --- | --- |
|  | Chromosome | 3 | 3 | 16 | 22 |
|  | Position<br>(GRCh37.p13) | 14257709 | 14272914 | 2183449 | 24155111 |
|  | Locus | LSM3 | LSM3 | PKD1 | SMARCB1 |
|  | Risk Allele | A | C | T | C |
| **Discovery** |  |  |  |  |  |
|  | RAF<sup>a</sup> | 0.23 | 0.70 | 0.82 | 0.81 |
| | Imputation $r^2$ | 0.96 | 0.99 | 0.89 | 0.99 |
|  | Allelic OR | 1.36 | 1.28 | 1.35 | 1.32 |
|  | [95% CI] | [1.24-1.49] | [1.17-1.40] | [1.21-1.50] | [1.20-1.46] |
| | P | $8.7 \cdot 10^{-11}$ | $8.4 \cdot 10^{-9}$ | $3.0 \cdot 10^{-8}$ | $3.3 \cdot 10^{-8}$ |
| **Replication** |  |  |  |  |  |
| Dutch study | RAF<sup>a</sup> | 0.22 | 0.70 | 0.84 | 0.79 |
| | Imputation $r^2$ | 0.95 | 0.99 | 0.92 | 0.99 |
|  | Allelic OR | 1.54 | 1.45 | 1.21 | 1.75 |
|  | [95% CI] | [1.00-2.35] | [1.03-2.04] | [0.77-1.90] | [1.44-2.68] |
| | $p^b$ | 0.024 | 0.017 | 0.199 | $4 \cdot 10^{-3}$ |
| German study | RAF<sup>a</sup> | 0.22 | 0.71 | 0.84 | 0.82 |
| | Imputation $r^2$ | NA | NA | NA | NA |
|  | Allelic OR | 1.36 | 1.19 | 1.13 | 1.26 |
|  | [95% CI] | [1.08-1.71] | [0.96-1.46] | [0.88-1.46] | [0.99-1.61] |
| | $p^b$ | $5.6 \cdot 10^{-3}$ | 0.046 | 0.172 | 0.031 |
| Sub meta-analysis | Allelic OR | 1.38 | 1.26 | 1.16 | 1.39 |
|  | [95% CI] | [1.13-1.69] | [1.05-1.51] | [0.91-1.47] | [1.12-1.72] |
| | $P^c$ | $7.7 \cdot 10^{-4}$ | $6 \cdot 10^{-3}$ | 0.11 | $1.4 \cdot 10^{-3}$ |
| | $Q^d$ | 0.30 | 0.87 | 0.05 | 0.85 |
| | $I^2^e$ | 0 | 0 | 0 | 0 |
| | $P_{het}^f$ | 0.58 | 0.35 | 0.81 | 0.36 |
| **Combined Discovery + Replication** |  |  |  |  |  |
|  | Allelic OR | 1.36 | 1.27 | 1.31 | 1.33 |
|  | [95% CI] | [1.25-1.48] | [1.18-1.37] | [1.19-1.45] | [1.22-1.46] |
| | $P^c$ | $5.3 \cdot 10^{-13}$ | $4.8 \cdot 10^{-10}$ | $3.4 \cdot 10^{-8}$ | $5.0 \cdot 10^{-10}$ |
| | $Q^d$ | 0.33 | 0.89 | 1.33 | 1.82 |
| | $I^2^e$ | 0 | 0 | 0 | 0 |
| | $P_{het}^f$ | 0.85 | 0.64 | 0.51 | 0.40 |

\*For German replication, association analysis was done with rs35786 serving as a proxy for rs148248535 ( $r^2=0.97$ )
<sup>a</sup> Risk Allele Frequency
<sup>b</sup> One-sided p-value
<sup>c</sup> Two-sided combined p-value derived from a fixed effect meta-analysis of the discovery and replication results.
<sup>d</sup> Cochran's Q estimates heterogeneity across studies
<sup>e</sup> $I^2$ index describes the magnitude of the heterogeneity
<sup>f</sup> p-value of the heterogeneity test across studies

**Figure 1.**
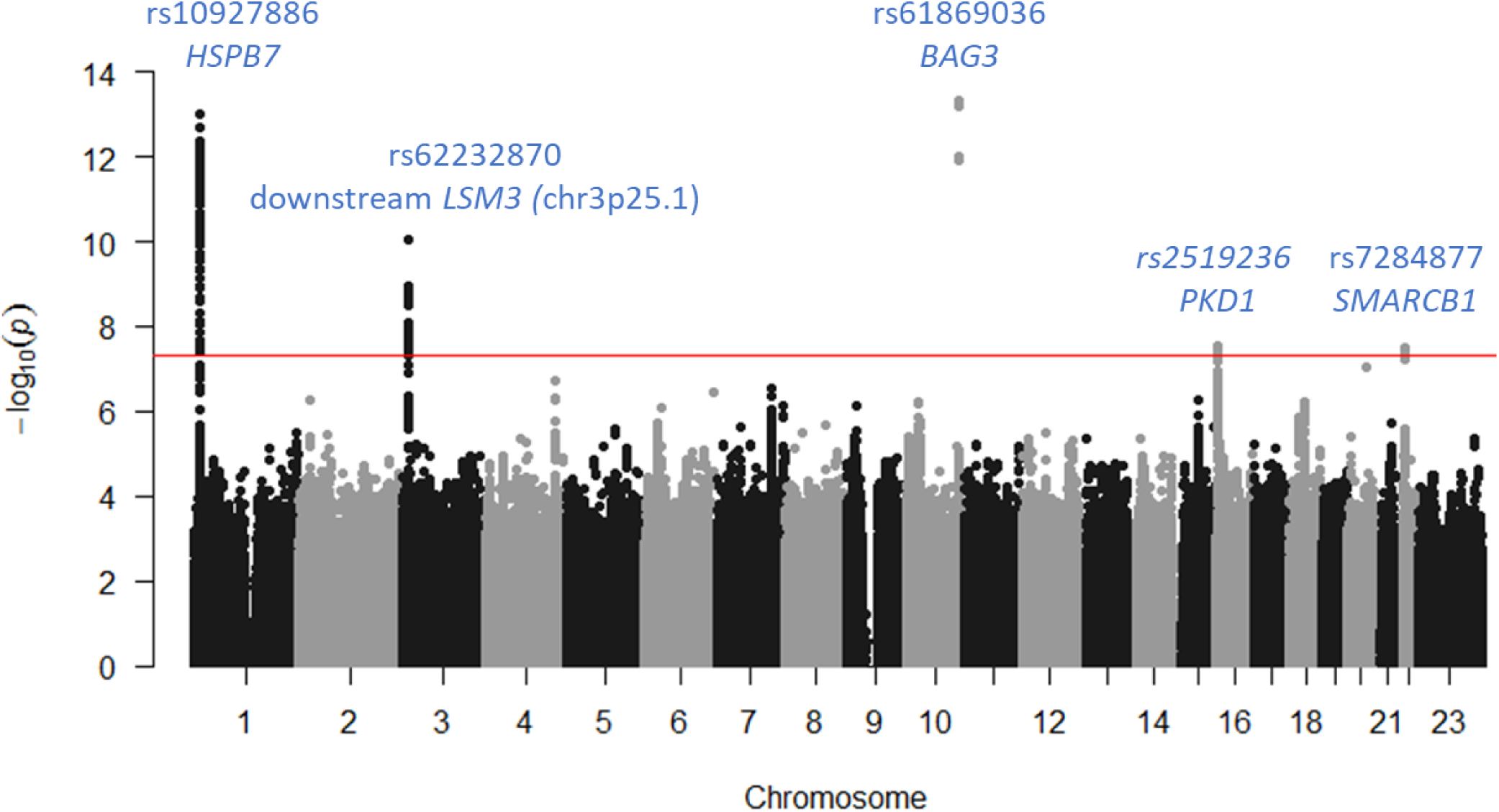
Manhattan plot summarizing the DCM GWAS results.

Conditional GWAS adjusted for the 5 lead SNPs did not reveal any new genome-wide association signal (**Supplementary Figure 7 and 8**).

At chr3p25 locus, a second SNP, rs4684185, showed a strong statistical association (p = 8.4 10^−9^) and is in negative LD (r^2^ = 0.12, D’ = −0.95) with the lead rs62232870. While the rs62232870-A allele with frequency ∼0.21 was associated with an increased OR for DCM of 1.36 [1.24 - 1.49], the common rs4684185-C allele with frequency ∼0.70 was associated with an OR of 1.28 [1.17 – 1.40]. After adjusting on rs62232870 lead SNP, the association of rs4684185 is no longer significant but a residual signal remained (p = 5.10^−4^) suggesting a more complex interaction. A detailed haplotype analysis of these SNPs (**Supplementary Table 2**) showed that, compared to the most frequent rs62232870-G/rs4684185-C haplotype, the rs62232870- A/rs4684185-C haplotype was associated with an increased risk of DCM (OR = 1.22 [1.11 – 1.33]) while its yin-yang haplotype defined by the rs6223870-G/rs4684185-T alleles was protective against DCM (OR = 0.82 [0.76 – 0.89]).

We sought to replicate the observed associations at chromosome 3, 16 and 22 loci in two independent studies totaling 584 DCM patients and 966 controls. The *PKD1* rs148248535 did not show any statistical evidence for replication (p = 0.11) but we confirmed the associations observed at chr3p25.1(p = 7.70 10^−4^ and p = 6.10^−3^ for rs6223870 and rs4684185, respectively), including the yin-yang haplotype association (**Supplementary Table 2**), and at chr22q11.23 (p = 1.40 10^−3^ for rs7284877) (**Table 1)**.

In a combined meta-analysis of the discovery and replication findings, the resulting ORs for DCM were 1.36 [1.25 - 1.48] (p = 5.3 10^−13^) and 1.27 [1.18 - 1.37] (p = 4.8 10^−10^) for chr3p25.1 rs6223870 and rs4684185, respectively and 1.33 [1.22 - 1.46] (p = 5.0 10^−10^) for chr22q11.23 *SMARCB1* rs7284877, with no evidence for heterogeneity across studies (**Table 1**).

### Genetic risk score analysis

We built both an unweighted and a weighted genetic risk score (GRS) using the 4 lead SNPs at the two already known loci (*BAG3* and *HSPB7*) and those at the two replication loci, chr3p25.1 (rs62232870) and chr22q11.23 (rs7284877).

GRS findings are summarized in **Figure 2A and B** and **Table 2A and 2B**. Briefly, the risk of DCM in the discovery cohort for the unweighted GRS (**Table 2A**) was increased by 27% for the subjects with 8 risk alleles (1.27 [1.14-1.42]) and decreased by 21% for those having only one risk allele (0.79 [0.66-0.95]) as compared with the 5 risk alleles reference group (**Figure 2A**). Similar results are found for the weighted GRS (30% increase (1.30 [1.16-1.45]) and 19% decreased (0.81 [0.67-0.97] for those having the lowest and highest score as compared with the 1.6 score reference group (**Figure 2B**). An OR increase with each risk allele increment is still visible, but not significant, in the replication cohorts (**Supplementary Table 3**). To improve size homogeneity between the groups, a quintile repartition was also realized (**Supplementary Figure 9**).

**Table 2A:**
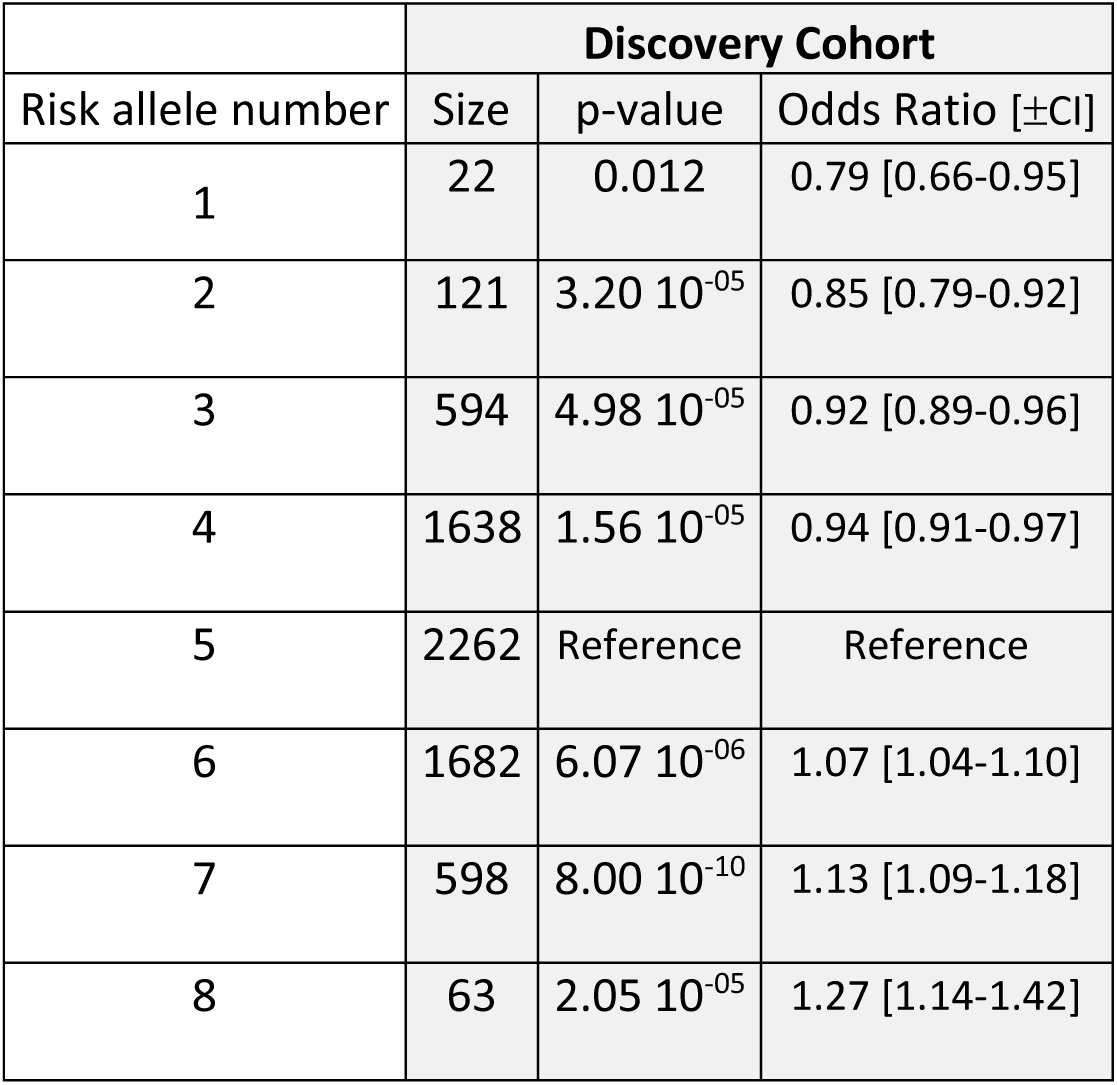
Unweighted Genetic Score for the 6,980 individuals of the discovery cohort and associated OR taking the score 5 as reference.

**Table 2B:**
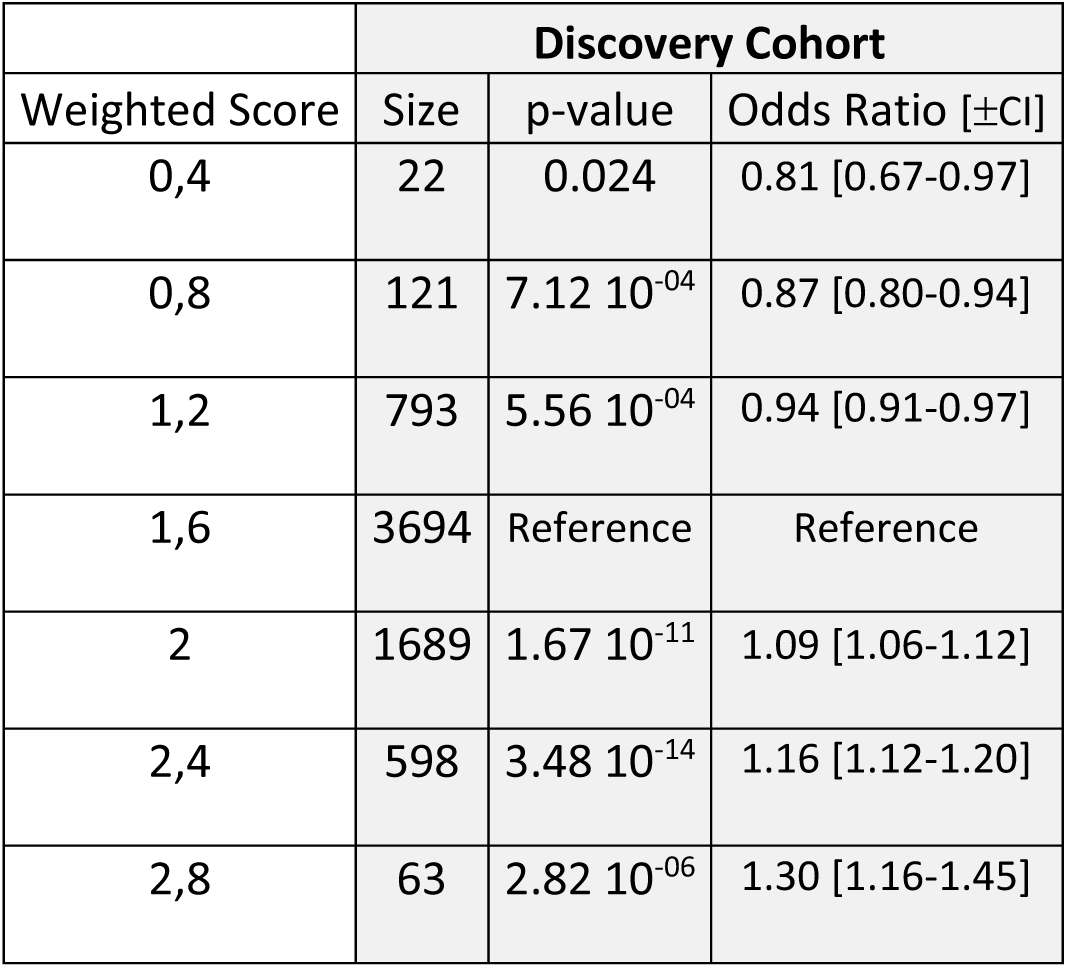
Weighted Genetic Score for the 6,980 individuals of the discovery cohort and associated OR taking the score 1.6 as reference. Weighing was realized taking the sub-meta-analysis (two replication cohorts) beta value

**Figure2A.**
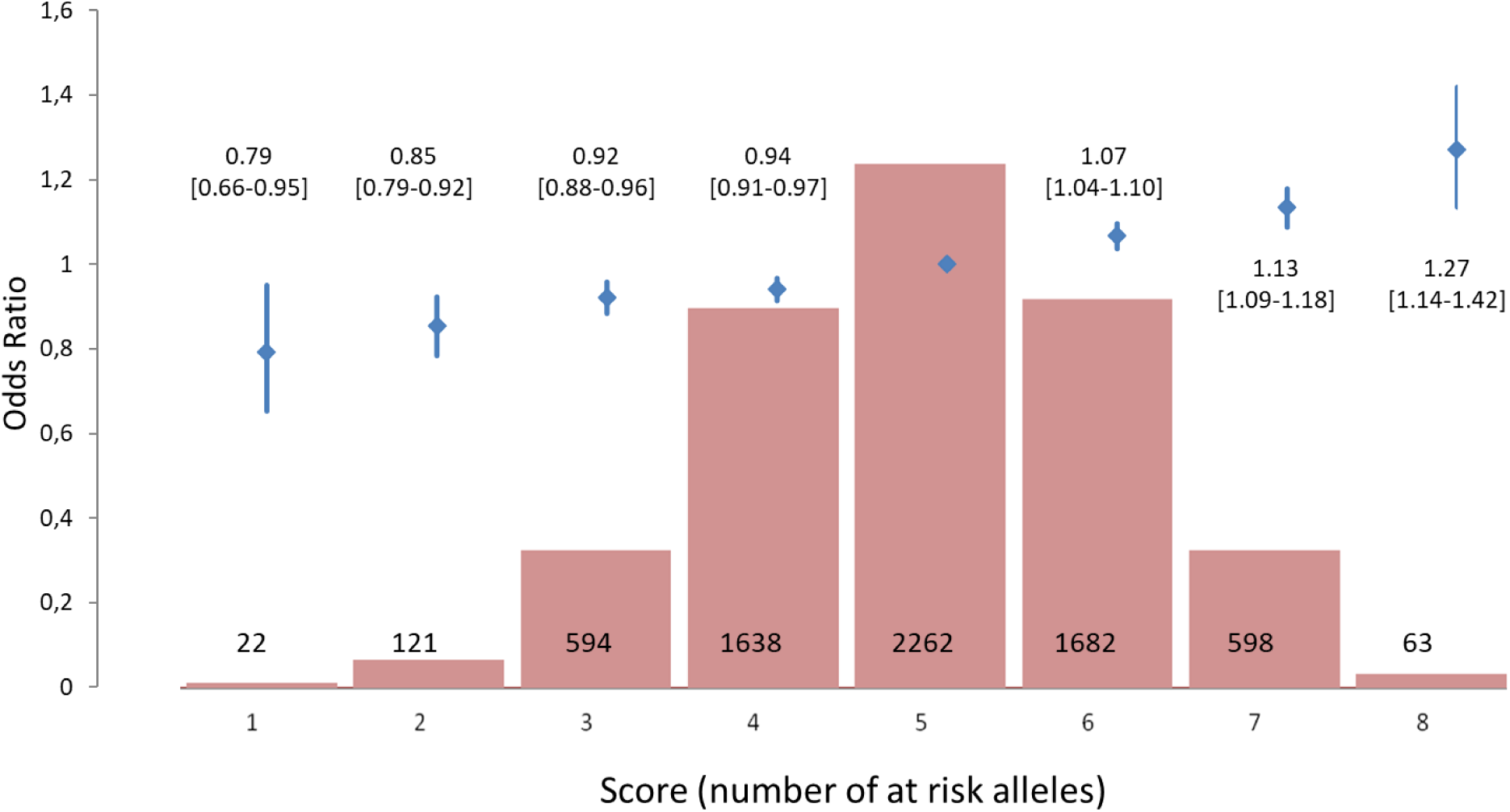
Unweighted Genetic Risk Score for the 6,980 individuals of the discovery cohort and associated OR taking score 5 (presence of 5 risk alleles) as reference

**Figure2B.**
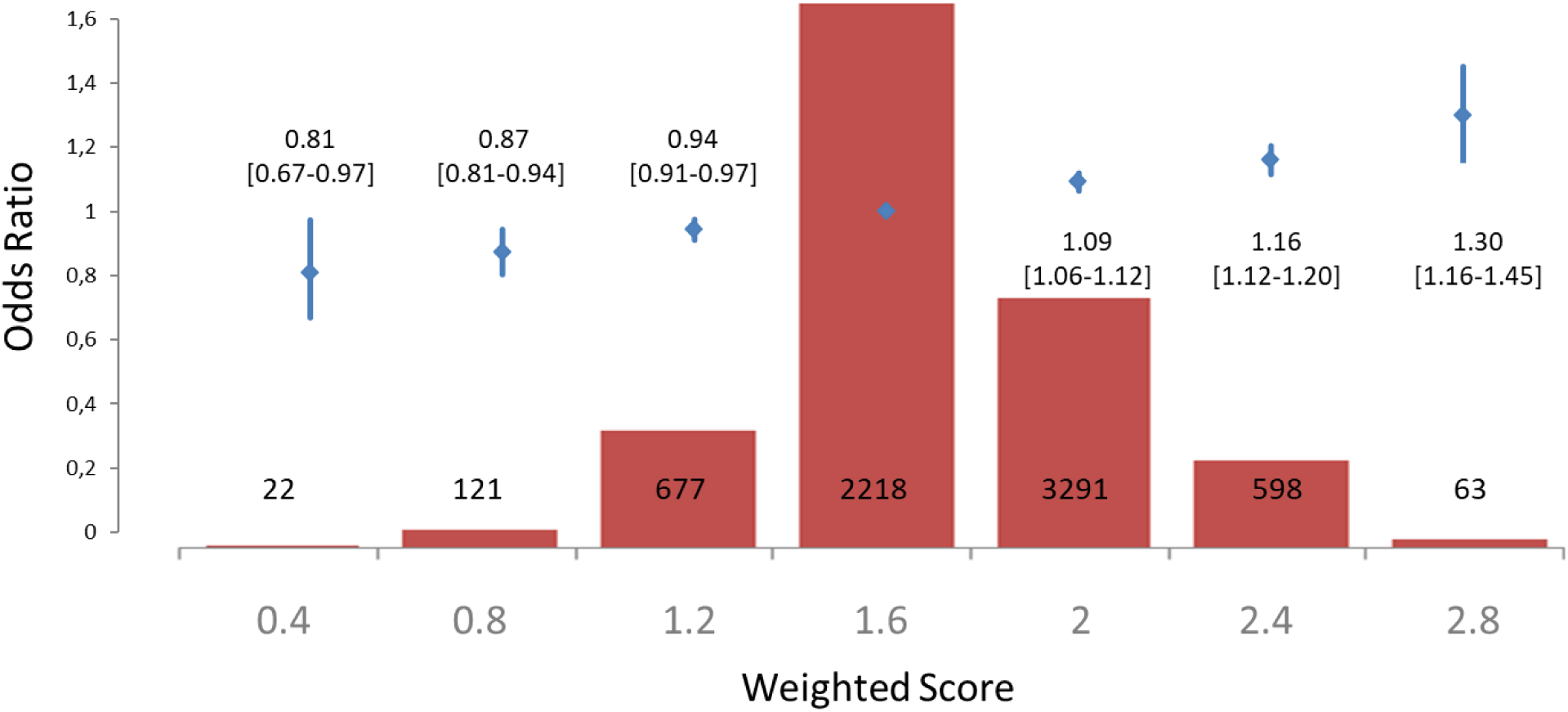
Weighted^*^ Genetic Risk Score for the 6,980 individuals of the discovery cohort and associated OR taking the score 1.6 as reference. ^*^Score of each SNP weighted by the beta value of this SNP in the sub meta-analysis of the two replication cohorts

### Heritability

Using our GWAS summary statistics, the estimated genome-wide DCM heritability in our European populations was around 31% ±8.4. We also examined the genetic correlation between DCM and various cardiovascular traits (e.g. coronary artery disease, heart diseases) but did not observe much shared genetic heritability. The strongest genetic correlation was observed with lipid- and obesity-related traits, e.g. waist circumference (ρ = 0.47, p = 3.2 10^−8^), whole-body mass fat (ρ =0.46, p = 6 10^−8^), or adiponectin (ρ =0.38, p = 7 10^−6^).

### Candidate culprit gene selection strategy at chr3p25.1

As shown in **Figure 3A**, the top SNP, rs62232870, is located at the edge of an active enhancer region lying distal to *LSM3* as evidenced by H3K27ac and H3K4me3 histone marks from ENCODE human LV samples (GSM910575, GSM910580, GSM908951). Of note, this enhancer region is absent from the seven default ENCODE non-cardiomyocyte cell lines suggesting cardiac tissue-specific expression. Other remarkable features support this region as regulatory active, such as significant sequence conservation in a subset of vertebrates, predicted regulatory elements from ORegAnno, hypersensitive sites for DNAseI, and multiple transcription factor (TF) chromatin immunoprecipitation sequencing (ChIP-seq).

**Figure 3.**
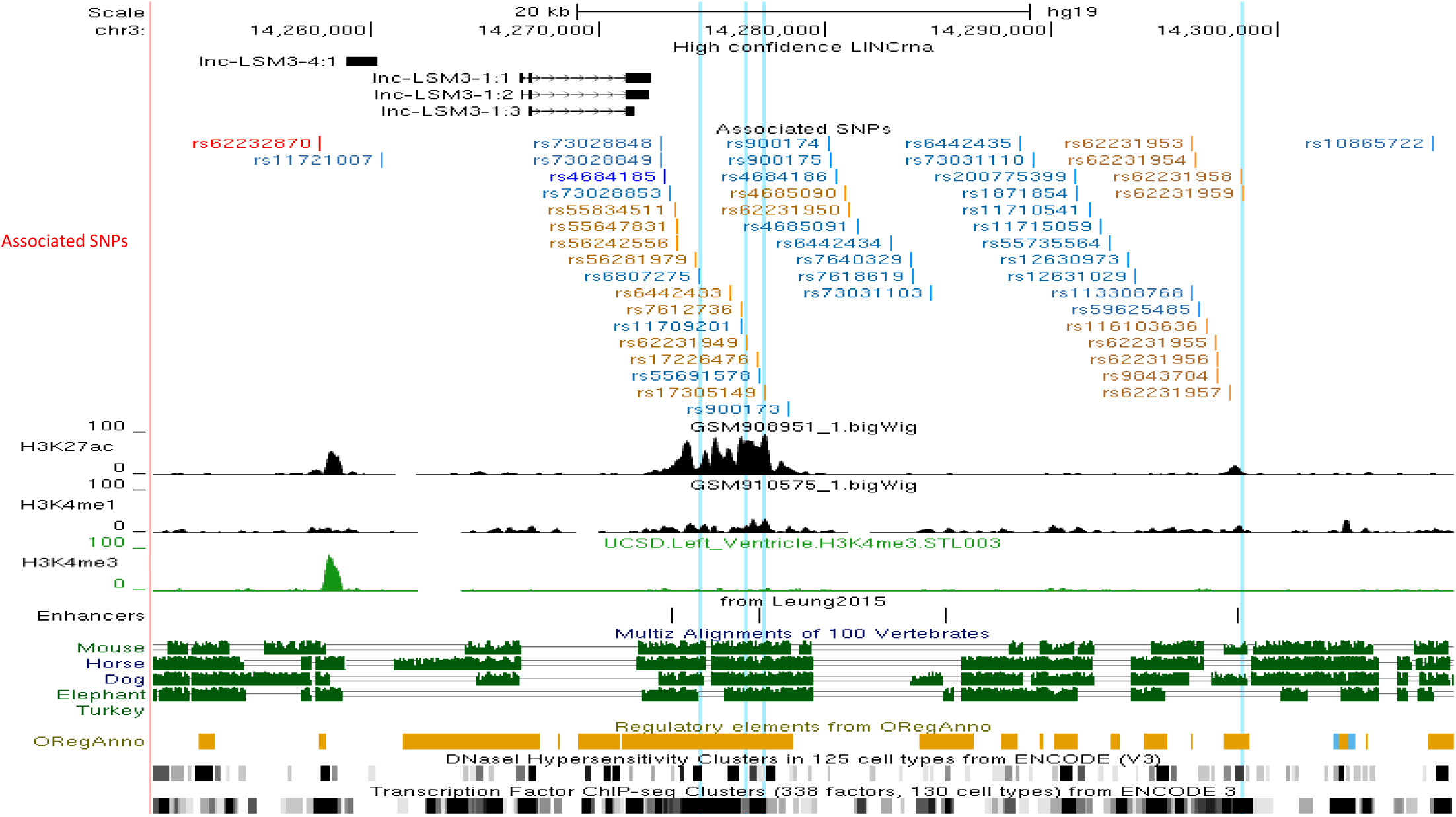
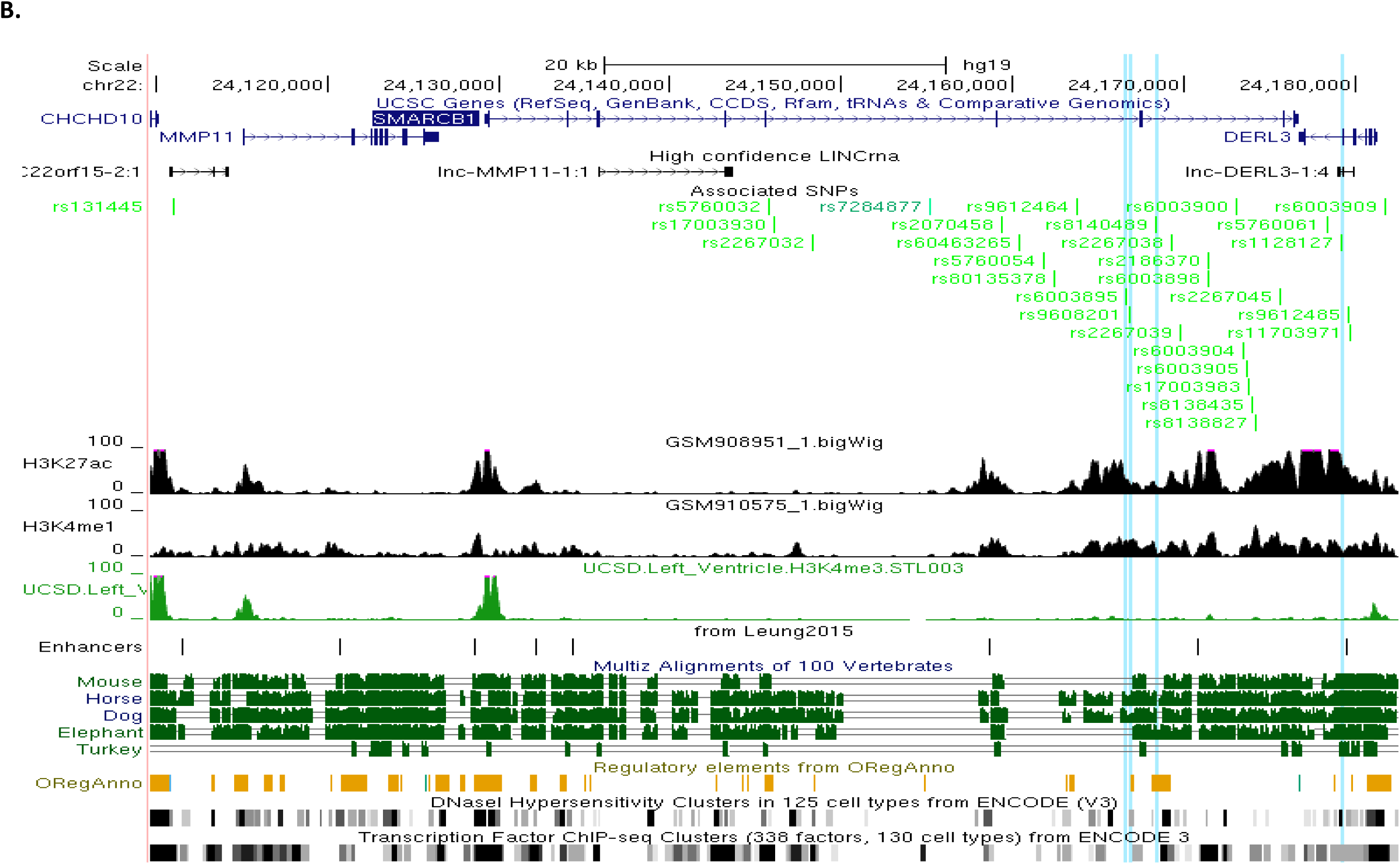
Maps of regulatory DNA features of chromosome 3p25.1 (A) and 22q11.23 (B) associated regions. The associated region is located in a gene desert with long non-coding RNA (lncRNA) sequences for chr3p25.1 [chr3:14,257,356-14,307,016] but covered *MMP11, SMARCB1, DERL3* and lncRNA at chr22q11.23 [chr22:24,110,180-24,182,174]. All SNPs with association p-value < 5 10^−8^ and/or in LD (r^2^ ≥ 0.7) with the lead SNPs (rs62232870 in red; rs4684185 in dark blue; rs7284877 in dark green) are indicated. The SNPs in LD with the lead SNPs are colored orange, light blue and light green, respectively. Features associated with regulatory sequence elements are aligned under the SNPs track (Associated SNPs track) and show that the loci contain enhancer signature according to H3K27ac, H3K4me1 and H3K4me3 ChIP-seq signals prediction for left ventricle enhancers by Leung et al^25^, positive OregAnno regulatory element score^26^, conservation between vertebrates species^27^, DNaseI hypersensitivity and ChiP-seq signal for chromatin interacting proteins linked to transcription activity. Vertical blue lines highlight SNPs with Regulome (http://www.regulomedb.org/)^33^ prediction score below 4 indicating significant potential for being regulatory variants (see **Table 4** for more details)

The associated block of SNPs covers ∼50kb [chr3:14,257,356-14,307,016] overlapping with another partially independent block of SNPs (in LD with the second lead SNP rs4684185, r^2^>0.87) (**Supplementary Figure 3** and **Supplementary Table 4**) where predicted LV H3K27ac and H3K4me1 enhancer marks, reported by Leung *et al.*(28), are present. It is located in a clearly delimited TAD spanning [chr3:14,160,000-14,680,000] (**Figure 4Ab**) which encompasses 6 genes (*CHCHD4, TMEM43, XPC, LSM3, SLC6A6* and *GRIP2*) (**Supplementary Table 5**) whose promoters interact with H3K27ac/H3K4me1 enhancer marks in iPSC-CM (**Figure 4Af**). Interestingly, promoter Hi-C data analysis revealed specific interactions of *SLC6A6* and *GRIP2* promoters with the chr3p25.1 lead SNP bait.

**Table 3:** Lead SNPs (or proxies) eQTls effect, according to GTEx portal, on candidate genes at chromosome 3p25.1 and 22q11.23 loci. ^*^ Lead SNP at chr3p25 .1 (rs62232870) and chr22q11.23 (rs7284877) or proxies whose r^2^ with the lead, >0.6, is given into brackets

| rsID* | eSNP localization | Chr | Regulated Gene | Chr | eSNP p-value | Tissue | p-value GWAS |
| --- | --- | --- | --- | --- | --- | --- | --- |
| rs7612736 ($r^2$: 0.71) | Downstream *LSM3* | 3 | ***SLC6A6*** | 3 | $5.8 \cdot 10^{-05}$ | Heart Atrial Appendage | $1.6 \cdot 10^{-09}$ |
| rs17305149 ($r^2$: 0.71) | Downstream *LSM3* | 3 | ***SLC6A6*** | 3 | $5.8 \cdot 10^{-05}$ | Heart Atrial Appendage | $2.6 \cdot 10^{-09}$ |
| rs62231950 ($r^2$: 0.71) | Downstream *LSM3* | 3 | ***SLC6A6*** | 3 | $5.5 \cdot 10^{-05}$ | Heart Atrial Appendage | $3.1 \cdot 10^{-09}$ |
| rs4685090 ($r^2$: 0.71) | Downstream *LSM3* | 3 | ***SLC6A6*** | 3 | $3.4 \cdot 10^{-05}$ | Heart Atrial Appendage | $2.4 \cdot 10^{-09}$ |
| rs113608926 ($r^2$: 0.69) | Downstream *LSM3* | 3 | ***SLC6A6*** | 3 | $8.5 \cdot 10^{-05}$ | Heart Atrial Appendage | $3.1 \cdot 10^{-08}$ |
| rs62231954 ($r^2$: 0.69) | Downstream *LSM3* | 3 | ***SLC6A6*** | 3 | $8.5 \cdot 10^{-05}$ | Heart Atrial Appendage | $3.4 \cdot 10^{-08}$ |
| rs62231956 ($r^2$: 0.69) | Downstream *LSM3* | 3 | ***SLC6A6*** | 3 | $8.5 \cdot 10^{-05}$ | Heart Atrial Appendage | $3.7 \cdot 10^{-08}$ |
| rs62231957 ($r^2$: 0.69) | Downstream *LSM3* | 3 | ***SLC6A6*** | 3 | $1.9 \cdot 10^{-05}$ | Heart Atrial Appendage | $3.8 \cdot 10^{-08}$ |
| rs62231958 ($r^2$: 0.69) | Downstream *LSM3* | 3 | ***SLC6A6*** | 3 | $4.9 \cdot 10^{-05}$ | Heart Atrial Appendage | $4.5 \cdot 10^{-08}$ |
| rs62231959 ($r^2$: 0.69) | Downstream *LSM3* | 3 | ***SLC6A6*** | 3 | $4.9 \cdot 10^{-05}$ | Heart Atrial Appendage | $4.5 \cdot 10^{-08}$ |
| rs55824951 ($r^2$: 0.69) | Downstream *LSM3* | 3 | ***SLC6A6*** | 3 | $4.9 \cdot 10^{-05}$ | Heart Atrial Appendage | $4.8 \cdot 10^{-08}$ |
| rs7284877 | *SMARCB1* | 22 | ***SMARCB1*** | 22 | $1.5 \cdot 10^{-13}$ | Heart Atrial Appendage | $3.3 \cdot 10^{-08}$ |
| rs7284877 | *SMARCB1* | 22 | ***SMARCB1*** | 22 | $2.8 \cdot 10^{-12}$ | Heart Left Ventricle | $3.3 \cdot 10^{-08}$ |
| rs7284877 | *SMARCB1* | 22 | ***MMP11*** | 22 | $2.8 \cdot 10^{-13}$ | Heart Left Ventricle | $3.3 \cdot 10^{-08}$ |
| rs7284877 | *SMARCB1* | 22 | ***MMP11*** | 22 | $4.3 \cdot 10^{-8}$ | Heart Atrial Appendage | $3.3 \cdot 10^{-08}$ |
| rs7284877 | *SMARCB1* | 22 | ***SMARCB1*** | 22 | $4.9 \cdot 10^{-13}$ | Skeletal muscle | $3.3 \cdot 10^{-08}$ |
| rs7284877 | *SMARCB1* | 22 | ***VPREB3*** | 22 | $1.4 \cdot 10^{-12}$ | Skeletal muscle | $3.3 \cdot 10^{-08}$ |
| rs7284877 | *SMARCB1* | 22 | ***DERL3*** | 22 | $3.5 \cdot 10^{-08}$ | Skeletal muscle | $3.3 \cdot 10^{-08}$ |
| rs9608201 ($r^2$: 1) | *SMARCB1* | 22 | ***MMP11*** | 22 | $2.5 \cdot 10^{-24}$ | Heart Left Ventricle | $3.8 \cdot 10^{-08}$ |
| rs9608201 ($r^2$: 1) | *SMARCB1* | 22 | ***MMP11*** | 22 | $3.3 \cdot 10^{-16}$ | Heart Atrial Appendage | $3.8 \cdot 10^{-08}$ |
| rs9608201 ($r^2$: 1) | *SMARCB1* | 22 | ***SMARCB1*** | 22 | $7.3 \cdot 10^{-22}$ | Heart Left Ventricle | $3.8 \cdot 10^{-08}$ |
| rs9608201 ($r^2$: 1) | *SMARCB1* | 22 | ***SMARCB1*** | 22 | $3.3 \cdot 10^{-16}$ | Heart Atrial Appendage | $3.8 \cdot 10^{-08}$ |
| rs9608201 ($r^2$: 1) | *SMARCB1* | 22 | ***SMARCB1*** | 22 | $3.0 \cdot 10^{-18}$ | Skeletal muscle | $3.8 \cdot 10^{-08}$ |
| rs9608201 (r<sup>2</sup>: 1) | SMARCB1 | 22 | **VPREB3** | 22 | 1.9 10<sup>-8</sup> | Heart Left Ventricle | 3.8 10<sup>-08</sup> |
| rs9608201 (r<sup>2</sup>: 1) | SMARCB1 | 22 | **VPREB3** | 22 | 3.7 10<sup>-18</sup> | Skeletal muscle | 3.8 10<sup>-08</sup> |
| rs9608201 (r<sup>2</sup>: 1) | SMARCB1 | 22 | **DERL3** | 22 | 1.2 10<sup>-6</sup> | Skeletal muscle | 3.8 10<sup>-08</sup> |
| rs2186370 (r<sup>2</sup>: 1) | SMARCB1 | 22 | **MMP11** | 22 | 2.5 10<sup>-24</sup> | Heart Left Ventricle | 3.8 10<sup>-08</sup> |
| rs2186370 (r<sup>2</sup>: 1) | SMARCB1 | 22 | **MMP11** | 22 | 3.3 10<sup>-16</sup> | Heart Atrial Appendage | 3.8 10<sup>-08</sup> |
| rs2186370 (r<sup>2</sup>: 1) | SMARCB1 | 22 | **SMARCB1** | 22 | 7.3 10<sup>-22</sup> | Heart Left Ventricle | 3.8 10<sup>-08</sup> |
| rs2186370 (r<sup>2</sup>: 1) | SMARCB1 | 22 | **SMARCB1** | 22 | 7.5 10<sup>-17</sup> | Heart Atrial Appendage | 3.8 10<sup>-08</sup> |
| rs2186370 (r<sup>2</sup>: 1) | SMARCB1 | 22 | **SMARCB1** | 22 | 3.0 10<sup>-18</sup> | Skeletal muscle | 3.8 10<sup>-08</sup> |
| rs2186370 (r<sup>2</sup>: 1) | SMARCB1 | 22 | **VPREB3** | 22 | 1.9 10<sup>-8</sup> | Heart Left Ventricle | 3.8 10<sup>-08</sup> |
| rs2186370 (r<sup>2</sup>: 1) | SMARCB1 | 22 | **VPREB3** | 22 | 3.7 10<sup>-18</sup> | Skeletal muscle | 3.8 10<sup>-08</sup> |
| rs2186370 (r<sup>2</sup>: 1) | SMARCB1 | 22 | **DERL3** | 22 | 8.1 10<sup>-5</sup> | Skeletal muscle | 3.8 10<sup>-08</sup> |
| rs5760054 (r<sup>2</sup>: 1) | SMARCB1 | 22 | **MMP11** | 22 | 2.5 10<sup>-24</sup> | Heart Left Ventricle | 3.9 10<sup>-08</sup> |
| rs5760054 (r<sup>2</sup>: 1) | SMARCB1 | 22 | **MMP11** | 22 | 3.3 10<sup>-16</sup> | Heart Atrial Appendage | 3.9 10<sup>-08</sup> |
| rs5760054 (r<sup>2</sup>: 1) | SMARCB1 | 22 | **SMARCB1** | 22 | 7.3 10<sup>-22</sup> | Heart Left Ventricle | 3.9 10<sup>-08</sup> |
| rs5760054 (r<sup>2</sup>: 1) | SMARCB1 | 22 | **SMARCB1** | 22 | 7.5 10<sup>-17</sup> | Heart Atrial Appendage | 3.9 10<sup>-08</sup> |
| rs5760054 (r<sup>2</sup>: 1) | SMARCB1 | 22 | **SMARCB1** | 22 | 3.0 10<sup>-18</sup> | Skeletal muscle | 3.9 10<sup>-08</sup> |
| rs5760054 (r<sup>2</sup>: 1) | SMARCB1 | 22 | **VPREB3** | 22 | 1.9 10<sup>-8</sup> | Heart Left Ventricle | 3.9 10<sup>-08</sup> |
| rs5760054 (r<sup>2</sup>: 1) | SMARCB1 | 22 | **VPREB3** | 22 | 3.7 10<sup>-18</sup> | Skeletal muscle | 3.9 10<sup>-08</sup> |
| rs5760054 (r<sup>2</sup>: 1) | SMARCB1 | 22 | **DERL3** | 22 | 1.2 10<sup>-6</sup> | Skeletal muscle | 3.9 10<sup>-08</sup> |
| rs2070458 (r<sup>2</sup>: 1) | SMARCB1 | 22 | **MMP11** | 22 | 2.5 10<sup>-24</sup> | Heart Left Ventricle | 3.9 10<sup>-08</sup> |
| rs2070458 (r<sup>2</sup>: 1) | SMARCB1 | 22 | **MMP11** | 22 | 3.3 10<sup>-16</sup> | Heart Atrial Appendage | 3.9 10<sup>-08</sup> |
| rs2070458 (r<sup>2</sup>: 1) | SMARCB1 | 22 | **SMARCB1** | 22 | 7.3 10<sup>-22</sup> | Heart Left Ventricle | 3.9 10<sup>-08</sup> |
| rs2070458 (r<sup>2</sup>: 1) | SMARCB1 | 22 | **SMARCB1** | 22 | 7.5 10<sup>-17</sup> | Heart Atrial Appendage | 3.9 10<sup>-08</sup> |
| rs2070458 (r<sup>2</sup>: 1) | SMARCB1 | 22 | **SMARCB1** | 22 | 3.0 10<sup>-18</sup> | Skeletal muscle | 3.9 10<sup>-08</sup> |
| rs2070458 (r<sup>2</sup>: 1) | SMARCB1 | 22 | **VPREB3** | 22 | 1.9 10<sup>-8</sup> | Heart Left Ventricle | 3.9 10<sup>-08</sup> |
| rs2070458 (r<sup>2</sup>: 1) | SMARCB1 | 22 | **VPREB3** | 22 | 3.7 10<sup>-18</sup> | Skeletal muscle | 3.9 10<sup>-08</sup> |
| rs2070458 (r<sup>2</sup>: 1) | SMARCB1 | 22 | **DERL3** | 22 | 1.2 10<sup>-6</sup> | Skeletal muscle | 3.9 10<sup>-08</sup> |
| rs2267032 (r<sup>2</sup>: 1) | SMARCB1 | 22 | **MMP11** | 22 | 4 10<sup>-20</sup> | Heart Left Ventricle | 4.2 10<sup>-08</sup> |
| rs2267032 (r<sup>2</sup>: 1) | SMARCB1 | 22 | **MMP11** | 22 | 5.7 10<sup>-15</sup> | Heart Atrial Appendage | 4.2 10<sup>-08</sup> |
| rs2267032 (r<sup>2</sup>: 1) | SMARCB1 | 22 | **SMARCB1** | 22 | 7.3 10<sup>-20</sup> | Heart Left Ventricle | 4.2 10<sup>-08</sup> |
| rs2267032 (r<sup>2</sup>: 1) | SMARCB1 | 22 | **SMARCB1** | 22 | 8.7 10<sup>-16</sup> | Heart Atrial Appendage | 4.2 10<sup>-08</sup> |
| rs2267032 (r<sup>2</sup>: 1) | SMARCB1 | 22 | **SMARCB1** | 22 | 1.3 10<sup>-16</sup> | Skeletal muscle | 4.2 10<sup>-08</sup> |
| rs2267032 (r<sup>2</sup>: 1) | SMARCB1 | 22 | **VPREB3** | 22 | 2.5 10<sup>-9</sup> | Heart Left Ventricle | 4.2 10<sup>-08</sup> |
| rs2267032 (r<sup>2</sup>: 1) | SMARCB1 | 22 | **VPREB3** | 22 | 1.0 10<sup>-17</sup> | Skeletal muscle | 4.2 10<sup>-08</sup> |
| rs2267032 (r<sup>2</sup>: 1) | SMARCB1 | 22 | **DERL3** | 22 | 6.5 10<sup>-6</sup> | Skeletal muscle | 4.2 10<sup>-08</sup> |
| rs5760032 (r<sup>2</sup>: 1) | SMARCB1 | 22 | **MMP11** | 22 | 4 10<sup>-20</sup> | Heart Left Ventricle | 4.4 10<sup>-08</sup> |
| rs5760032 (r<sup>2</sup>: 1) | SMARCB1 | 22 | **MMP11** | 22 | 5.7 10<sup>-15</sup> | Heart Atrial Appendage | 4.4 10<sup>-08</sup> |
| rs5760032 (r<sup>2</sup>: 1) | SMARCB1 | 22 | **SMARCB1** | 22 | 7.3 10<sup>-20</sup> | Heart Left Ventricle | 4.4 10<sup>-08</sup> |
| rs5760032 (r<sup>2</sup>: 1) | SMARCB1 | 22 | **SMARCB1** | 22 | 8.7 10<sup>-16</sup> | Heart Atrial Appendage | 4.4 10<sup>-08</sup> |
| rs5760032 (r<sup>2</sup>: 1) | SMARCB1 | 22 | **SMARCB1** | 22 | 7.8 10<sup>-17</sup> | Skeletal muscle | 4.4 10<sup>-08</sup> |
| rs5760032 (r<sup>2</sup>: 1) | SMARCB1 | 22 | **VPREB3** | 22 | 2.5 10<sup>-9</sup> | Heart Left Ventricle | 4.4 10<sup>-08</sup> |
| rs5760032 (r<sup>2</sup>: 1) | SMARCB1 | 22 | **VPREB3** | 22 | 1.1 10<sup>-17</sup> | Skeletal muscle | 4.4 10<sup>-08</sup> |
| rs5760032 (r<sup>2</sup>: 1) | SMARCB1 | 22 | **DERL3** | 22 | 6.2 10<sup>-6</sup> | Skeletal muscle | 4.4 10<sup>-08</sup> |
| rs5760061 (r<sup>2</sup>: 1) | SMARCB1 | 22 | **MMP11** | 22 | 2.5 10<sup>-24</sup> | Heart Left Ventricle | 4.6 10<sup>-08</sup> |
| rs5760061 (r<sup>2</sup>: 1) | SMARCB1 | 22 | **MMP11** | 22 | 3.3 10<sup>-16</sup> | Heart Atrial Appendage | 4.6 10<sup>-08</sup> |
| rs5760061 (r<sup>2</sup>: 1) | SMARCB1 | 22 | **SMARCB1** | 22 | 7.3 10<sup>-22</sup> | Heart Left Ventricle | 4.6 10<sup>-08</sup> |
| rs5760061 (r<sup>2</sup>: 1) | SMARCB1 | 22 | **SMARCB1** | 22 | 7.5 10<sup>-17</sup> | Heart Atrial Appendage | 4.6 10<sup>-08</sup> |
| rs5760061 (r<sup>2</sup>: 1) | SMARCB1 | 22 | **SMARCB1** | 22 | 6.1 10<sup>-18</sup> | Skeletal muscle | 4.6 10<sup>-08</sup> |
| rs5760061 (r<sup>2</sup>: 1) | SMARCB1 | 22 | **VPREB3** | 22 | 1.9 10<sup>-8</sup> | Heart Left Ventricle | 4.6 10<sup>-08</sup> |
| rs5760061 (r<sup>2</sup>: 1) | SMARCB1 | 22 | **VPREB3** | 22 | 1.1 10<sup>-17</sup> | Skeletal muscle | 4.6 10<sup>-08</sup> |
| rs5760061 (r<sup>2</sup>: 1) | SMARCB1 | 22 | **DERL3** | 22 | 1.1 10<sup>-4</sup> | Skeletal muscle | 4.6 10<sup>-08</sup> |
| rs2267039 (r<sup>2</sup>: 1) | SMARCB1 | 22 | **MMP11** | 22 | 2.5 10<sup>-24</sup> | Heart Left Ventricle | 3.9 10<sup>-08</sup> |
| rs2267039 (r<sup>2</sup>: 1) | SMARCB1 | 22 | **MMP11** | 22 | 3.3 10<sup>-16</sup> | Heart Atrial Appendage | 3.9 10<sup>-08</sup> |
| rs2267039 (r<sup>2</sup>: 1) | SMARCB1 | 22 | **SMARCB1** | 22 | 7.3 10<sup>-22</sup> | Heart Left Ventricle | 3.9 10<sup>-08</sup> |
| rs2267039 (r<sup>2</sup>: 1) | SMARCB1 | 22 | **SMARCB1** | 22 | 7.5 10<sup>-17</sup> | Heart Atrial Appendage | 3.9 10<sup>-08</sup> |
| rs2267039 (r<sup>2</sup>: 1) | SMARCB1 | 22 | **SMARCB1** | 22 | 3.0 10<sup>-18</sup> | Skeletal muscle | 3.9 10<sup>-08</sup> |
| rs2267039 (r<sup>2</sup>: 1) | SMARCB1 | 22 | **VPREB3** | 22 | 1.9 10<sup>-8</sup> | Heart Left Ventricle | 3.9 10<sup>-08</sup> |
| rs2267039 (r<sup>2</sup>: 1) | SMARCB1 | 22 | **VPREB3** | 22 | 3.7 10<sup>-18</sup> | Skeletal muscle | 3.9 10<sup>-08</sup> |
| rs2267039 (r<sup>2</sup>: 1) | SMARCB1 | 22 | **DERL3** | 22 | 1.2 10<sup>-6</sup> | Skeletal muscle | 3.9 10<sup>-08</sup> |
| rs2267038 (r<sup>2</sup>: 0.92) | SMARCB1 | 22 | **MMP11** | 22 | 2.6 10<sup>-21</sup> | Heart Left Ventricle | 3.7 10<sup>-08</sup> |
| rs2267038 (r<sup>2</sup>: 0.92) | SMARCB1 | 22 | **MMP11** | 22 | 5.1 10<sup>-14</sup> | Heart Atrial Appendage | 3.7 10<sup>-08</sup> |
| rs2267038 (r<sup>2</sup>: 0.92) | SMARCB1 | 22 | **SMARCB1** | 22 | 2.1 10<sup>-24</sup> | Heart Left Ventricle | 3.7 10<sup>-08</sup> |
| rs2267038 (r<sup>2</sup>: 0.92) | SMARCB1 | 22 | **SMARCB1** | 22 | 2.0 10<sup>-20</sup> | Heart Atrial Appendage | 3.7 10<sup>-08</sup> |
| rs2267038 (r<sup>2</sup>: 0.92) | SMARCB1 | 22 | **SMARCB1** | 22 | 4.2 10<sup>-23</sup> | Skeletal muscle | 3.7 10<sup>-08</sup> |
| rs2267038 (r<sup>2</sup>: 0.92) | SMARCB1 | 22 | **VPREB3** | 22 | 8.9 10<sup>-9</sup> | Heart Left Ventricle | 3.7 10<sup>-08</sup> |
| rs2267038 (r<sup>2</sup>: 0.92) | SMARCB1 | 22 | **VPREB3** | 22 | 3.2 10<sup>-6</sup> | Heart Atrial Appendage | 3.7 10<sup>-08</sup> |
| rs2267038 (r<sup>2</sup>: 0.92) | SMARCB1 | 22 | **VPREB3** | 22 | 1.8 10<sup>-20</sup> | Skeletal muscle | 3.7 10<sup>-08</sup> |
| rs2267038 (r<sup>2</sup>: 0.92) | SMARCB1 | 22 | **DERL3** | 22 | 5.4 10<sup>-6</sup> | Skeletal muscle | 3.7 10<sup>-08</sup> |
| rs6003909 (r<sup>2</sup>: 0.76) | DERL3 | 22 | **MMP11** | 22 | 2.5 10<sup>-14</sup> | Heart Left Ventricle | 6.3 10<sup>-08</sup> |
| rs6003909 (r<sup>2</sup>: 0.76) | DERL3 | 22 | **MMP11** | 22 | 5.0 10<sup>-10</sup> | Heart Atrial Appendage | 6.3 10<sup>-08</sup> |
| rs6003909 (r<sup>2</sup>: 0.76) | DERL3 | 22 | **SMARCB1** | 22 | 3.5 10<sup>-14</sup> | Heart Left Ventricle | 6.3 10<sup>-08</sup> |
| rs6003909 (r<sup>2</sup>: 0.76) | DERL3 | 22 | **SMARCB1** | 22 | 5.4 10<sup>-17</sup> | Heart Atrial Appendage | 6.3 10<sup>-08</sup> |
| rs6003909 (r<sup>2</sup>: 0.76) | DERL3 | 22 | **SMARCB1** | 22 | 1.1 10<sup>-18</sup> | Skeletal muscle | 6.3 10<sup>-08</sup> |
| rs6003909 (r<sup>2</sup>: 0.76) | DERL3 | 22 | **VPREB3** | 22 | 3.0 10<sup>-7</sup> | Heart Left Ventricle | 6.3 10<sup>-08</sup> |
| rs6003909 (r<sup>2</sup>: 0.76) | DERL3 | 22 | **VPREB3** | 22 | 4.5 10<sup>-13</sup> | Skeletal muscle | 6.3 10<sup>-08</sup> |
| rs6003909 (r<sup>2</sup>: 0.76) | DERL3 | 22 | **DERL3** | 22 | 1.4 10<sup>-8</sup> | Skeletal muscle | 6.3 10<sup>-08</sup> |
| rs80135378 (r<sup>2</sup>: 0.73) | SMARCB1 | 22 | **MMP11** | 22 | 5.6 10<sup>-21</sup> | Heart Left Ventricle | 1.8 10<sup>-05</sup> |
| rs80135378 (r<sup>2</sup>: 0.73) | SMARCB1 | 22 | **MMP11** | 22 | 1.1 10<sup>-14</sup> | Heart Atrial Appendage | 1.8 10<sup>-05</sup> |
| rs80135378 (r<sup>2</sup>: 0.73) | SMARCB1 | 22 | **SMARCB1** | 22 | 2.2 10<sup>-25</sup> | Heart Left Ventricle | 1.8 10<sup>-05</sup> |
| rs80135378 (r<sup>2</sup>: 0.73) | SMARCB1 | 22 | **SMARCB1** | 22 | 5.4 10<sup>-17</sup> | Heart Atrial Appendage | 1.8 10<sup>-05</sup> |
| rs80135378 (r<sup>2</sup>: 0.73) | SMARCB1 | 22 | **SMARCB1** | 22 | 2.0 10<sup>-24</sup> | Skeletal muscle | 1.8 10<sup>-05</sup> |
| rs80135378 (r<sup>2</sup>: 0.73) | SMARCB1 | 22 | **VPREB3** | 22 | 8.3 10<sup>-10</sup> | Heart Left Ventricle | 1.8 10<sup>-05</sup> |
| rs80135378 (r<sup>2</sup>: 0.73) | SMARCB1 | 22 | **VPREB3** | 22 | 4.5 10<sup>-13</sup> | Heart Atrial Appendage | 1.8 10<sup>-05</sup> |
| rs80135378 (r<sup>2</sup>: 0.73) | SMARCB1 | 22 | **VPREB3** | 22 | 3.9 10<sup>-18</sup> | Skeletal muscle | 1.8 10<sup>-05</sup> |
| rs9612464 (r<sup>2</sup>: 0.73) | SMARCB1 | 22 | **MMP11** | 22 | 5.6 10<sup>-21</sup> | Heart Left Ventricle | 1.0 10<sup>-05</sup> |
| rs9612464 (r<sup>2</sup>: 0.73) | SMARCB1 | 22 | **MMP11** | 22 | 1.1 10<sup>-14</sup> | Heart Atrial Appendage | 1.0 10<sup>-05</sup> |
| rs9612464 (r<sup>2</sup>: 0.73) | SMARCB1 | 22 | **SMARCB1** | 22 | 2.2 10<sup>-25</sup> | Heart Left Ventricle | 1.0 10<sup>-05</sup> |
| rs9612464 (r<sup>2</sup>: 0.73) | SMARCB1 | 22 | **SMARCB1** | 22 | 7.7 10<sup>-21</sup> | Heart Atrial Appendage | 1.0 10<sup>-05</sup> |
| rs9612464 (r<sup>2</sup>: 0.73) | SMARCB1 | 22 | **SMARCB1** | 22 | 2.0 10<sup>-24</sup> | Skeletal muscle | 1.0 10<sup>-05</sup> |
| rs9612464 (r<sup>2</sup>: 0.73) | SMARCB1 | 22 | **VPREB3** | 22 | 8.3 10<sup>-10</sup> | Heart Left Ventricle | 1.0 10<sup>-05</sup> |
| rs9612464 (r<sup>2</sup>: 0.73) | SMARCB1 | 22 | **VPREB3** | 22 | 8.4 10<sup>-6</sup> | Heart Atrial Appendage | 1.0 10<sup>-05</sup> |
| rs9612464 (r<sup>2</sup>: 0.73) | SMARCB1 | 22 | **VPREB3** | 22 | 3.9 10<sup>-18</sup> | Skeletal muscle | 1.0 10<sup>-05</sup> |
| rs17003930 (r<sup>2</sup>: 0.73) | SMARCB1 | 22 | **MMP11** | 22 | 1.7 10<sup>-14</sup> | Heart Left Ventricle | 1.9 10<sup>-05</sup> |
| rs17003930 (r<sup>2</sup>: 0.73) | SMARCB1 | 22 | **MMP11** | 22 | 1.3 10<sup>-11</sup> | Heart Atrial Appendage | 1.9 10<sup>-05</sup> |
| rs17003930 (r<sup>2</sup>: 0.73) | SMARCB1 | 22 | **SMARCB1** | 22 | 3.0 10<sup>-16</sup> | Heart Left Ventricle | 1.9 10<sup>-05</sup> |
| rs17003930 (r<sup>2</sup>: 0.73) | SMARCB1 | 22 | **SMARCB1** | 22 | 1.3 10<sup>-16</sup> | Heart Atrial Appendage | 1.9 10<sup>-05</sup> |
| rs17003930 (r<sup>2</sup>: 0.73) | SMARCB1 | 22 | **SMARCB1** | 22 | 1.6 10<sup>-19</sup> | Skeletal muscle | 1.9 10<sup>-05</sup> |
| rs17003930 (r<sup>2</sup>: 0.73) | SMARCB1 | 22 | **VPREB3** | 22 | 3.8 10<sup>-12</sup> | Heart Left Ventricle | 1.9 10<sup>-05</sup> |
| rs17003930 (r<sup>2</sup>: 0.73) | SMARCB1 | 22 | **VPREB3** | 22 | 5.0 10<sup>-6</sup> | Heart Atrial Appendage | 1.9 10<sup>-05</sup> |
| rs17003930 (r<sup>2</sup>: 0.73) | SMARCB1 | 22 | **VPREB3** | 22 | 3.4 10<sup>-17</sup> | Skeletal muscle | 1.9 10<sup>-05</sup> |
| rs17003930 (r<sup>2</sup>: 0.73) | SMARCB1 | 22 | **DERL3** | 22 | 2.8 10<sup>-5</sup> | Skeletal muscle | 1.9 10<sup>-05</sup> |
| rs6003895 (r<sup>2</sup>: 0.73) | SMARCB1 | 22 | **MMP11** | 22 | 5.6 10<sup>-21</sup> | Heart Left Ventricle | 9.6 10<sup>-06</sup> |
| rs6003895 (r<sup>2</sup>: 0.73) | SMARCB1 | 22 | **MMP11** | 22 | 1.1 10<sup>-14</sup> | Heart Atrial Appendage | 9.6 10<sup>-06</sup> |
| rs6003895 (r<sup>2</sup>: 0.73) | SMARCB1 | 22 | **SMARCB1** | 22 | 2.2 10<sup>-25</sup> | Heart Left Ventricle | 9.6 10<sup>-06</sup> |
| rs6003895 (r<sup>2</sup>: 0.73) | SMARCB1 | 22 | **SMARCB1** | 22 | 7.7 10<sup>-21</sup> | Heart Atrial Appendage | 9.6 10<sup>-06</sup> |
| rs6003895 (r<sup>2</sup>: 0.73) | SMARCB1 | 22 | **SMARCB1** | 22 | 2.0 10<sup>-24</sup> | Skeletal muscle | 9.6 10<sup>-06</sup> |
| rs6003895 (r<sup>2</sup>: 0.73) | SMARCB1 | 22 | **VPREB3** | 22 | 8.3 10<sup>-10</sup> | Heart Left Ventricle | 9.6 10<sup>-06</sup> |
| rs6003895 (r<sup>2</sup>: 0.73) | SMARCB1 | 22 | **VPREB3** | 22 | 8.4 10<sup>-6</sup> | Heart Atrial Appendage | 9.6 10<sup>-06</sup> |
| rs6003895 (r<sup>2</sup>: 0.73) | SMARCB1 | 22 | **VPREB3** | 22 | 3.9 10<sup>-18</sup> | Skeletal muscle | 9.6 10<sup>-06</sup> |
| rs8140489 (r<sup>2</sup>: 0.73) | SMARCB1 | 22 | **MMP11** | 22 | 5.6 10<sup>-21</sup> | Heart Left Ventricle | 9.5 10<sup>-06</sup> |
| rs8140489 (r<sup>2</sup>: 0.73) | SMARCB1 | 22 | **MMP11** | 22 | 1.1 10<sup>-14</sup> | Heart Atrial Appendage | 9.5 10<sup>-06</sup> |
| rs8140489 (r<sup>2</sup>: 0.73) | SMARCB1 | 22 | **SMARCB1** | 22 | 2.2 10<sup>-25</sup> | Heart Left Ventricle | 9.5 10<sup>-06</sup> |
| rs8140489 (r<sup>2</sup>: 0.73) | SMARCB1 | 22 | **SMARCB1** | 22 | 7.7 10<sup>-21</sup> | Heart Atrial Appendage | 9.5 10<sup>-06</sup> |
| rs8140489 (r<sup>2</sup>: 0.73) | SMARCB1 | 22 | **SMARCB1** | 22 | 2.0 10<sup>-24</sup> | Skeletal muscle | 9.5 10<sup>-06</sup> |
| rs8140489 (r<sup>2</sup>: 0.73) | SMARCB1 | 22 | **VPREB3** | 22 | 8.3 10<sup>-10</sup> | Heart Left Ventricle | 9.5 10<sup>-06</sup> |
| rs8140489 (r<sup>2</sup>: 0.73) | SMARCB1 | 22 | **VPREB3** | 22 | 8.4 10<sup>-6</sup> | Heart Atrial Appendage | 9.5 10<sup>-06</sup> |
| rs8140489 (r<sup>2</sup>: 0.73) | SMARCB1 | 22 | **VPREB3** | 22 | 3.9 10<sup>-18</sup> | Skeletal muscle | 9.5 10<sup>-06</sup> |
| rs6003898 (r<sup>2</sup>: 0.73) | SMARCB1 | 22 | **MMP11** | 22 | 5.6 10<sup>-21</sup> | Heart Left Ventricle | 9.2 10<sup>-06</sup> |
| rs6003898 (r<sup>2</sup>: 0.73) | SMARCB1 | 22 | **MMP11** | 22 | 1.1 10<sup>-14</sup> | Heart Atrial Appendage | 9.2 10<sup>-06</sup> |
| rs6003898 (r<sup>2</sup>: 0.73) | SMARCB1 | 22 | **SMARCB1** | 22 | 2.2 10<sup>-25</sup> | Heart Left Ventricle | 9.2 10<sup>-06</sup> |
| rs6003898 (r<sup>2</sup>: 0.73) | SMARCB1 | 22 | **SMARCB1** | 22 | 7.7 10<sup>-21</sup> | Heart Atrial Appendage | 9.2 10<sup>-06</sup> |
| rs6003898 (r<sup>2</sup>: 0.73) | SMARCB1 | 22 | **SMARCB1** | 22 | 2.0 10<sup>-24</sup> | Skeletal muscle | 9.2 10<sup>-06</sup> |
| rs6003898 (r<sup>2</sup>: 0.73) | SMARCB1 | 22 | **VPREB3** | 22 | 8.3 10<sup>-10</sup> | Heart Left Ventricle | 9.2 10<sup>-06</sup> |
| rs6003898 (r<sup>2</sup>: 0.73) | SMARCB1 | 22 | **VPREB3** | 22 | 8.4 10<sup>-6</sup> | Heart Atrial Appendage | 9.2 10<sup>-06</sup> |
| rs6003898 (r<sup>2</sup>: 0.73) | SMARCB1 | 22 | **VPREB3** | 22 | 3.9 10<sup>-18</sup> | Skeletal muscle | 9.2 10<sup>-06</sup> |
| rs6003900 (r<sup>2</sup>: 0.73) | SMARCB1 | 22 | **MMP11** | 22 | 5.6 10<sup>-21</sup> | Heart Left Ventricle | 8.8 10<sup>-06</sup> |
| rs6003900 (r<sup>2</sup>: 0.73) | SMARCB1 | 22 | **MMP11** | 22 | 1.1 10<sup>-14</sup> | Heart Atrial Appendage | 8.8 10<sup>-06</sup> |
| rs6003900 (r<sup>2</sup>: 0.73) | SMARCB1 | 22 | **SMARCB1** | 22 | 2.2 10<sup>-25</sup> | Heart Left Ventricle | 8.8 10<sup>-06</sup> |
| rs6003900 (r<sup>2</sup>: 0.73) | SMARCB1 | 22 | **SMARCB1** | 22 | 7.7 10<sup>-21</sup> | Heart Atrial Appendage | 8.8 10<sup>-06</sup> |
| rs6003900 (r<sup>2</sup>: 0.73) | SMARCB1 | 22 | **SMARCB1** | 22 | 9.7 10<sup>-25</sup> | Skeletal muscle | 8.8 10<sup>-06</sup> |
| rs6003900 ($r^2$: 0.73) | SMARCB1 | 22 | **VPREB3** | 22 | $8.3 \cdot 10^{-10}$ | Heart Left Ventricle | $8.8 \cdot 10^{-06}$ |
| rs6003900 ($r^2$: 0.73) | SMARCB1 | 22 | **VPREB3** | 22 | $8.4 \cdot 10^{-6}$ | Heart Atrial Appendage | $8.8 \cdot 10^{-06}$ |
| rs6003900 ($r^2$: 0.73) | SMARCB1 | 22 | **VPREB3** | 22 | $4.3 \cdot 10^{-18}$ | Skeletal muscle | $8.8 \cdot 10^{-06}$ |
| rs6003904 ($r^2$: 0.73) | SMARCB1 | 22 | **MMP11** | 22 | $5.6 \cdot 10^{-21}$ | Heart Left Ventricle | $8.8 \cdot 10^{-06}$ |
| rs6003904 ($r^2$: 0.73) | SMARCB1 | 22 | **MMP11** | 22 | $1.1 \cdot 10^{-14}$ | Heart Atrial Appendage | $8.8 \cdot 10^{-06}$ |
| rs6003904 ($r^2$: 0.73) | SMARCB1 | 22 | **SMARCB1** | 22 | $2.2 \cdot 10^{-25}$ | Heart Left Ventricle | $8.8 \cdot 10^{-06}$ |
| rs6003904 ($r^2$: 0.73) | SMARCB1 | 22 | **SMARCB1** | 22 | $7.7 \cdot 10^{-21}$ | Heart Atrial Appendage | $8.8 \cdot 10^{-06}$ |
| rs6003904 ($r^2$: 0.73) | SMARCB1 | 22 | **SMARCB1** | 22 | $9.7 \cdot 10^{-25}$ | Skeletal muscle | $8.8 \cdot 10^{-06}$ |
| rs6003904 ($r^2$: 0.73) | SMARCB1 | 22 | **VPREB3** | 22 | $8.3 \cdot 10^{-10}$ | Heart Left Ventricle | $8.8 \cdot 10^{-06}$ |
| rs6003904 ($r^2$: 0.73) | SMARCB1 | 22 | **VPREB3** | 22 | $8.4 \cdot 10^{-6}$ | Heart Atrial Appendage | $8.8 \cdot 10^{-06}$ |
| rs6003904 ($r^2$: 0.73) | SMARCB1 | 22 | **VPREB3** | 22 | $4.3 \cdot 10^{-18}$ | Skeletal muscle | $8.8 \cdot 10^{-06}$ |
| rs6003905 ($r^2$: 0.73) | SMARCB1 | 22 | **MMP11** | 22 | $5.6 \cdot 10^{-21}$ | Heart Left Ventricle | $8.1 \cdot 10^{-06}$ |
| rs6003905 ($r^2$: 0.73) | SMARCB1 | 22 | **MMP11** | 22 | $1.1 \cdot 10^{-14}$ | Heart Atrial Appendage | $8.1 \cdot 10^{-06}$ |
| rs6003905 ($r^2$: 0.73) | SMARCB1 | 22 | **SMARCB1** | 22 | $2.2 \cdot 10^{-25}$ | Heart Left Ventricle | $8.1 \cdot 10^{-06}$ |
| rs6003905 ($r^2$: 0.73) | SMARCB1 | 22 | **SMARCB1** | 22 | $7.7 \cdot 10^{-21}$ | Heart Atrial Appendage | $8.1 \cdot 10^{-06}$ |
| rs6003905 ($r^2$: 0.73) | SMARCB1 | 22 | **SMARCB1** | 22 | $9.7 \cdot 10^{-25}$ | Skeletal muscle | $8.1 \cdot 10^{-06}$ |
| rs6003905 ($r^2$: 0.73) | SMARCB1 | 22 | **VPREB3** | 22 | $8.3 \cdot 10^{-10}$ | Heart Left Ventricle | $8.1 \cdot 10^{-06}$ |
| rs6003905 ($r^2$: 0.73) | SMARCB1 | 22 | **VPREB3** | 22 | $8.4 \cdot 10^{-6}$ | Heart Atrial Appendage | $8.1 \cdot 10^{-06}$ |
| rs6003905 ($r^2$: 0.73) | SMARCB1 | 22 | **VPREB3** | 22 | $4.3 \cdot 10^{-18}$ | Skeletal muscle | $8.1 \cdot 10^{-06}$ |
| rs17003983 ($r^2$: 0.73) | SMARCB1 | 22 | **MMP11** | 22 | $1.5 \cdot 10^{-15}$ | Heart Left Ventricle | $7.3 \cdot 10^{-06}$ |
| rs17003983 ($r^2$: 0.73) | SMARCB1 | 22 | **MMP11** | 22 | $1.2 \cdot 10^{-11}$ | Heart Atrial Appendage | $7.3 \cdot 10^{-06}$ |
| rs17003983 ($r^2$: 0.73) | SMARCB1 | 22 | **SMARCB1** | 22 | $1.3 \cdot 10^{-17}$ | Heart Left Ventricle | $7.3 \cdot 10^{-06}$ |
| rs17003983 ($r^2$: 0.73) | SMARCB1 | 22 | **SMARCB1** | 22 | $2.5 \cdot 10^{-16}$ | Heart Atrial Appendage | $7.3 \cdot 10^{-06}$ |
| rs17003983 ($r^2$: 0.73) | SMARCB1 | 22 | **SMARCB1** | 22 | $3.5 \cdot 10^{-23}$ | Skeletal muscle | $7.3 \cdot 10^{-06}$ |
| rs17003983 ($r^2$: 0.73) | SMARCB1 | 22 | **VPREB3** | 22 | $1.3 \cdot 10^{-12}$ | Heart Left Ventricle | $7.3 \cdot 10^{-06}$ |
| rs17003983 ($r^2$: 0.73) | SMARCB1 | 22 | **VPREB3** | 22 | $2.7 \cdot 10^{-6}$ | Heart Atrial Appendage | $7.3 \cdot 10^{-06}$ |
| rs17003983 ($r^2$: 0.73) | SMARCB1 | 22 | **VPREB3** | 22 | $3.5 \cdot 10^{-17}$ | Skeletal muscle | $7.3 \cdot 10^{-06}$ |
| rs17003983 ($r^2$: 0.73) | SMARCB1 | 22 | **DERL3** | 22 | $1.7 \cdot 10^{-6}$ | Skeletal muscle | $7.3 \cdot 10^{-06}$ |
| rs8138435 ($r^2$: 0.73) | SMARCB1 | 22 | **MMP11** | 22 | $5.6 \cdot 10^{-21}$ | Heart Left Ventricle | $8.8 \cdot 10^{-06}$ |
| rs8138435 ($r^2$: 0.73) | SMARCB1 | 22 | **MMP11** | 22 | $1.1 \cdot 10^{-14}$ | Heart Atrial Appendage | $8.8 \cdot 10^{-06}$ |
| rs8138435 ($r^2$: 0.73) | SMARCB1 | 22 | **SMARCB1** | 22 | $2.2 \cdot 10^{-25}$ | Heart Left Ventricle | $8.8 \cdot 10^{-06}$ |
| rs8138435 ($r^2$: 0.73) | SMARCB1 | 22 | **SMARCB1** | 22 | $7.7 \cdot 10^{-21}$ | Heart Atrial Appendage | $8.8 \cdot 10^{-06}$ |
| rs8138435 ($r^2$: 0.73) | SMARCB1 | 22 | **SMARCB1** | 22 | $9.7 \cdot 10^{-25}$ | Skeletal muscle | $8.8 \cdot 10^{-06}$ |
| rs8138435 ($r^2$: 0.73) | SMARCB1 | 22 | **VPREB3** | 22 | $8.3 \cdot 10^{-10}$ | Heart Left Ventricle | $8.8 \cdot 10^{-06}$ |
| rs8138435 ($r^2$: 0.73) | SMARCB1 | 22 | **VPREB3** | 22 | $8.4 \cdot 10^{-6}$ | Heart Atrial Appendage | $8.8 \cdot 10^{-06}$ |
| rs8138435 ($r^2$: 0.73) | SMARCB1 | 22 | **VPREB3** | 22 | $4.3 \cdot 10^{-18}$ | Skeletal muscle | $8.8 \cdot 10^{-06}$ |
| rs8138827 ($r^2$: 0.73) | SMARCB1 | 22 | **MMP11** | 22 | $5.6 \cdot 10^{-21}$ | Heart Left Ventricle | $8.8 \cdot 10^{-06}$ |
| rs8138827 ($r^2$: 0.73) | SMARCB1 | 22 | **MMP11** | 22 | $1.1 \cdot 10^{-14}$ | Heart Atrial Appendage | $8.8 \cdot 10^{-06}$ |
| rs8138827 ($r^2$: 0.73) | SMARCB1 | 22 | **SMARCB1** | 22 | $2.2 \cdot 10^{-25}$ | Heart Left Ventricle | $8.8 \cdot 10^{-06}$ |
| rs8138827 ($r^2$: 0.73) | SMARCB1 | 22 | **SMARCB1** | 22 | $7.7 \cdot 10^{-21}$ | Heart Atrial Appendage | $8.8 \cdot 10^{-06}$ |
| rs8138827 ($r^2$: 0.73) | SMARCB1 | 22 | **SMARCB1** | 22 | $9.7 \cdot 10^{-25}$ | Skeletal muscle | $8.8 \cdot 10^{-06}$ |
| rs8138827 ($r^2$: 0.73) | SMARCB1 | 22 | **VPREB3** | 22 | $8.3 \cdot 10^{-10}$ | Heart Left Ventricle | $8.8 \cdot 10^{-06}$ |
| rs8138827 ($r^2$: 0.73) | SMARCB1 | 22 | **VPREB3** | 22 | $8.4 \cdot 10^{-6}$ | Heart Atrial Appendage | $8.8 \cdot 10^{-06}$ |
| rs8138827 ($r^2$: 0.73) | SMARCB1 | 22 | **VPREB3** | 22 | $4.3 \cdot 10^{-18}$ | Skeletal muscle | $8.8 \cdot 10^{-06}$ |
| rs2267045 ($r^2$: 0.73) | SMARCB1 | 22 | **MMP11** | 22 | $6.1 \cdot 10^{-23}$ | Heart Left Ventricle | $8.5 \cdot 10^{-06}$ |
| rs2267045 ($r^2$: 0.73) | SMARCB1 | 22 | **MMP11** | 22 | $2.4 \cdot 10^{-15}$ | Heart Atrial Appendage | $8.5 \cdot 10^{-06}$ |
| rs2267045 ($r^2$: 0.73) | SMARCB1 | 22 | **SMARCB1** | 22 | $3.2 \cdot 10^{-26}$ | Heart Left Ventricle | $8.5 \cdot 10^{-06}$ |
| rs2267045 ($r^2$: 0.73) | SMARCB1 | 22 | **SMARCB1** | 22 | $4.7 \cdot 10^{-21}$ | Heart Atrial Appendage | $8.5 \cdot 10^{-06}$ |
| rs2267045 ($r^2$: 0.73) | SMARCB1 | 22 | **SMARCB1** | 22 | $1.9 \cdot 10^{-24}$ | Skeletal muscle | $8.5 \cdot 10^{-06}$ |
| rs2267045 ($r^2$: 0.73) | SMARCB1 | 22 | **VPREB3** | 22 | $2.0 \cdot 10^{-9}$ | Heart Left Ventricle | $8.5 \cdot 10^{-06}$ |
| rs2267045 ($r^2$: 0.73) | SMARCB1 | 22 | **VPREB3** | 22 | $2.2 \cdot 10^{-5}$ | Heart Atrial Appendage | $8.5 \cdot 10^{-06}$ |
| rs2267045 ($r^2$: 0.73) | SMARCB1 | 22 | **VPREB3** | 22 | $3.3 \cdot 10^{-18}$ | Skeletal muscle | $8.5 \cdot 10^{-06}$ |
| rs1128127 (r<sup>2</sup>: 0.73) | SMARCB1 | 22 | **MMP11** | 22 | 5.6 10<sup>-21</sup> | Heart Left Ventricle | 1.4 10<sup>-05</sup> |
| rs1128127 (r<sup>2</sup>: 0.73) | SMARCB1 | 22 | **MMP11** | 22 | 1.1 10<sup>-14</sup> | Heart Atrial Appendage | 1.4 10<sup>-05</sup> |
| rs1128127 (r<sup>2</sup>: 0.73) | SMARCB1 | 22 | **SMARCB1** | 22 | 2.2 10<sup>-25</sup> | Heart Left Ventricle | 1.4 10<sup>-05</sup> |
| rs1128127 (r<sup>2</sup>: 0.73) | SMARCB1 | 22 | **SMARCB1** | 22 | 7.7 10<sup>-21</sup> | Heart Atrial Appendage | 1.4 10<sup>-05</sup> |
| rs1128127 (r<sup>2</sup>: 0.73) | SMARCB1 | 22 | **SMARCB1** | 22 | 4.9 10<sup>-24</sup> | Skeletal muscle | 1.4 10<sup>-05</sup> |
| rs1128127 (r<sup>2</sup>: 0.73) | SMARCB1 | 22 | **VPREB3** | 22 | 8.3 10<sup>-10</sup> | Heart Left Ventricle | 1.4 10<sup>-05</sup> |
| rs1128127 (r<sup>2</sup>: 0.73) | SMARCB1 | 22 | **VPREB3** | 22 | 8.4 10<sup>-6</sup> | Heart Atrial Appendage | 1.4 10<sup>-05</sup> |
| rs1128127 (r<sup>2</sup>: 0.73) | SMARCB1 | 22 | **VPREB3** | 22 | 1.2 10<sup>-17</sup> | Skeletal muscle | 1.4 10<sup>-05</sup> |
| rs9612485 (r<sup>2</sup>: 0.73) | SMARCB1 | 22 | **MMP11** | 22 | 5.6 10<sup>-21</sup> | Heart Left Ventricle | 1.5 10<sup>-05</sup> |
| rs9612485 (r<sup>2</sup>: 0.73) | SMARCB1 | 22 | **MMP11** | 22 | 1.1 10<sup>-14</sup> | Heart Atrial Appendage | 1.5 10<sup>-05</sup> |
| rs9612485 (r<sup>2</sup>: 0.73) | SMARCB1 | 22 | **SMARCB1** | 22 | 2.2 10<sup>-25</sup> | Heart Left Ventricle | 1.5 10<sup>-05</sup> |
| rs9612485 (r<sup>2</sup>: 0.73) | SMARCB1 | 22 | **SMARCB1** | 22 | 7.7 10<sup>-21</sup> | Heart Atrial Appendage | 1.5 10<sup>-05</sup> |
| rs9612485 (r<sup>2</sup>: 0.73) | SMARCB1 | 22 | **SMARCB1** | 22 | 4.9 10<sup>-24</sup> | Skeletal muscle | 1.5 10<sup>-05</sup> |
| rs9612485 (r<sup>2</sup>: 0.73) | SMARCB1 | 22 | **VPREB3** | 22 | 8.3 10<sup>-10</sup> | Heart Left Ventricle | 1.5 10<sup>-05</sup> |
| rs9612485 (r<sup>2</sup>: 0.73) | SMARCB1 | 22 | **VPREB3** | 22 | 8.4 10<sup>-6</sup> | Heart Atrial Appendage | 1.5 10<sup>-05</sup> |
| rs9612485 (r<sup>2</sup>: 0.73) | SMARCB1 | 22 | **VPREB3** | 22 | 1.2 10<sup>-17</sup> | Skeletal muscle | 1.5 10<sup>-05</sup> |
| rs11703971 (r<sup>2</sup>: 0.73) | SMARCB1 | 22 | **MMP11** | 22 | 5.6 10<sup>-21</sup> | Heart Left Ventricle | 1.8 10<sup>-05</sup> |
| rs11703971 (r<sup>2</sup>: 0.73) | SMARCB1 | 22 | **MMP11** | 22 | 1.1 10<sup>-14</sup> | Heart Atrial Appendage | 1.8 10<sup>-05</sup> |
| rs11703971 (r<sup>2</sup>: 0.73) | SMARCB1 | 22 | **SMARCB1** | 22 | 2.2 10<sup>-25</sup> | Heart Left Ventricle | 1.8 10<sup>-05</sup> |
| rs11703971 (r<sup>2</sup>: 0.73) | SMARCB1 | 22 | **SMARCB1** | 22 | 7.7 10<sup>-21</sup> | Heart Atrial Appendage | 1.8 10<sup>-05</sup> |
| rs11703971 (r<sup>2</sup>: 0.73) | SMARCB1 | 22 | **SMARCB1** | 22 | 4.9 10<sup>-24</sup> | Skeletal muscle | 1.8 10<sup>-05</sup> |
| rs11703971 (r<sup>2</sup>: 0.73) | SMARCB1 | 22 | **VPREB3** | 22 | 8.3 10<sup>-10</sup> | Heart Left Ventricle | 1.8 10<sup>-05</sup> |
| rs11703971 (r<sup>2</sup>: 0.73) | SMARCB1 | 22 | **VPREB3** | 22 | 8.4 10<sup>-6</sup> | Heart Atrial Appendage | 1.8 10<sup>-05</sup> |
| rs11703971 (r<sup>2</sup>: 0.73) | SMARCB1 | 22 | **VPREB3** | 22 | 1.2 10<sup>-17</sup> | Skeletal muscle | 1.8 10<sup>-05</sup> |

**Figure 4.**
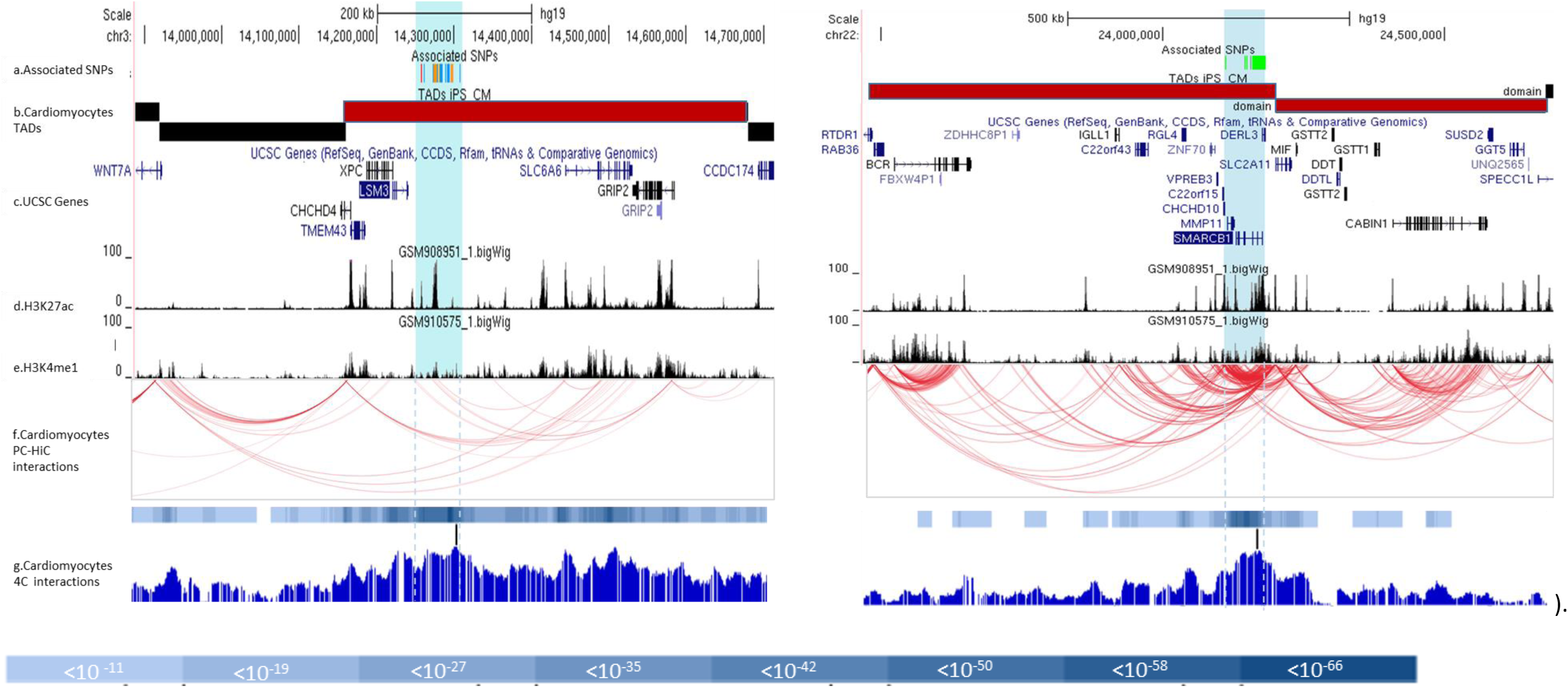
Positional candidate genes located in TAD domains at chromosome 3p25.1 (left) and 22q11.23 (right) loci. To delimitate the chromatin Topology Associated Domains (TADs) where genes are accessible to intra-TAD regulatory elements such as enhancers^(28)^, we first looked for publicly available Left Ventricle Topology Associating Domain^25^ (track b). TAD boundaries were comforted by the results of in-house Circular Chromatin Conformation Capture (4C)-Sequencing data produced on a iPSC-derived cardiomyocyte line from a donor (fully described in Supplemental Material iPSC reprogramming paragraph); 4C baits localization is schematized as a vertical black bar (track g). The p-values for interaction below 10^−8^ are shown as a blue scale colored bar given below. We then looked at the preferential chromatin interactions measured *via* PCHi-C (Promoter Chromatin Hi-C)^29^ on iPS derived cardiomyocytes (gene promoters predicted from GenHancer (https://www.genecards.org/)) that revealed preferential contact inside TADs as shown by the red curves (track f) and defined the intra-TAD candidate gene list (track c) prone to be regulated in *cis* by associated regions (blue highlight, track a. The color code for the SNP is identical to that of close figure 3). Specific DNA interactions are joining associated regions with histone enhancer marks (H3K27ac; H3K4me1, track d and e respectively

**Table 4.** SNP annotation. This excel table is given in a supplemental excel file.

The 4C-seq results from in-house iPSC-CM cell lines presented in **Figure 4Ag**, show more extended significant interaction (p<10^−8^) between the associated region bait [chr3:14,257,356-14,307,016] and intra-TAD regional promoters, confirming TAD boundaries and enhancer function of the chromosome 3p25.1 locus. The strongest interaction signals (dark blue scale figure 4Ag; p < 10^−50^, **Supplementary Table 6**) are localized on *SLC6A6, XPC/LSM3* region as well as in the intergenic region between those two genes where the bait is localized. Several baits specific of the 4C region and interacting with the associated SNPs block were designed to test for reverse interactions and confirmed the specificity of those *cis*-interaction signals (data not shown).

For each **Supplementary Table 5** positional candidate gene, we filtered out the intra-TAD better candidates based on cardiac expression data from GTEx, Montefiori *et al*.(32) and Henig *et al*.(33) as well as publicly available resources for gene annotation.

Looking at LV and atrial appendage expression, all those genes are expressed with from the most to the less expressed *TMEM43, CHCHD4, LSM3, SLC6A*6, *XPC* and *GRIP2* (**Supplementary Table 7A**). Moreover, *XPC* (p = 8.3 10^−15^) and *SLC6A6* (p = 6.9 10^−6^) LV expressions were significantly increased in DCM patients compared to healthy donors (**Supplementary Table 7A**) while *LSM3* expression was significantly decreased (p = 7.6 10^−8^). Functional candidate at this locus may include *TMEM43*, implied in arrhythmogenic right ventricular cardiomyopathy (ARVC)(39–41), and *SLC6A6*, for which a homozygous deletion affecting a splice site was found by whole-exome sequencing in a patient with idiopathic DCM(42).

Among the associated SNPs’ block, we then screened the GTEx database and Lemire *et al* results for eQTL, sQTL and mQTL(38). Even though we did not observe any evidence that the lead SNP, rs62232870, could influence the expression of nearby genes in relevant heart and skeletal muscle tissues, **Table 3** presents DCM associated SNPs, in moderate LD, associated with *SLC6A6* expression in heart atrial appendage. For example, rs7612736 (r^2^ = 0.71, D’ = 0.90, association p-value 1.6 10^−9^) moderately associates with that gene expression (p = 5.8 10^−5^). No sQTL was present but all the SNPs are associated with methylation level of several neighbor genes (**Table 4**). rs62232870 and the SNPs in strong LD with it are strongly associated with the methylation level of CpG site in *SLC6A6* and *TMEM43* (cg08926287 and cg15025640 with p-value around 10^−30^ and 10^−16^, respectively) while the rs4684185 LD SNP block associates primarily with *SLC6A6* and *XPC* methylation levels (cg08926287 and cg23070574 with p-value around 10^−70^ and 10^−36^, respectively).

Combining all the data available so far (**Table 4**), the best candidate to take shape at this locus is *SLC6A6* which is expressed in heart, differently between DCM and controls, and for which associated SNPs modulates methylation and expression (mQTLs and eQTLs).

### Candidate culprit gene selection strategy at chr22q11.23 locus

The LD block of DCM-associated SNPs (**Supplementary Figure 6** - **Supplementary Table 4**) extends over 70kb from the 5’ region of *MMP11* and *CHCHD10* to the 5’ region of *DERL3* including *SMARCB1* where the lead SNP maps to [chr22:24,110,180-24,182,174]. Multiple features witnessing the regulatory role of that locus, including LV enhancers(28), are observed (**Figure 3B)** and published ChIP-Seq experiments demonstrate that it is also strongly associated with H3K27ac and H3K4me1 LV marks (**Figure 3B**) providing support for an active enhancer function of that locus. TAD analysis at chromosome 22 showed that the associated SNPs block is located at the edge of 2 CM predicted TADs covering 1.2Mb [chr22:23,480,001-24,680,000] (**Figure 4B**). As multiple and strong intra- and inter-TADs interactions are found in that region (**Figure 4B**), the 21 genes covered by the two TADs should be considered as positional candidates (**Supplementary Table 5**).

Cardiomyocytes 4C-seq using the bait at chromosome 22 associated region revealed strong interactions with enhancer (H3K27ac/H3K4me1 positive) elements located close by (*SMARCB1, DERL3, MMP11, SLC2A11*, and *CHCHD10* principally), and to a lesser extent with the *BCR* gene locus and an intergenic 50kb 3’ of *IGLL1* and 18kb 3’ of *RGL4* and with the promoter rich region adjacent to the bait up to *CABIN1* (**Figure 4Bd-f**). The strongest 4C interaction signals (**Figure 4Bg**), are found for *SMARCB1* and *DERL3* (**Supplementary Table 8**; p < 10^−50^). Reverse interactions were tested with several baits confirming the specificity of those *cis*-interaction signals (data not shown).

Cardiac expression data showed that the most strongly expressed gene was *CHCHD10* (**Supplementary Table 7B**) followed by *GSTT1, DDT* and *SMARCB1* and, to a lesser extent, *CABIN1* and *SLC2A11*. The other 15 genes (*MIF, ZNF70, DERL3, MMP11* and further genes) appeared to be lower or not expressed. In LV tissue, *CHCHD10* and to a lesser extent *DDT, SMARCB1, SLC2A11*, and *CABIN1* were found to be differentially expressed between DCM and healthy donors (**Supplementary Table 7B**).

Among the association block, eQTL and sQTL were checked in GTEx database. **Table 3** presented the significant eSNP in relevant cardiac and skeletal tissues. From these 6 candidate genes, only *SMARCB1* expression was observed to be consistently strongly influenced by DCM associated SNPs, including the lead rs7284877 in the heart (**Supplementary Figure 10**) and skeletal muscle tissues. Three other genes, *MMP11, VPREB3*, and *DERL3*, even though not strongly expressed in heart, are also influenced by those associated SNPs in heart and/or skeletal muscle tissues (**Table 3**). No sQTL was present but rs7284877 and the SNPs in LD with it are associated with methylation level variation of several nearby genes (**Table 4**). The strongest regulation signals are detected for *SMARCB1* and *DERL3* (cg08219923 and cg25907215) with the best p-values below 10^−200^.

Combining all the data available so far (**Table 4**), *SMARCB1* that is expressed in heart, differentially in DCM and control heart explants and for which associated SNPs modulates methylation and expression (mQTLs and eQTLs) appears to be the strongest candidate.

## Discussion

By adopting a GWAS strategy, performed in the largest population of DCM assembled so far, we identified and replicated two new susceptibility loci for DCM while confirming two previously reported DCM associated ones, *HSPB7* and *BAG3*.

The first novel locus maps to chr3p25.1 where the minor A allele of the lead rs62232870 was associated with an increased OR of 1.36 [1.25 – 1.48] (p = 5.3 10^−13^) in the combined discovery and replication samples. The pattern of association observed at this locus extends over several genes among which two, *TMEM43* and *SLC6A6*, are expressed in heart and have previously been suspected to be involved in human structural cardiac disorders. Rare pathogenic mutations in the *TMEM43*, transmembrane protein 43, have been reported in arrhythmogenic right ventricular cardiomyopathy(39–41) while a homozygous splice site deletion has been observed in *SLC6A6* in an idiopathic DCM patient(42). Taurine (TAU) is a highly concentrated amino-acide with cyto-protective action in numerous tissues, especially in contractile ones such as heart(43). Oral TAU supplementation is cardio-protective, while depletion associates with dilated cardiomyopathy in several mammalian species(44,45). Intracellular TAU concentration relies on its transmembrane transporter (TauT) expression and activity encoded by *SLC6A6*. Accordingly, mice KO for *slc6a6* present tissue TAU level depletion and dilated cardiomyopathy(46) and a recent publication revealed that long-term treatment of dystrophic mice with taurine could prevent late heart dysfunction (47). It is thus tempting to speculate that *SLC6A6* expression regulation in humans could influence cardiac function as well. Our finding of a genetic association of *SLC6A6* alleles with DCM is remarkable in this context. Moreover, based on LV transcriptomic data, we observed that *SLC6A6* expression was decreased in DCM patients compared to controls and that, for several SNPs in LD with rs62232870-A lead-SNP, the allele associated with the rs62232870-A risk allele, showed a slight decrease in *SLC6A6* expression in heart atrial appendage (*i.e.* rs62231957 A allele is associated with the rs62232870-A risk allele and showed a decreased *SLC6A6* expression (p=1.9 10^−5^) (**Table 3** and **Supplementary Figure 11**).

As a whole, *in silico* annotation, chromatin interactions analysis showing specific interaction of the SNP LD block with SLC6A6 regulatory elements and the well-reported role of taurine in cardiac function strongly suggest *SLC6A6* as the most likely gene influencing DCM phenotype at the 3p25.1 locus. The underlying pathway leading to heart failure remains to be fully studied in humans, but our results may suggest the potential for therapeutic perspective through taurine administration or up-regulation.

The second novel DCM locus maps to chr22q11.23, where the rs7284877 lead SNP, intronic to *SMARCB1*, is associated with DCM, the C-risk allele presenting an OR of 1.33 [1.22 – 1.46] (p = 5.0 10^−10^). Our fine-mapping revealed that, among the positional candidates, six demonstrated significant expression in heart and difference in LV gene expression between DCM and healthy heart (from the highest to the lowest: *CHCHD10, GSTT1, DDT, SMARCB1, CABIN1*, and *SLC2A11*) among which only *SMARCB1* (SWI/SNF related matrix-associated actin-dependent regulator of chromatin subfamily b member 1) was under the influence of the lead rs7284877 in LV. Interestingly, the lead SNP is in complete LD with *SMARCB1*-rs5760054 recently reported to associate with LV internal dimension in systole and fractional shortening in a multi-trait GWAS conducted in 162,255 Japanese individuals(48). Even though not regulated by rs7284877, two other highlighted genes were already implied in cardiovascular disorders and/or phenotypes. First, *GSTT1*, glutathione S-transferase theta 1, has been suggested to associate with various cardiovascular diseases(49,50). It is shown *via* RNA-seq studies to be differentially expressed in heart depending on the pathological state: overexpression in male left ventricular hypertrophy (LVF) but downregulation in DCM heart compared to control(51); overexpression in DCM or non-failure *vs.* ischemic heart(52); Second, *CABIN1*, calcineurin binding protein 1, is a protein that binds to and inhibits calcineurin (CaN), a calcium-regulated phosphatase. CABIN1 is cleaved by calpain that is upregulated in heart failure LV. Calpain upregulation may increase cabin1 cleavage and thus lead to CaN overexpression in HF patients which can contribute to cardiac remodeling (53,54). Two other genes at this locus, *DERL3* and *MMP11*, were not included in our candidates because of their low cardiac expression but could nonetheless be interesting candidates. *DERL3*-rs5760061, in complete LD with rs7284877, associated with LV internal dimension and fractional shortening in the Japanese GWAS. Another SNP of this gene, rs6003909, in high LD (r2=0.8) with rs7284877, also associated with LV geometric remodeling in a UK Biobank GWAS of LV image-derived phenotypes (55). We also found that another SNP, rs2186370, in complete LD with the lead SNP, is a mQTL strongly associated (p=8.4 10^−71^) with the methylation level of CpG cg25907215 lying in the 3’UTR of *DERL3. DERLIN3* (*DERL3*) plays a role in heart homeostasis maintenance (56,57) and its expression in skeletal muscle is under the control of DCM associated SNPs including the lead. *MMP11* has an expression consistently influenced by rs7284877. Interestingly, unlike the 10 other rs7284877 cis-regulated expressions, the eQTL activity on *MMP11* is quasi restricted to cardiac tissues (p= 4 10^−20^, 5.7 10^−15^ and 7.4 10^−5^ in heart left ventricle, heart atrial appendage, and cells-cultured fibroblast, respectively). *MMP11*, matrix metallopeptidase 11, encodes an enzyme implied in the regulation of extracellular matrix and in the development of myocardial fibrosis and ventricular remodeling(58,59) functions that could be related with DCM. Even though we did not select a striking candidate at this locus, *SMARCB1*, whose expression and methylation are directly regulated by the lead SNP and associates with various cardiac phenotypes seems to be the stronger.

However, for both of these DNA loci with a regulatory function, it is also plausible that a single gene does not explain the whole functional activity detected(60). Given the strong bi-allelic activity of the enhancer regions in both loci and interaction shown in 4C data, multiple genes expressed in cardiomyocytes might be regulated in parallel to produce DCM phenotype.

Beyond the discovery of two novel DCM-associated loci, this GWAS investigation provided an innovative estimate of the heritability of the disease in European descent individuals, h^2^ = 31% ±8%, a value consistent with that (h^2^ ∼30%) recently reported in a population of African origin(12). However, the four independent lead SNPs (*BAG3, HSPB7*, chr3p25.1-*SLC6A6*, and chr22q11.23-*SMARCB1*) reaching genome-wide significance only contribute to 2% of the heritability suggesting the role of additional genetic factors or gene/gene and gene/environment interactions yet to be identified. Despite their modest contribution to the DCM heritability, these 4 SNPs may have clinical utility. Indeed, a GRS constructed on these 4 SNPs revealed that individuals with 8 risk alleles were at 30% increased risk of DCM and those with 1 risk allele at 19% reduced risk of DCM compared to individuals with 5 the risk alleles referral. This GRS, the first of its kind developed in the field of DCM, may have practical implications for better management of subjects at risk for DCM or systolic dysfunction, such as patients with drugs increasing the risk of myocardial dysfunction, or relatives in DCM families. Further clinical studies are warranted to validate its clinical utility.

Despite its innovative findings in the context of DCM, this study suffers from some limitations. First, we robustly identified two new DCM loci but were not able to fully demonstrate which culprit variant(s) and gene(s) are directly responsible for the observed susceptibility to the disease. Further intensive molecular and cellular investigations are needed to fill this gap. Second, despite being the largest GWAS ever performed on DCM, with both a discovery and a replication phase, our GWAS may have been suboptimal in identifying common susceptibility DCM alleles due to the absence of perfectly matched healthy controls for the British and US populations. Therefore, we performed our discovery GWAS on combined individual data while handling any potential hidden population stratification through adjustment on genetically derived principal components. Nevertheless, the robust replication of 2 out of 3 genome-wide significant associations in two case-control studies of Dutch and German origins provides strong support for the validity of our general strategy. The replication of the reported genetic associations in non-European ancestry populations is now needed. Similarly, the in-depth interrogation of electronic health records database with GWAS data, such as UK biobank (https://www.ukbiobank.ac.uk/)(61) or the Million Veteran Program (https://www.research.va.gov/mvp/)(62), and DCM information, may increase the power to detect additional DCM associated loci.

In conclusion, we identified two new genetic loci associated with DCM. While *SLC6A6* clearly stands out as the most likely candidate at the chr3p25.1 locus, several plausible candidates *SMARCB1, DERL3*, and *MMP11* could be proposed at chr22q11.23 locus. A genetic risk score was built with a potential clinical application for the prediction of systolic heart failure. These findings add both on the understanding of the genetic architecture of DCM and on potential new players/pathways involved in the pathophysiology of systolic heart failure.

## Supporting information

Table4_SNPannotation

## Supplementary Tables and Figures

**Supplementary Figure 1.** QQplot summarizing the discovery GWAS results

**Supplementary Figure 2 to 6.** Regional-association-plot at chromosome 1, 3, 10, 16 and 22

**Supplementary Figure 7-8.** Manhattan and QQ summarizing the discovery GWAS conditional-analysis results

**Supplementary Figure 9.** Barplot of the weighted scores grouped by quintile

**Supplementary Figure 10.** Violin-plot showing the regulation of *SMARCB1* (a) and *MMP11* (b) expression by rs7284877 lead SNP

**Supplementary Figure 11.** Violin-plot showing the regulation of *SLC6A* by rs62231950

**Supplementary Table 1.** Genotyping methods and quality controls

**Supplementary Table 2.** Haplotypic-association signal at chromosome 3 locus

**Supplementary Table 3.** Unweighted (A) and weighted (B) scores replications

**Supplementary Table 4.** SNPs’ LD block at chromosome 3 and 22 loci

**Supplementary Table 5.** List of the gene at chromosome 3 and 22 loci

**Supplementary Table 6.** 4C interaction at chromosome 3 locus

**Supplementary Table 7.** Gene Expression level at chromosome 3 and 22 loci

**Supplementary Table 8.** 4C interaction at chromosome 22 locus

**Supplementary Table 9.** 4C-seq primers

**Table 4: SNP annotation.** This excel table is given in a supplemental excel file.

## References

1. Elliott P, Andersson B, Arbustini E, Bilinska Z, Cecchi F, Charron P, et al. Classification of the cardiomyopathies: a position statement from the European Society Of Cardiology Working Group on Myocardial and Pericardial Diseases. Eur Heart J. janv 2008;29(2):270–6.

2. Jefferies JL, Towbin JA. Dilated cardiomyopathy. Lancet Lond Engl. 27 févr 2010;375(9716):752–62.

3. Tayal U, Prasad S, Cook SA. Genetics and genomics of dilated cardiomyopathy and systolic heart failure. Genome Med. 22 2017;9(1):20.

4. Pinto YM, Elliott PM, Arbustini E, Adler Y, Anastasakis A, Böhm M, et al. Proposal for a revised definition of dilated cardiomyopathy, hypokinetic non-dilated cardiomyopathy, and its implications for clinical practice: a position statement of the ESC working group on myocardial and pericardial diseases. Eur Heart J. 14 2016;37(23):1850–8.

5. McNally EM, Golbus JR, Puckelwartz MJ. Genetic mutations and mechanisms in dilated cardiomyopathy. J Clin Invest. janv 2013;123(1):19–26.

6. Harakalova M, Kummeling G, Sammani A, Linschoten M, Baas AF, van der Smagt J, et al. A systematic analysis of genetic dilated cardiomyopathy reveals numerous ubiquitously expressed and muscle-specific genes. Eur J Heart Fail. mai 2015;17(5):484–93.

7. Stark K, Esslinger UB, Reinhard W, Petrov G, Winkler T, Komajda M, et al. Genetic association study identifies HSPB7 as a risk gene for idiopathic dilated cardiomyopathy. PLoS Genet. 21 oct 2010;6(10):e1001167.

8. Cappola TP, Li M, He J, Ky B, Gilmore J, Qu L, et al. Common variants in HSPB7 and FRMD4B associated with advanced heart failure. Circ Cardiovasc Genet. avr 2010;3(2):147–54.

9. Villard E, Perret C, Gary F, Proust C, Dilanian G, Hengstenberg C, et al. A genome-wide association study identifies two loci associated with heart failure due to dilated cardiomyopathy. Eur Heart J. mai 2011;32(9):1065–76.

10. Meder B, Rühle F, Weis T, Homuth G, Keller A, Franke J, et al. A genome-wide association study identifies 6p21 as novel risk locus for dilated cardiomyopathy. Eur Heart J. avr 2014;35(16):1069–77.

11. Esslinger U, Garnier S, Korniat A, Proust C, Kararigas G, Müller-Nurasyid M, et al. Exome-wide association study reveals novel susceptibility genes to sporadic dilated cardiomyopathy. PloS One. 2017;12(3):e0172995.

12. Xu H, Dorn GW, Shetty A, Parihar A, Dave T, Robinson SW, et al. A Genome-Wide Association Study of Idiopathic Dilated Cardiomyopathy in African Americans. J Pers Med. 26 2018;8(1).

13. Aragam KG, Chaffin M, Levinson RT, McDermott G, Choi S-H, Shoemaker MB, et al. Phenotypic Refinement of Heart Failure in a National Biobank Facilitates Genetic Discovery. Circulation. 11 nov 2018;

14. Norton N, Li D, Rieder MJ, Siegfried JD, Rampersaud E, Züchner S, et al. Genome-wide studies of copy number variation and exome sequencing identify rare variants in BAG3 as a cause of dilated cardiomyopathy. Am J Hum Genet. 11 mars 2011;88(3):273–82.

15. Henry WL, Gardin JM, Ware JH. Echocardiographic measurements in normal subjects from infancy to old age. Circulation. nov 1980;62(5):1054–61.

16. Mestroni L, Maisch B, McKenna WJ, Schwartz K, Charron P, Rocco C, et al. Guidelines for the study of familial dilated cardiomyopathies. Collaborative Research Group of the European Human and Capital Mobility Project on Familial Dilated Cardiomyopathy. Eur Heart J. janv 1999;20(2):93–102.

17. Sammani A, Jansen M, Linschoten M, Bagheri A, de Jonge N, Kirkels H, et al. UNRAVEL: big data analytics research data platform to improve care of patients with cardiomyopathies using routine electronic health records and standardised biobanking. Neth Heart J Mon J Neth Soc Cardiol Neth Heart Found. 27 mai 2019;

18. Angelow A, Schmidt M, Hoffmann W. Towards risk factor assessment in inflammatory dilated cardiomyopathy: the SFB/TR 19 study. Eur J Cardiovasc Prev Rehabil Off J Eur Soc Cardiol Work Groups Epidemiol Prev Card Rehabil Exerc Physiol. oct 2007;14(5):686–93.

19. Völzke H, Alte D, Schmidt CO, Radke D, Lorbeer R, Friedrich N, et al. Cohort profile: the study of health in Pomerania. Int J Epidemiol. avr 2011;40(2):294–307.

20. Li Y, Willer CJ, Ding J, Scheet P, Abecasis GR. MaCH: using sequence and genotype data to estimate haplotypes and unobserved genotypes. Genet Epidemiol. déc 2010;34(8):816–34.

21. Li Y, Willer C, Sanna S, Abecasis G. Genotype imputation. Annu Rev Genomics Hum Genet. 2009;10:387–406.

22. Tregouet DA, Garelle V. A new JAVA interface implementation of THESIAS: testing haplotype effects in association studies. Bioinforma Oxf Engl. 15 avr 2007;23(8):1038–9.

23. Willer CJ, Li Y, Abecasis GR. METAL: fast and efficient meta-analysis of genomewide association scans. Bioinforma Oxf Engl. 1 sept 2010;26(17):2190–1.

24. Zheng J, Erzurumluoglu AM, Elsworth BL, Kemp JP, Howe L, Haycock PC, et al. LD Hub: a centralized database and web interface to perform LD score regression that maximizes the potential of summary level GWAS data for SNP heritability and genetic correlation analysis. Bioinforma Oxf Engl. 15 2017;33(2):272–9.

25. Kent WJ, Sugnet CW, Furey TS, Roskin KM, Pringle TH, Zahler AM, et al. The human genome browser at UCSC. Genome Res. juin 2002;12(6):996–1006.

26. ENCODE Project Consortium. An integrated encyclopedia of DNA elements in the human genome. Nature. 6 sept 2012;489(7414):57–74.

27. Saha A, Kim Y, Gewirtz ADH, Jo B, Gao C, McDowell IC, et al. Co-expression networks reveal the tissue-specific regulation of transcription and splicing. Genome Res. 2017;27(11):1843–58.

28. Leung D, Jung I, Rajagopal N, Schmitt A, Selvaraj S, Lee AY, et al. Integrative analysis of haplotype-resolved epigenomes across human tissues. Nature. 19 févr 2015;518(7539):350–4.

29. Lesurf R, Cotto KC, Wang G, Griffith M, Kasaian K, Jones SJM, et al. ORegAnno 3.0: a community-driven resource for curated regulatory annotation. Nucleic Acids Res. 4 janv 2016;44(D1):D126–132.

30. Blanchette M, Kent WJ, Riemer C, Elnitski L, Smit AFA, Roskin KM, et al. Aligning multiple genomic sequences with the threaded blockset aligner. Genome Res. avr 2004;14(4):708–15.

31. Dixon JR, Gorkin DU, Ren B. Chromatin Domains: The Unit of Chromosome Organization. Mol Cell. 02 2016;62(5):668–80.

32. Montefiori LE, Sobreira DR, Sakabe NJ, Aneas I, Joslin AC, Hansen GT, et al. A promoter interaction map for cardiovascular disease genetics. eLife. 10 2018;7.

33. Heinig M, Adriaens ME, Schafer S, van Deutekom HWM, Lodder EM, Ware JS, et al. Natural genetic variation of the cardiac transcriptome in non-diseased donors and patients with dilated cardiomyopathy. Genome Biol. 14 2017;18(1):170.

34. Wang K, Li M, Hakonarson H. ANNOVAR: functional annotation of genetic variants from high-throughput sequencing data. Nucleic Acids Res. sept 2010;38(16):e164.

35. Boyle AP, Hong EL, Hariharan M, Cheng Y, Schaub MA, Kasowski M, et al. Annotation of functional variation in personal genomes using RegulomeDB. Genome Res. sept 2012;22(9):1790–7.

36. Smedley D, Schubach M, Jacobsen JOB, Köhler S, Zemojtel T, Spielmann M, et al. A Whole-Genome Analysis Framework for Effective Identification of Pathogenic Regulatory Variants in Mendelian Disease. Am J Hum Genet. 01 2016;99(3):595–606.

37. Kircher M, Witten DM, Jain P, O’Roak BJ, Cooper GM, Shendure J. A general framework for estimating the relative pathogenicity of human genetic variants. Nat Genet. mars 2014;46(3):310–5.

38. Lemire M, Zaidi SHE, Ban M, Ge B, Aïssi D, Germain M, et al. Long-range epigenetic regulation is conferred by genetic variation located at thousands of independent loci. Nat Commun. 26 févr 2015;6:6326.

39. Honda T, Kanai Y, Ohno S, Ando H, Honda M, Niwano S, et al. Fetal arrhythmogenic right ventricular cardiomyopathy with double mutations in TMEM43. Pediatr Int Off J Jpn Pediatr Soc. mai 2016;58(5):409–11.

40. Siragam V, Cui X, Masse S, Ackerley C, Aafaqi S, Strandberg L, et al. TMEM43 mutation p.S358L alters intercalated disc protein expression and reduces conduction velocity in arrhythmogenic right ventricular cardiomyopathy. PloS One. 2014;9(10):e109128.

41. Bao J, Wang J, Yao Y, Wang Y, Fan X, Sun K, et al. Correlation of ventricular arrhythmias with genotype in arrhythmogenic right ventricular cardiomyopathy. Circ Cardiovasc Genet. déc 2013;6(6):552–6.

42. Shakeel M, Irfan M, Khan IA. Rare genetic mutations in Pakistani patients with dilated cardiomyopathy. Gene. 5 oct 2018;673:134–9.

43. Xu Y-J, Arneja AS, Tappia PS, Dhalla NS. The potential health benefits of taurine in cardiovascular disease. Exp Clin Cardiol. 2008;13(2):57–65.

44. Pion PD, Kittleson MD, Rogers QR, Morris JG. Myocardial failure in cats associated with low plasma taurine: a reversible cardiomyopathy. Science. 14 août 1987;237(4816):764–8.

45. Kaplan JL, Stern JA, Fascetti AJ, Larsen JA, Skolnik H, Peddle GD, et al. Taurine deficiency and dilated cardiomyopathy in golden retrievers fed commercial diets. PloS One. 2018;13(12):e0209112.

46. Ito T, Oishi S, Takai M, Kimura Y, Uozumi Y, Fujio Y, et al. Cardiac and skeletal muscle abnormality in taurine transporter-knockout mice. J Biomed Sci. 24 août 2010;17 Suppl 1:S20.

47. Mele A, Mantuano P, De Bellis M, Rana F, Sanarica F, Conte E, et al. A long-term treatment with taurine prevents cardiac dysfunction in mdx mice. Transl Res J Lab Clin Med. 2019;204:82–99.

48. Kanai M, Akiyama M, Takahashi A, Matoba N, Momozawa Y, Ikeda M, et al. Genetic analysis of quantitative traits in the Japanese population links cell types to complex human diseases. Nat Genet. mars 2018;50(3):390–400.

49. Song Y, Shan Z, Luo C, Kang C, Yang Y, He P, et al. Glutathione S-Transferase T1 (GSTT1) Null Polymorphism, Smoking, and Their Interaction in Coronary Heart Disease: A Comprehensive Meta-Analysis. Heart Lung Circ. avr 2017;26(4):362–70.

50. García-González I, López-Díaz RI, Canché-Pech JR, Solís-Cárdenas A de J, Flores-Ocampo JA, Mendoza-Alcocer R, et al. Epistasis analysis of metabolic genes polymorphisms associated with ischemic heart disease in Yucatan. Clin E Investig En Arterioscler Publicacion Of Soc Espanola Arterioscler. juin 2018;30(3):102–11.

51. Newman MS, Nguyen T, Watson MJ, Hull RW, Yu H-G. Transcriptome profiling reveals novel BMI- and sex-specific gene expression signatures for human cardiac hypertrophy. Physiol Genomics. 1 juill 2017;49(7):355–67.

52. Liu Y, Morley M, Brandimarto J, Hannenhalli S, Hu Y, Ashley EA, et al. RNA-Seq identifies novel myocardial gene expression signatures of heart failure. Genomics. févr 2015;105(2):83–9.

53. Yang D, Ma S, Tan Y, Li D, Tang B, Zhang X, et al. Increased expression of calpain and elevated activity of calcineurin in the myocardium of patients with congestive heart failure. Int J Mol Med. juill 2010;26(1):159–64.

54. Yang Y, Zhu W, Zhou X, Zhang X, Zhu Z. [Calpain involved in signal transduction of myocardial remodeling in patients with congestive heart failure]. Zhonghua Xin Xue Guan Bing Za Zhi. mars 2005;33(3):247–50.

55. Aung N, Vargas JD, Yang C, Cabrera CP, Warren HR, Fung K, et al. Genome-Wide Analysis of Left Ventricular Image-Derived Phenotypes Identifies Fourteen Loci Associated With Cardiac Morphogenesis and Heart Failure Development. Circulation. 15 oct 2019;140(16):1318–30.

56. Belmont PJ, Chen WJ, San Pedro MN, Thuerauf DJ, Gellings Lowe N, Gude N, et al. Roles for endoplasmic reticulum-associated degradation and the novel endoplasmic reticulum stress response gene Derlin-3 in the ischemic heart. Circ Res. 5 févr 2010;106(2):307–16.

57. Glembotski CC. Roles for ATF6 and the sarco/endoplasmic reticulum protein quality control system in the heart. J Mol Cell Cardiol. juin 2014;71:11–5.

58. Li YY, McTiernan CF, Feldman AM. Interplay of matrix metalloproteinases, tissue inhibitors of metalloproteinases and their regulators in cardiac matrix remodeling. Cardiovasc Res. mai 2000;46(2):214–24.

59. Creemers EE, Cleutjens JP, Smits JF, Daemen MJ. Matrix metalloproteinase inhibition after myocardial infarction: a new approach to prevent heart failure? Circ Res. 3 août 2001;89(3):201–10.

60. Haitjema S, Meddens CA, van der Laan SW, Kofink D, Harakalova M, Tragante V, et al. Additional Candidate Genes for Human Atherosclerotic Disease Identified Through Annotation Based on Chromatin Organization. Circ Cardiovasc Genet. avr 2017;10(2).

61. Sudlow C, Gallacher J, Allen N, Beral V, Burton P, Danesh J, et al. UK biobank: an open access resource for identifying the causes of a wide range of complex diseases of middle and old age. PLoS Med. mars 2015;12(3):e1001779.

62. Gaziano JM, Concato J, Brophy M, Fiore L, Pyarajan S, Breeling J, et al. Million Veteran Program: A mega-biobank to study genetic influences on health and disease. J Clin Epidemiol. févr 2016;70:214–23.

