## Supplemental-Data for "Genome wide association analysis in dilated cardiomyopathy reveals two new key players in systolic heart failure on chromosome 3p25.1 and 22q11.23"

### Supplemental Material

### Cohort description

#### Discovery GWAS cohorts

The discovery phase built upon 2,719 DCM cases coming from 5 populations (France, Germany, Italy, United Kingdom, and the USA) and 4,440 controls coming from 3 populations (France, Germany and Italy).

*French DCM samples.* 696 DCM patients from 3 national clinical studies, CARDIGENE (n=408), EUROGENE (EHF) (n=84) and PHRC (n=204), were involved in the current GWAS.

**CARDIGENE study**(1). Cases were French patients with a diagnosis of idiopathic DCM (enlarged left ventricle end-diastolic volume/diameter, LVEDD, >140 ml/m$^2$ on ventriculography or >34 mm/m$^2$ on echocardiography and low ejection fraction, EF, ≤40% confirmed over a six-month period, in the absence of causal factors such as coronary artery disease (coronary angiography was mandatory if DCM occurred after 35 years of age) or sustained arterial hypertension, intrinsic valvular disease, documented myocarditis, congenital malformation, and insulin-dependent diabetes. Only apparently sporadic DCM cases without an additional (first-degree) relative with DCM were included. All were of European origin (born in France, from parents born in France or neighboring countries).

Recruitment was performed in ten hospitals from six French regions (Lille, Lyon, Nancy, Nantes, Paris-Ile de France, and Strasbourg) from September 1994 to February 1996.

A total of 408 DCM cases were included (322 men and 86 women, among them 212 had undergone cardiac transplantation). The mean age of patients at diagnosis was 45.1 ± 10.7 years, mean left ventricular ejection fraction (LVEF) was 24.0 ± 8.2% and mean end-diastolic volume was 8.68 ± 9.86 mm. The study was supported by grants from Délégation à la recherche clinique AP-HP (EMUL and PHRC n°AOM95082).

**EUROGENE study (EHF)**(2)**.** All cases were patients of European origin (all born in Europe, from parents and grandparents born in France or neighboring countries) with a diagnosis of idiopathic DCM, i.e. left ventricle end-diastolic volume/diameter >117% of predicted value according to age and body surface area on echocardiography and low ejection fraction (<45%) confirmed over a three-month period, in the absence of causal factors such as coronary artery disease (coronary angiography or coronary CT scan was mandatory if DCM occurred after 35 years of age) or intrinsic valvular disease, documented myocarditis, systemic disease, sustained rapid supraventricular arrhythmia, or congenital malformation. Recruitment was performed in 11 hospitals in seven European countries from September 2000 to February 2005. French patients originated from 3 centers in Paris-Ile de France and a total of 83 DCM cases were included (66 men and 18 women, no patient had undergone cardiac transplantation at inclusion). The mean age of patients at diagnosis was 46.9 ± 13.2 years, mean LVEF was 29.2 ± 10.2% and mean end-diastolic diameter was 67.5 ± 8.1 mm. The study was supported by grants from the “Fondation LEDUCQ”.

**PHRC study.** DCM cases were French patients of European origin (all born in France, from parents and grandparents born in France or neighboring countries; some patients of Maghreb origin were retrospectively excluded) with a diagnosis of idiopathic DCM enlarged left ventricle end-diastolic volume/diameter >117% of predicted value according to age and body surface area on echocardiography and low ejection fraction (<45%) clinically stable over a three-month period, in the absence of causal factors such as coronary artery disease (coronary angiography or coronary CT scan was mandatory if DCM occurred after 35 years of age) or intrinsic valvular disease, documented myocarditis, or congenital malformation. Only apparently sporadic DCM cases without additional (first degree) relative with DCM were included. Recruitment was performed in eight hospitals in six regions in France (Lille, Lyon, Nantes, Nice, Paris-Ile de France, and Tours) from October 2005 to November 2008. A total of 204 DCM cases were included (163 men and 41 women, no patient had undergone cardiac transplantation at inclusion). The mean age of patients at diagnosis was 52.0 ± 13.0 years, mean LVEF was 28.2 ± 8.9% and mean end-diastolic diameter was 68.5 ± 9.0 mm. The study was supported by grants from "Programme Hospitalier de Recherche Clinique" (PHRC n°AOM 04141).

German DCM samples. 1,201 DCM cases coming from two German cohorts one originating from Berlin (n=987) and the EHF study(2) (n = 214) were included.

**Berlin Cohort.** German idiopathic DCM cases were recruited at the German Heart Institute Berlin and were of white European origin. Inclusion criteria for DCM cases were the following: reduced systolic function (LVEF <45%), after exclusion of major coronary artery disease (by angiography), significant (>grade 2) valvular heart disease, hypertensive heart disease, congenital heart disease, myocarditis or other secondary forms of heart failure.

Patients with a positive family history were also excluded. A total of 987 DCM patients were included (815 men and 172 women). Mean age of patients at diagnosis was 43.97 ± 11.6 years, mean LVEF was 24.15 ± 9.7% and mean end-diastolic diameter was 68 ± 10.0 mm

**German EUROGENE (EHF)**(2) **cases.** DCM definition was the same as the one used above for the EHF French patients. German patients were recruited in 3 centers (Regensburg, Marburg and Munster). A total of 214 DCM cases were included (170 men and 44 women, only 1 patient had undergone a cardiac transplantation at inclusion). The mean age of patients at diagnosis was 45.9 ± 11.7 years, mean LVEF was 30.3 ± 11.3% and mean end-diastolic diameter was 69.8 ± 9.7 mm.

*UK DCM samples.* Patients referred to the Royal Brompton and Harefield Hospitals NHS Foundation Trust (RBHT) cardiovascular magnetic resonance (CMR) unit from July 2001 to August 2012 for evaluation of a possible diagnosis of DCM and who agreed to provide samples for biobanking were prospectively recruited at the National Institute for Health Research Cardiovascular Biomedical Research Unit, RBHT and Imperial College London. Referrals were from centers across Southern England. A diagnosis of DCM was confirmed, and evaluated against published CMR criteria (ejection fraction >2sd below and end-diastolic volume >2sd above the mean normalized for age and sex) by two independent Level 3 accredited CMR cardiologists. No patients had clinical symptoms or signs of active myocarditis or CMR evidence of infiltrative disease. Significant coronary artery disease (CAD) (>50% diameter luminal stenosis in any coronary artery) was excluded by coronary angiography. A total of 109 DCM cases were included (79 men and 30 women). The mean age of patients at diagnosis was 54.9 ± 13.4, mean LVEF was 30 ± 8.7.

*Italian DCM samples.* DCM patients were selected from the (EHF)(2) study. 82 cases (67 men and 15 women) were recruited from the city of Pavia. No patient had undergone a cardiac transplantation at inclusion. The mean age of patients at diagnosis was 43.1 ± 13.4 years, mean LVEF was 27.2 ± 7.9%, and mean end-diastolic diameter was 66.5 ± 8.5 mm.

*US DCM samples.* 631 patients with dilated cardiomyopathy, defined as patients with heart failure and an ejection fraction < 40% in the absence of hypertension, primary valvular disease, or coronary artery disease were recruited from the Myocardial Applied Genomics Network (MAGNet) Study**.** All subjects provided written informed consent using protocols approved by relevant institutional review boards. Whole-genome SNP genotypes were generated in the Center for Applied Genomics (Philadelphia, USA), using the Illumina *HumanOmniExpressExome8v1_A BeadChip*. Genotype calling was done using *GenomeStudio* v2011.1.

*PPS3 study*(3)**.** 1,084 French controls (731 men and 353 women) were selected from the Paris Prospective Study III (PPS3), which is an ongoing French prospective study. From June 2008 to May 2012, 10,157 men and women aged 50-75 years who had a preventive medical check-up at the Centre d'Investigations Préventives et Cliniques in Paris, were enrolled in the PPS3 study, after signing informed consent. A detailed study report is described elsewhere(3). The mean age of the participants in 2012 was 62 ± 6.4 years

*KORA F4.* 3,264 German controls (1579 men and 1685 women) were selected from the KORA F4 study (Cooperative Health Research in the Augsburg Region) which is a 7-year follow-up study to the population-based KORA Survey S4(4,5). The baseline survey KORA S4 was conducted in the years 1999–2001 in the city of Augsburg, Southern Germany, and its two adjacent counties. All survey participants are residents of German nationality identified through the registration office within the age range 25-74 years and gave written consent. The mean age of the participants was 57.4 ± 12.9 years. Genotyping was performed at the Helmholtz Zentrum (Munich, Germany) and KORA F4 data transmitted were imputed ones. Only genotyped data (587 050) were kept for the combined analysis of all populations

*Italian control subject*

Control subjects for Italian DCM cases were selected from healthy consultants or hospital professional workers in clinical centers in Italy (92 controls, 70 men and 22 women) and were part of the EHF study(2). Mean age of the participants in 2012 was 49.8 ± 12.7 years

*Replication cohorts*

**iGeneTRAIN HTx.** Three iGeneTRAiN HTx cohorts from Madrid, Utrecht and Pennsylvania totalizing 145 cases and 527 controls, selected to be more than 18 years of age, unrelated and of European descent were used as DCM replication datasets. The Madrid cohort included 51 heart transplanted idiopathic DCM patients (7 females, 44 males) and 139 heart transplant donors from the heart transplant program at Hospital Universitario Puerta de Hierro. Authorization was obtained from Hospital Universitario Puerta de Hierro ethics committee. All samples were of European descent and unrelated. The mean age at transplantation was 47.2 (standard deviation 12.0). For the Utrecht cohort, 50 DCM patients who underwent heart transplantation (39 male, 11 female) and 145 heart transplant donor controls were included. All patients were transplanted at the University Medical Center Utrecht, the Netherlands, and were included in the UNRAVEL research data platform(6). The study was approved by the local ethics committee, and all patients provided written informed consent. Samples were unrelated and of European ancestry, mean age at transplantation 47.2±10.6. The Penn Heart Tx study consists of adult heart transplant recipients and matching donors.

All patients underwent cardiac transplantation in the Penn Transplant Institute and written informed consent was obtained. For the current study, we included 44 DCM cases and 243 healthy donors as controls. All samples were of European descent. Twelve DCM patients were female, 32 were male, mean age was 54.3±12.4 and all samples were unrelated.

**German.** All cases were Caucasian participants of the SFB-TR19 registry(7) which enrolled patients with suspected DCM. In total, 439 subjects with genetic data (350 men and 89 women, mean age 54.4 ± 11.8 years) and a low left ventricular ejection fraction (<45%) were included in the analysis. Among them, 394 patients had a dilated left ventricular end-diastolic diameter (defined as >117% of predicted value according to age and body surface area) and 45 subjects not meeting the dilatation criteria and therefore classified as hypokinetic non-dilated cardiomyopathy (HNDC). 439 age- and sex-matched controls (350 men and 89 women with a mean age of 54.4 ± 11.6 years) were selected from the population-based Study of Health in Pomerania (SHIP-1)(8). All control subjects had a normal left ventricular ejection fraction (>60%) and a normal left ventricular end-diastolic diameter as determined by echocardiography.

**Gender Repartition in the 5 cohorts**

| | | N | Males N(%) | Females N(%) |
|---|---|---|---|---|
| **Germany** | Controls | 3,264 | 1,579 (48.4%) | 1685 (51.6%) |
| | Cases | 1,165 | 956 (82.1%) | 209 (17.9%) |
| **France** | Controls | 973 | 656 (67.4%) | 317 (32.6%) |
| | Cases | 673 | 532 (79.01%) | 141 (20.9%) |
| **Italy** | Controls | 92 | 69 (75.0%) | 23 (25.0%) |
| | Cases | 78 | 63 (80.8%) | 15 (19.2%) |
| **United Kingdom** | Cases | 103 | 75 (72.8%) | 28 (27.2%) |
| **USA** | Cases | 631 | 420 (66. 6%) | 211 (33.4%) |

**Genotyping**

Detailed descriptions of the used genotyping arrays as well as the adopted quality control procedures are shown in **Supplementary Table 1A and B**.

**General framework for imputation and association analysis**

Quality control was performed in each population separately with the 1.9 version of the PLINK software(9) (http://pngu.mgh.harvard.edu/~purcell/plink/plink2.shtml) as described in **Supplementary Table 1**. Subsequently, all autosomal SNP data were merged and a similar QC procedure was adopted to identify genotyped SNPs shared in all individuals of the discovery cohorts (n = 557,776). This procedure identified 7 pairs of duplicated individuals, for which only one individual per pair was then kept, and 13 individuals with call rate <95% that were later discarded. A further round of QC was performed to identify additional genetic outliers.

Based on the analysis of IBS state distance matrix, 149 additional samples (44 cases and 105 controls) were discarded leaving 6,980 (2,651 cases and 4,329 controls) individuals for imputation and association analysis (**Figure A** and **B**, below). To minimize the risk of ambiguous SNPs (A/T or G/C) during the imputation step, such SNPs (n = 1519) were removed prior to imputation analyses leaving 554,257 autosomal variants. For X chromosome SNP data, the application of a similar procedure led to the selection of 10 471 SNPs in 6,874 samples.

***Figure A****: Matrix distance plot before (left) and after (right) outlier exclusion. The numbered red dots correspond to the individuals that were removed considered outlier and removed.*

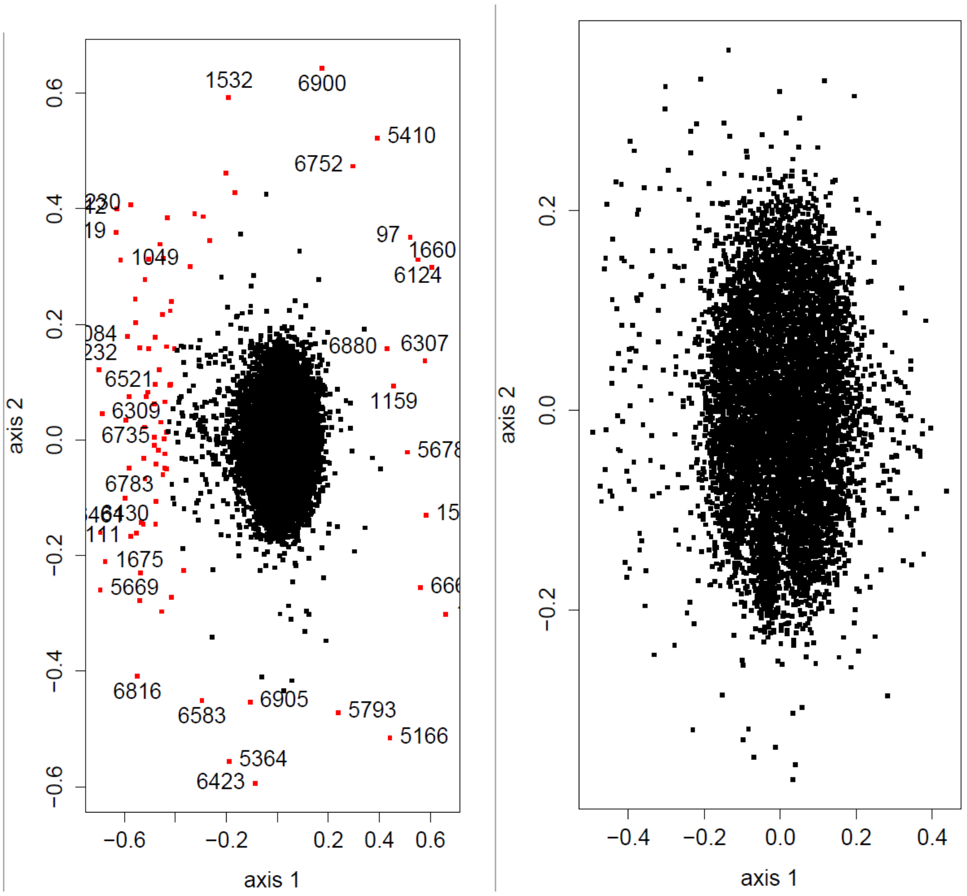

***Figure B****: First two axes of PCA analysis after exclusion of the 149 individuals showing the genetic homogeneity between the 5 populations (colored dots represent samples from: black, Germany; green: France; red: UK; light blue: US and dark blue: Italy)*

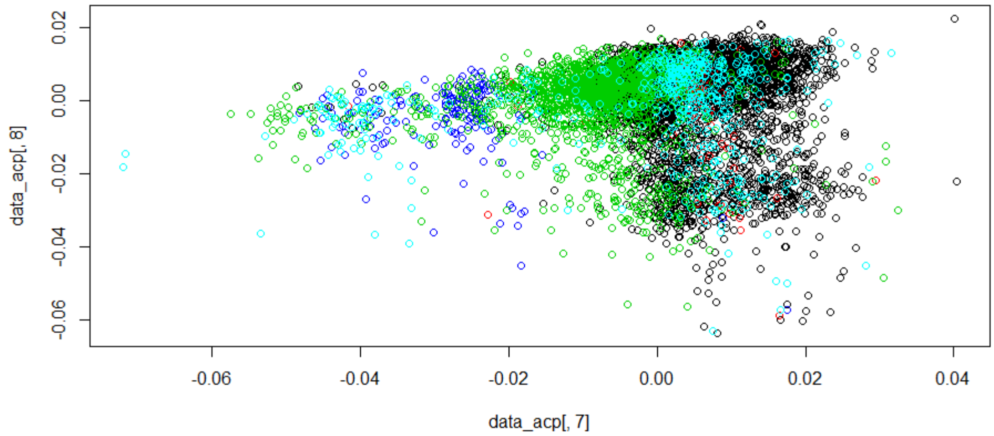

All QC-validated genotyped samples were then imputed for the 1000 Genomes phase 3 version 5 reference panel using the minimac2 software (10,11). A total of 47,109,465 autosomal and 1,345,744 X SNPs were imputed among which 8,945,129 and 207,754, respectively, had imputation criteria > 0.5.

***Utrecht iPSC reprogramming, maintenance, and differentiation***

**iPSC reprogramming**

Dermal fibroblasts were obtained from a 65 years old male donor with no cardiac abnormalities observed on ECG. Genotyping revealed the person is a heterozygous carrier of the R14del variant in the *PLN* gene.

Integration-free iPSC clones were generated from the donor's dermal fibroblasts using the OSKM-dTOMATO lentivirus reprogramming vector(12). Briefly, fibroblasts were transduced with OKSM.dTomato virus and 1:1000 polybrene at day one.

At day 2, medium was refreshed and after 6 days the transduced fibroblasts were seeded on irradiated mouse embryonic fibroblasts (Amsbio) and cultured in human embryonic stem cell (hESC) medium consisting of DMEM-F12 supplemented with 20% knock-out serum replacement, 10 µg/ml penicillin, 10 µg/ml streptomycin, 2 mM L-glutamine, 0.1 mM MEM-NEAA, 0.1 mM β-mercaptoethanol, and 10 ng/ml basic fibroblast growth factor. The hESC medium was refreshed daily. Three clonal iPSC lines were derived from the individual. When iPSC colonies were ready for transfer, the colonies were picked and transferred to 12 wells for individual expansion. Briefly, iPSC lines were maintained in the Essential 8™ (GIBCO) media on tissue culture plates coated with hESC-qualified Matrigel (BD Biosciences) in 5% $CO_2$/5% $O_2$/90% $N_2$ environment at 37°C. All cell lines were free of mycoplasm contamination and tested for their pluripotency. It should be noted that karyotyping revealed missing chromosome 21 and Y and additional chromatin on the short arm of chromosome 16 and 22 as integrative lentivirus confers a higher risk for cytogenetic abnormalities. However, no abnormalities were detected in the GWAS loci: the additional chromatin on the short arm chromosome 22 was distant from the long arm of chromosome 22 where the actual GWAS locus was detected and our 4C analysis probed cis-regulatory regions only.

**Maintenance of human iPSC**

Fibroblast-derived human iPSCs were maintained in Essential8 medium (ThermoFisher Scientific) on 1:400 diluted Matrigel (Corning) and the E8 medium was replaced 7 times per week. Cells were non-enzymatically passaged using 0.5 mM EDTA (Invitrogen, 15575-038) every 4 to 5 days. In brief, cells were washed with PBS, and 1 ml 0.5 mM EDTA-PBS was added per 9.6 $cm^2$ surface area. Cells were incubated at room temperature for 3-5 minutes until cells began to separate uniformly throughout the colonies. PBS-EDTA was removed and iPSC colonies were washed off swiftly using 1 ml E8 medium.

iPSC clumps were passaged in a splitting ratio of 1:10-1:15 routinely at 70-80% confluence. To improve cell survival, split ratio reliability and to reduce selective pressure, ROCK inhibitor was used in the first 24 hrs.

**Differentiation of iPSC into cardiomyocytes**.

To obtain iPSC-derived CMs, hiPSCs (>p30 <p60) were grown to ~90% confluence in 6 wells format. All iPSC lines were maintained in E8 medium for at least five passages before starting cardiac lineage differentiation.

Upon differentiation, the medium was changed to CDM3 medium(13) and supplemented with 6 µM CHIR99021 (Selleck Chemicals), which was replaced with CDM3 containing 2 µM Wnt-C59 (Tocris Bioscience) after 48 hr (day 2). Subsequently, medium changes were performed every other day, and contracting cells were seen from day 7. To metabolically select and purify iPSC-CMs, CDM3 was replaced with purification medium(13) for at least 4 days.

### Circular chromatin conformation capture (4C)-sequencing

#### 4C-Template preparation

4C-chromatin was prepared as described previously(14) with the selection of baits inside LD blocks at each associated locus. In brief, $5x10^6$ cells were used for chromatin preparation per donor. Cells were crosslinked in 2% formaldehyde, lysed in lysis buffer and chromatin was isolated before digestion with DpnII (NEB, #R0543L). After heat inactivation of the restriction enzyme, samples were diluted and ligated by T4 DNA ligase. The second digestion was then performed using CviQI (NEB, #R069S) and inactivated by phenol/chloroform extraction. Finally, the chromatin was diluted, ligated and purified. Digestion and ligation quality were analyzed for proper fragment lengths on 1% agarose gels.

**4C- Primer design**

Primer sequences are listed in **Supplementary table 9**. Primers were designed as was described previously(14). Forward (reading) primers were designed on top of the first restriction enzyme site. The reverse (non-reading) primer was designed close to (if possible, at a maximum 100 bp away from) the second restriction enzyme site.

**4C-seq library preparation**

4C-sequencing library preparation was performed as described previously(14) with minor adaptations (as described here(15)):

the PCR of 4C template was performed with 800 ng to 1,6 µg of 4C template per reaction. Multiple primer pairs were multiplexed in the initial PCR reaction (primer sequences are listed in Supplementary table XX). Primer pairs were pooled according to primer efficiency. PCR products were purified after an initial PCR reaction of 6 cycles (reaction volume = 200 µL) and divided among 8-10 PCR reactions containing single primer pairs for another 26 cycles (reaction volume = 25 µL). Thereafter, PCR products derived from the same cells were pooled in equimolar amounts and a final 6 cycle PCR reaction containing 20 ng of pooled PCR product (reaction volume = 100 µL) was performed with primers that contained sequencing adaptor sequences (Supplementary table XX). All fragments >700 bp were removed using size selection on a 1% agarose gel follow by gel extraction of the selected products (Qiagen, #28704).

**4C-seq data analysis**

Libraries were sequenced using the NextSeq500 platform (Illumina) producing single-end reads of 75 bp. Raw sequenced reads were de-multiplexed based on viewpoint specific primer. Reads were then trimmed to 16 bases and mapped to an in silico generated library of fragends (fragment ends) neighboring all DpnII sites in the human genome (NCBI37/hg19).

No mismatches were allowed during the mapping and the reads mapping to only one fragend were used for further analysis. The interacting domains were identified as described previously(15). In brief: we first calculated the number of covered fragends within a running window of k fragends throughout the whole chromosome where the viewpoint is located. The k was set separately for every viewpoint so it contains on average 20 covered fragends in the area around the viewpoint (+/- 100 kbp). Next, we compared the number of covered fragends in each running window to the random distribution. The windows with a significantly higher number of covered fragends compared to random distribution ($p<10^{-8}$ based on binominal cumulative distribution function; R pbinom) were considered as significant 4C interaction signals.

### References

1. Charron P, Tesson F, Poirier O, Nicaud V, Peuchmaurd M, Tiret L, et al. Identification of a genetic risk factor for idiopathic dilated cardiomyopathy. Involvement of a polymorphism in the endothelin receptor type A gene. CARDIGENE group. Eur Heart J. nov 1999;20(21):1587-91.

2. Duboscq-Bidot L, Charron P, Ruppert V, Fauchier L, Richter A, Tavazzi L, et al. Mutations in the ANKRD1 gene encoding CARP are responsible for human dilated cardiomyopathy. Eur Heart J. sept 2009;30(17):2128-36.

3. Empana J-P, Bean K, Guibout C, Thomas F, Bingham A, Pannier B, et al. Paris Prospective Study III: a study of novel heart rate parameters, baroreflex sensitivity and risk of sudden death. Eur J Epidemiol. nov 2011;26(11):887-92.

4. Holle R, Happich M, Löwel H, Wichmann HE, MONICA/KORA Study Group. KORA--a research platform for population based health research. Gesundheitswesen Bundesverb Arzte Offentlichen Gesundheitsdienstes Ger. août 2005;67 Suppl 1:S19-25.

5. Wichmann H-E, Gieger C, Illig T, MONICA/KORA Study Group. KORA-gen--resource for population genetics, controls and a broad spectrum of disease phenotypes. Gesundheitswesen Bundesverb Arzte Offentlichen Gesundheitsdienstes Ger. août 2005;67 Suppl 1:S26-30.

6. Sammani A, Jansen M, Linschoten M, Bagheri A, de Jonge N, Kirkels H, et al. UNRAVEL: big data analytics research data platform to improve care of patients with cardiomyopathies using routine electronic health records and standardised biobanking. Neth Heart J Mon J Neth Soc Cardiol Neth Heart Found. 27 mai 2019;

7. Angelow A, Schmidt M, Hoffmann W. Towards risk factor assessment in inflammatory dilated cardiomyopathy: the SFB/TR 19 study. Eur J Cardiovasc Prev Rehabil Off J Eur Soc Cardiol Work Groups Epidemiol Prev Card Rehabil Exerc Physiol. oct 2007;14(5):686-93.

8. Völzke H, Alte D, Schmidt CO, Radke D, Lorbeer R, Friedrich N, et al. Cohort profile: the study of health in Pomerania. Int J Epidemiol. avr 2011;40(2):294-307.

9. Purcell S, Neale B, Todd-Brown K, Thomas L, Ferreira MAR, Bender D, et al. PLINK: a tool set for whole-genome association and population-based linkage analyses. Am J Hum Genet. sept 2007;81(3):559-75.

10. Fuchsberger C, Abecasis GR, Hinds DA. minimac2: faster genotype imputation. Bioinforma Oxf Engl. 1 mars 2015;31(5):782-4.

11. Howie B, Fuchsberger C, Stephens M, Marchini J, Abecasis GR. Fast and accurate genotype imputation in genome-wide association studies through pre-phasing. Nat Genet. 22 juill 2012;44(8):955-9.

12. Warlich E, Kuehle J, Cantz T, Brugman MH, Maetzig T, Galla M, et al. Lentiviral Vector Design and Imaging Approaches to Visualize the Early Stages of Cellular Reprogramming. Mol Ther. avr 2011;19(4):782-9.

13. Burridge PW, Matsa E, Shukla P, Lin ZC, Churko JM, Ebert AD, et al. Chemically defined generation of human cardiomyocytes. Nat Methods. août 2014;11(8):855-60.

14. van de Werken HJG, de Vree PJP, Splinter E, Holwerda SJB, Klous P, de Wit E, et al. 4C technology: protocols and data analysis. Methods Enzymol. 2012;513:89-112.

15. Whyte WA, Orlando DA, Hnisz D, Abraham BJ, Lin CY, Kagey MH, et al. Master transcription factors and mediator establish super-enhancers at key cell identity genes. Cell. 11 avr 2013;153(2):307-19.

16. Li H, Durbin R. Fast and accurate short read alignment with Burrows-Wheeler transform. Bioinforma Oxf Engl. 15 juill 2009;25(14):1754-60.

17. Ji H, Jiang H, Ma W, Johnson DS, Myers RM, Wong WH. An integrated software system for analyzing ChIP-chip and ChIP-seq data. Nat Biotechnol. nov 2008;26(11):1293-300.

18. Li H, Handsaker B, Wysoker A, Fennell T, Ruan J, Homer N, et al. The Sequence Alignment/Map format and SAMtools. Bioinforma Oxf Engl. 15 août 2009;25(16):2078-9.

**Supplementary Figure 1. QQ-plot of the discovery GWAS results**

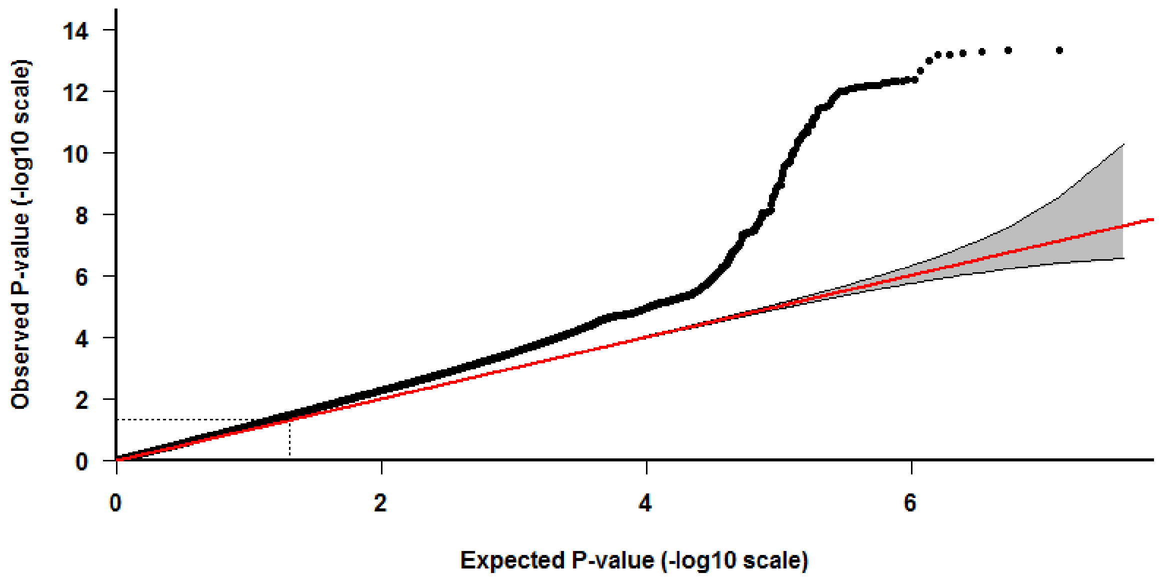

The inflation factor, λ, was evaluated at 1.14

**Supplementary figure 2: Regional association plot at Chromosome 1/*HSPB7* locus**

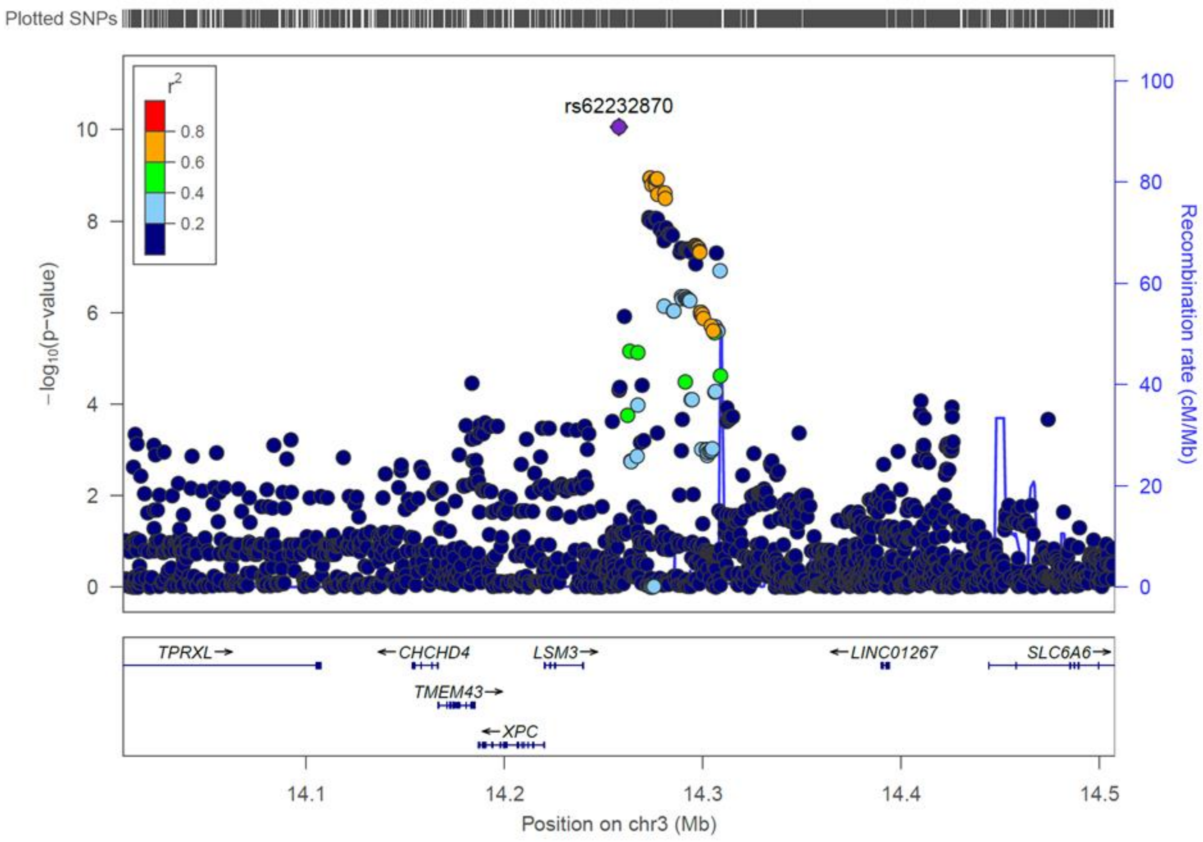

**Supplementary figure 3: Regional association plot at chromosome 3p25.1 locus**

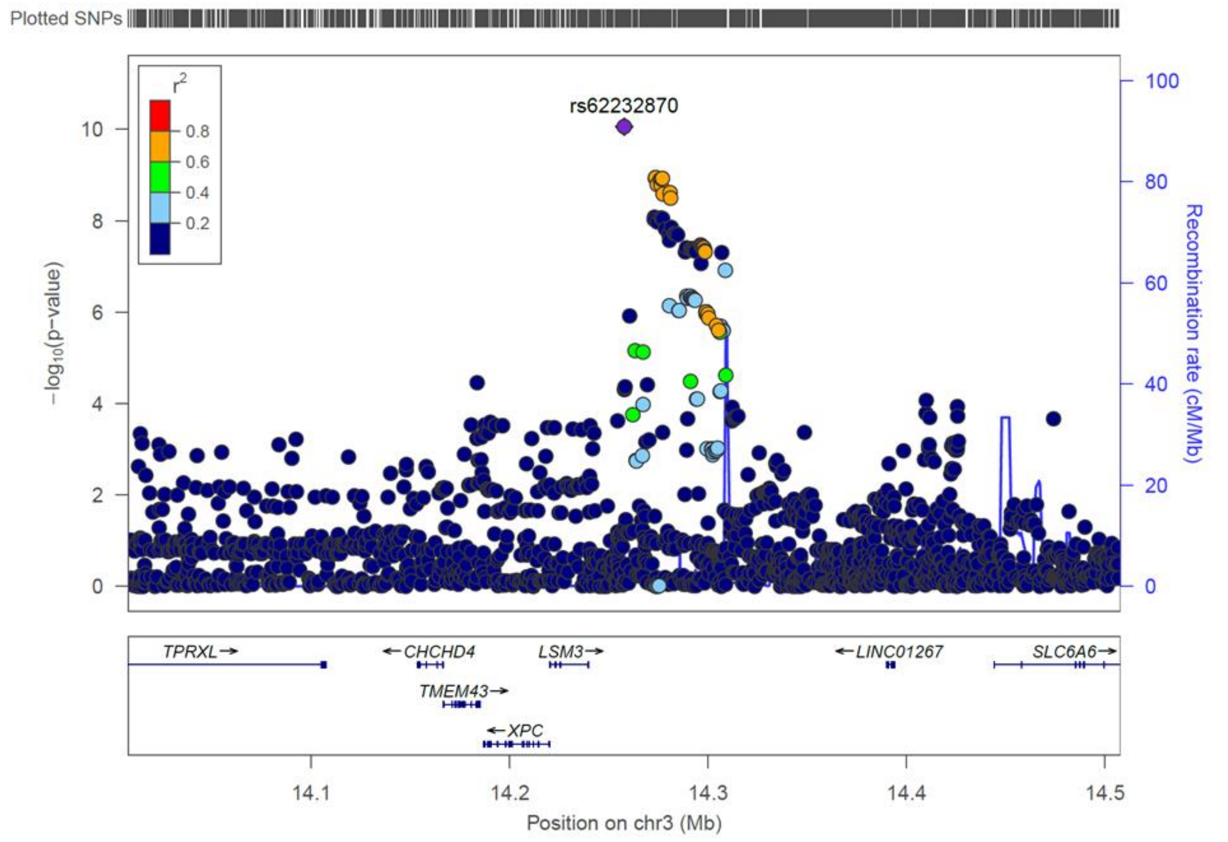

**Supplementary figure 4: Regional association plot at chromosome 10/*BAG3* locus**

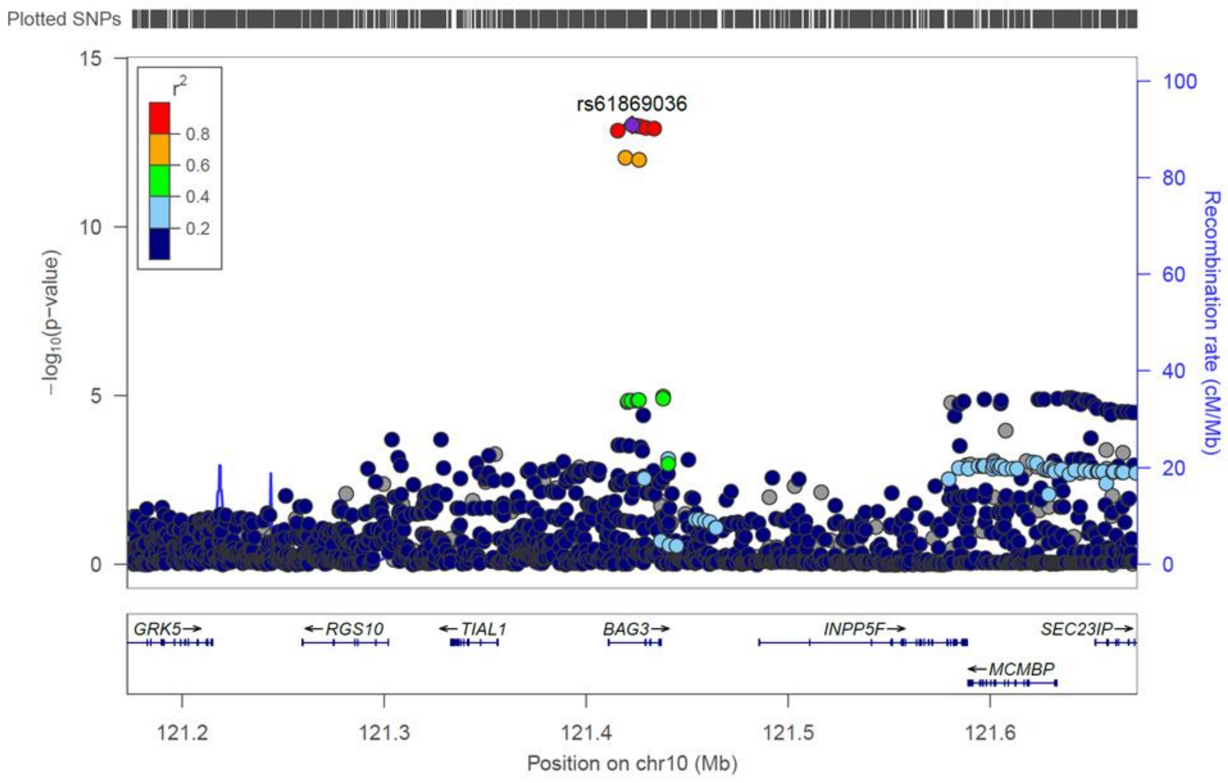

**Supplementary figure 5: Regional association plot at chromosome 16/*PKD1* locus**

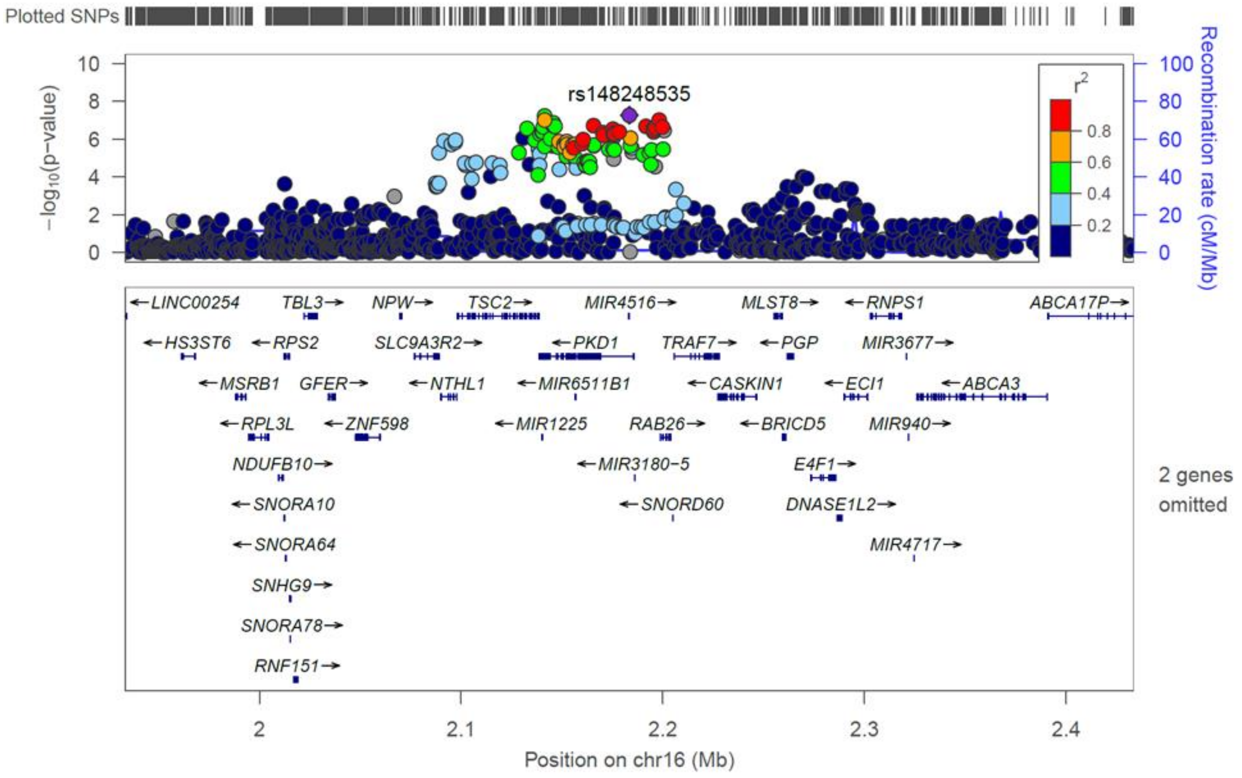

**Supplementary figure 6: Regional association plot at chromosome 22q11.23 /*SMARCB1* locus**

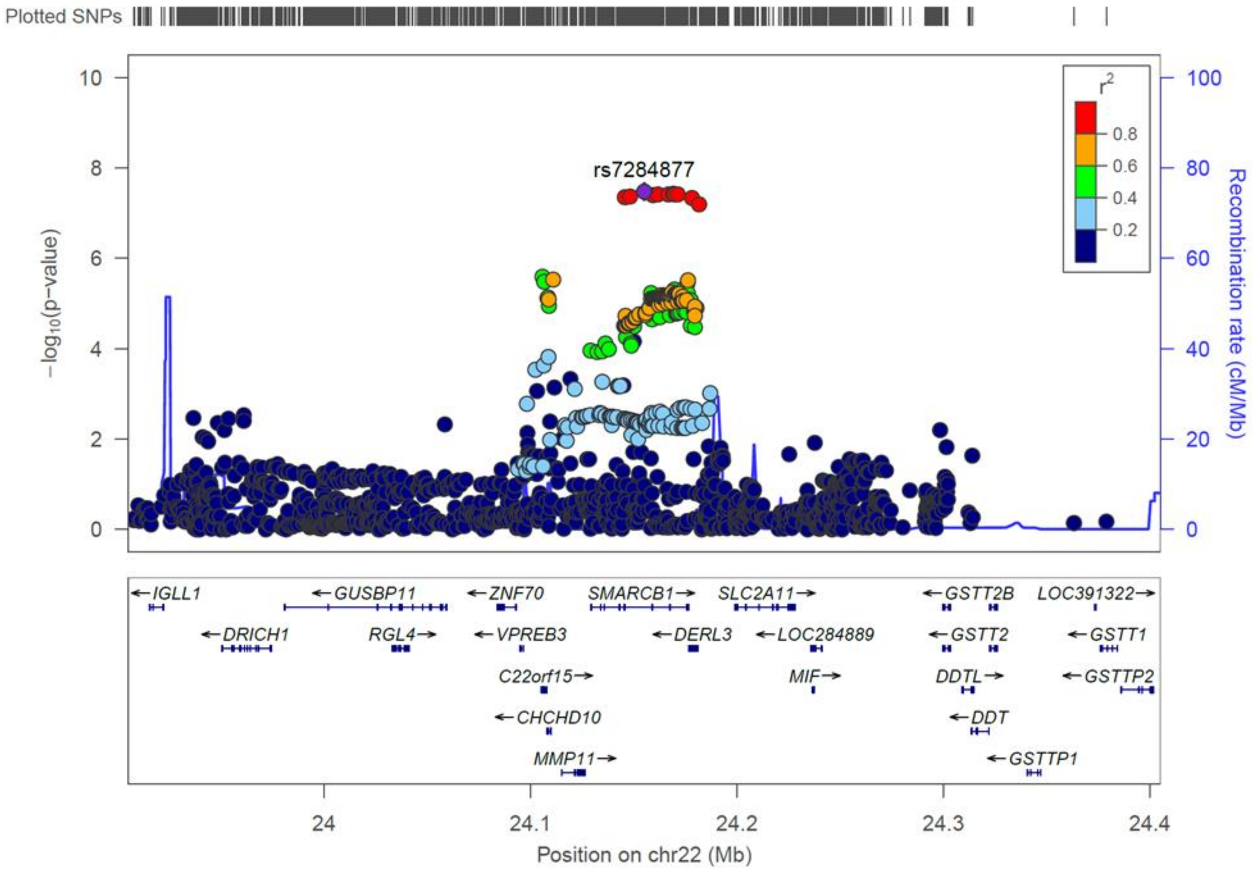

**Supplementary Figure 7. Manhattan plot summarizing the results of the discovery GWAS analysis adjusted on the 5 lead SNPs (rs10927886, rs2234962, rs62232870, rs2519236 and rs7284877)**

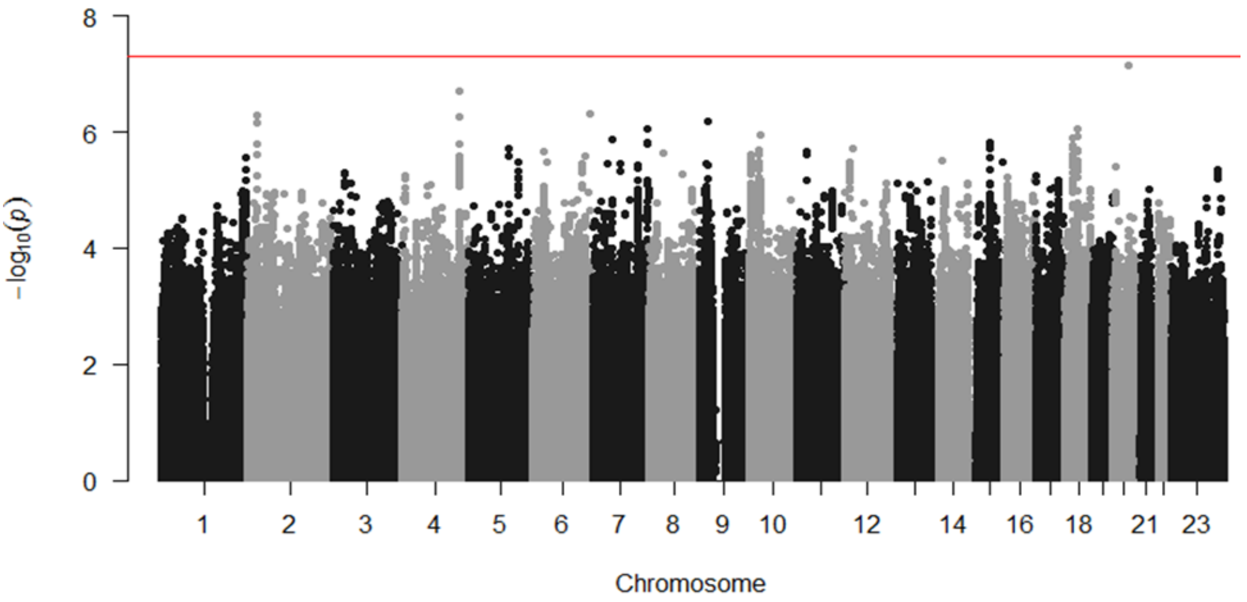

**Supplementary Figure 8. QQ-plot derived from the discovery GWAS association results after adjustment on the 5 lead SNPs (rs10927886, rs2234962, rs62232870, rs2519236 and rs7284877)**

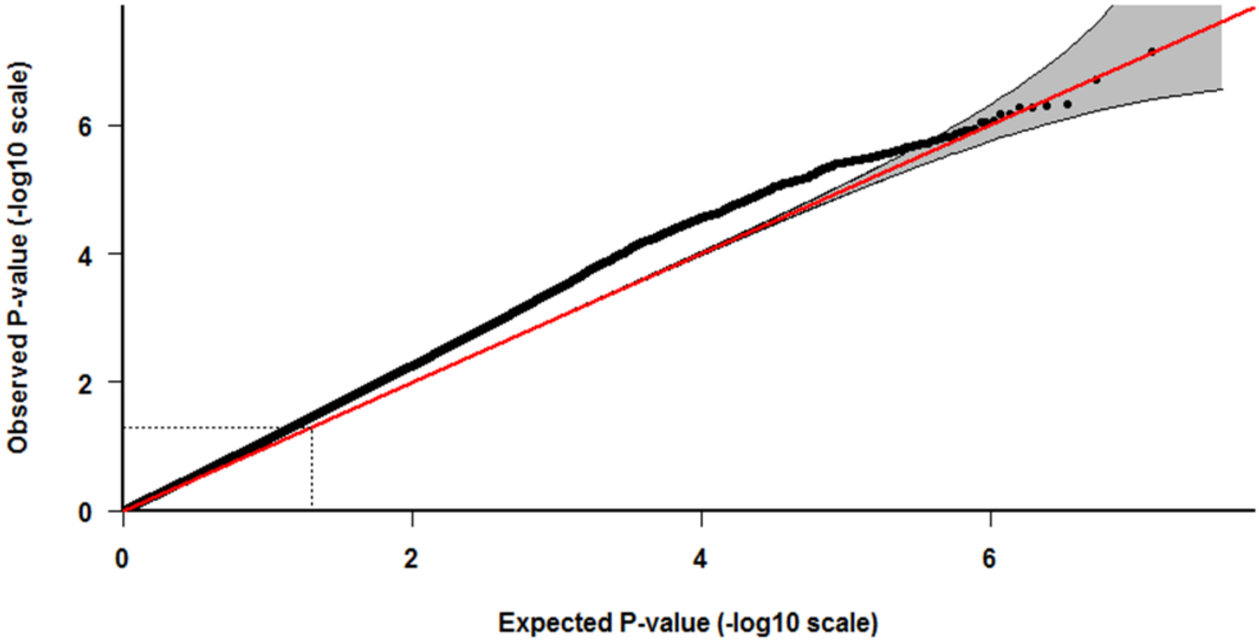

**Supplementary Figure 9. Weighted* Genetic Risk Score grouped by quintile for the 6,980 individuals of the discovery cohort and associated OR taking the quintile 40-60% (score 1.55) as reference.**

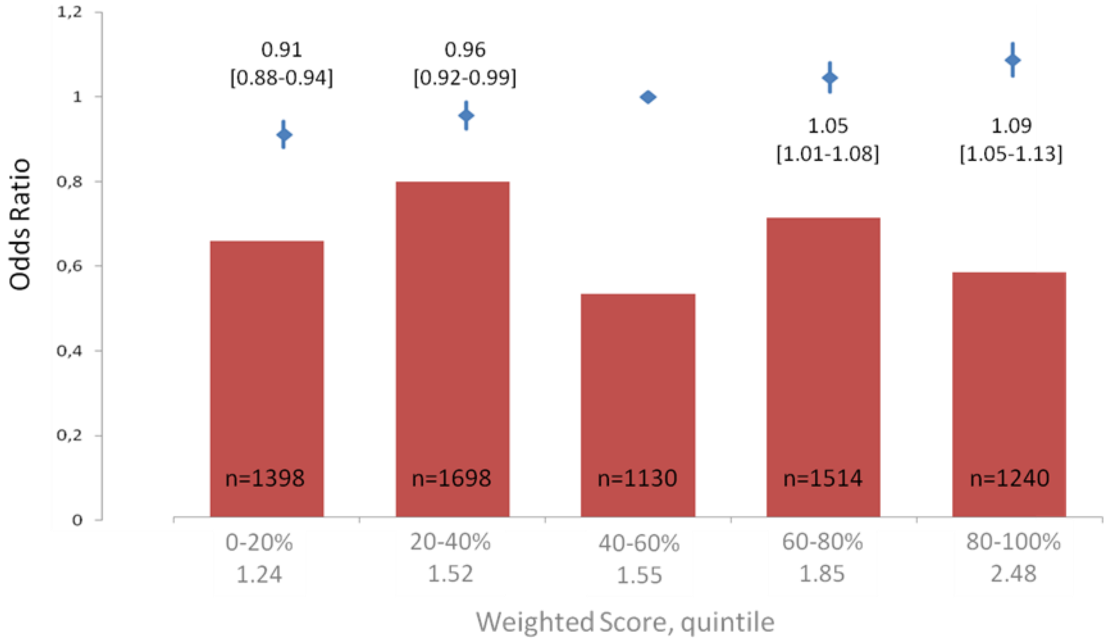

*Score of each SNP weighted by the beta value of this SNP in the sub meta-analysis of the two replication cohorts

**Supplementary Figure 10. Regulation of *SMARCB1* expression by rs7284877 lead SNP (22_24155111_G_C_b37) in Heart Left Ventricle and Atrial Appendage (GTEx)**

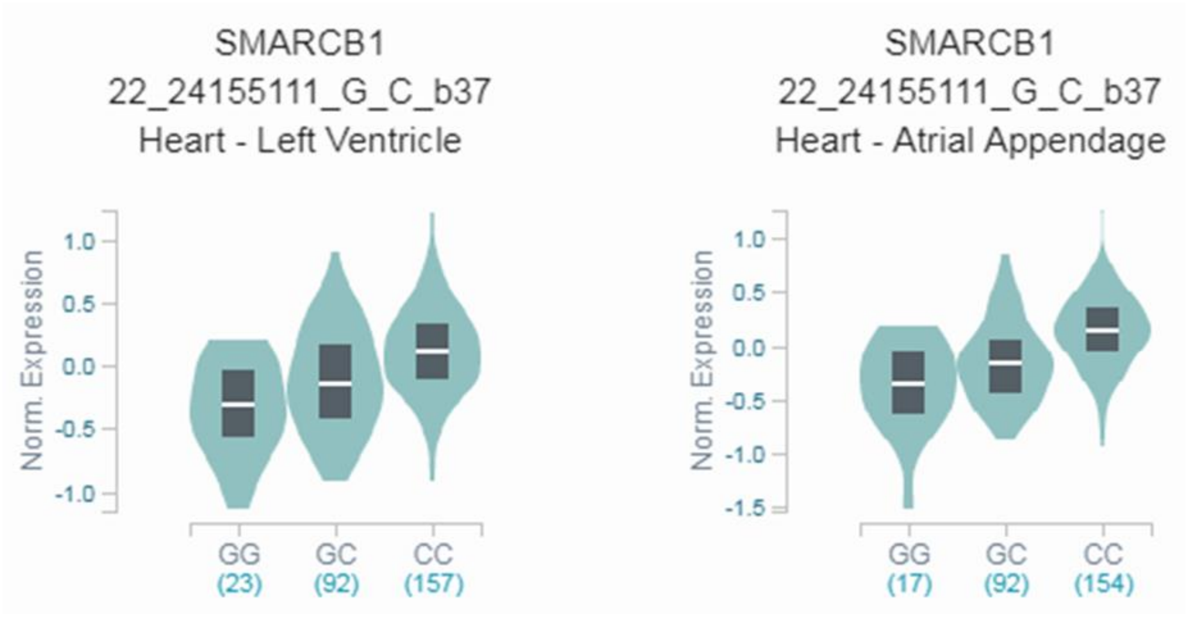

$p = 2.8\ 10^{-12}$ $p = 1.5\ 10^{-13}$

**Supplementary Figure 11. Regulation of *SLC6A* expression by rs62231957 (chr3_14256355_G_A_b38), $r^2$=0.7 with rs62232870 lead SNP, in Heart Atrial Appendage (GTEx)**

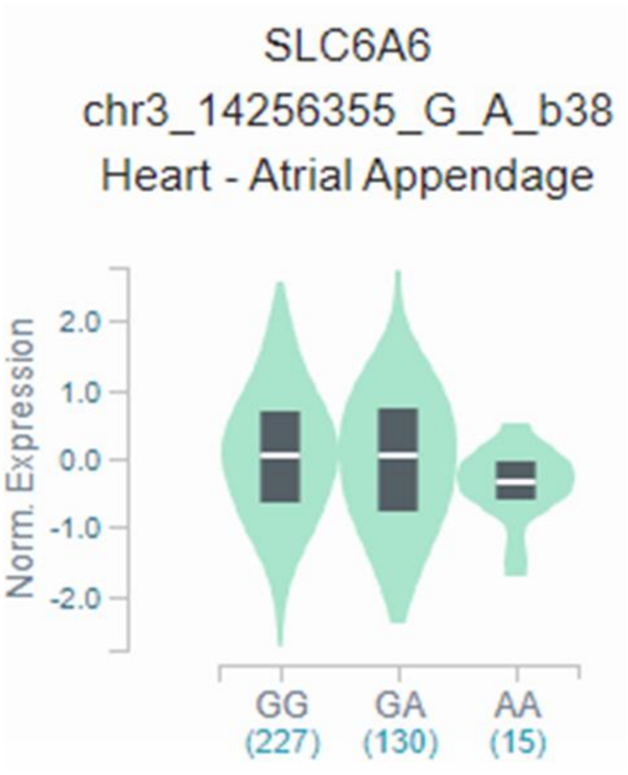

p = $1.9.10^{-5}$

**Supplementary Table 1: Genotyping/imputation information for each participating study.** This excel table is given in a supplemental excel file.

**Supplementary Table 2: Replication of the rs62232870-rs4684185 derived haplotypes association of with DCM**

| | | **Discovery Cohort** | | | **Dutch Replication Study** | | | **German Replication Study** | | | **Combined** |
|---|---|---|---|---|---|---|---|---|---|---|---|
| | | Haplotype frequency | | Haplotypic OR [95%CI] | Haplotype frequency | | Haplotypic OR [95%CI] | Haplotype frequency | | Haplotypic OR [95%CI] | Haplotypic OR [95%CI] |
| rs62232870 | rs4684185 | Controls | Cases | | Controls | Cases | | Controls | Cases | | |
| G | C | 0.475 | 0.481 | Reference | 0.472 | 0.506 | Reference | 0.503 | 0.484 | Reference | Reference |
| G | T | 0.317 | 0.264 | 0.819 [0.755 - 0.889] | 0.326 | 0.251 | 0.724 [0.533 -0.982] | 0.306 | 0.273 | 0.929 [0.743 - 1.162] | 0.844 [0.766 – 0.888] $p = 3.39\ 10^{-7}$ |
| A | C | 0.205 | 0.253 | 1.216 [1.114 - 1.327] | 0.201 | 0.244 | 1.157 [0.810 -1.655] | 0.188 | 0.240 | 1.310 [1.029 - 1.669] | 1.223 [1.129 – 1.325] $p = 8.82\ 10^{-7}$ |
| A | T | 0.003 | 0.002 | NA | 0.002 | 0.001 | NA | 0.002 | 0.002 | NA | NA |

**Supplementary Table 3A: Unweighted Genetic Score for the 6,980 individuals of the discovery cohort and associated OR taking the score 5 as reference.**

| | Discovery Cohort | | | Dutch Replication Study | | | German Replication Study | | |
|---|---|---|---|---|---|---|---|---|---|
| Risk allele number | Size | p-value | Odds Ratio [±CI] | Size | p-value | Odds Ratio [±CI] | Size | p-value | Odds Ratio [±CI] |
| 1 | 22 | 0.012 | 0.79 [0.66-0.95] | 2 | 0.29 | 0.77 [0.48-1.25] | 2 | 0.97 | NA |
| 2 | 121 | $3.20\ 10^{-05}$ | 0.85 [0.79-0.92] | 18 | 0.59 | 0.96 [0.81-1.13] | 23 | 0.07 | 0.42 [0.17-1.07] |
| 3 | 594 | $4.98\ 10^{-05}$ | 0.92 [0.89-0.96] | 51 | 0.08 | 0.91 [0.82-1.01] | 70 | 0.03 | 0.55 [0.32-0.96] |
| 4 | 1638 | $1.56\ 10^{-05}$ | 0.94 [0.91-0.97] | 145 | 0.37 | 1.03 [0.96-1.11] | 203 | 0.09 | 0.73 [0.50-1.05] |
| 5 | 2262 | Reference | Reference | 217 | Reference | Reference | 260 | Reference | Reference |
| 6 | 1682 | $6.07\ 10^{-06}$ | 1.07 [1.04-1.10] | 161 | 0.27 | 1.04 [0.97-1.12] | 240 | 0.37 | 1.18 [0.83-1.68] |
| 7 | 598 | $8.00\ 10^{-10}$ | 1.13 [1.09-1.18] | 56 | $7.33\ 10^{-04}$ | 1.19 [1.08-1.32] | 67 | 0.11 | 1.57 [0.90-2.73] |
| 8 | 63 | $2.05\ 10^{-05}$ | 1.27 [1.14-1.42] | 3 | 0.53 | 0.88 [0.59-1.31] | 13 | 0.03 | 5.42 [1.16-25.3] |

**Supplementary Table 3B: Weighted Genetic Score for the 6,980 individuals of the discovery cohort and associated OR taking the score 1.6 as reference.**

Weighing was realized taking the sub-meta-analysis (two replication cohorts) beta value

| | Discovery Cohort | | | Dutch Replication Study | | | German Replication Study | | |
|---|---|---|---|---|---|---|---|---|---|
| Weighted Score | Size | p-value | Odds Ratio [±CI] | Size | p-value | Odds Ratio [±CI] | Size | p-value | Odds Ratio [±CI] |
| 0,4 | 22 | 0.024 | 0.81 [0.67-0.97] | 2 | 0.28 | 0.76 [0.47-1.24] | 2 | 0.97 | NA |
| 0,8 | 121 | $7.12\ 10^{-04}$ | 0.87 [0.80-0.94] | 18 | 0.51 | 0.95 [0.80-1.12] | 23 | 0.12 | 0.49 [0.20-1.20] |
| 1,2 | 793 | $5.56\ 10^{-04}$ | 0.94 [0.91-0.97] | 71 | 0.18 | 0.94 [0.86-1.03] | 97 | 0.12 | 0.70 [0.44-1.10] |
| 1,6 | 3694 | Reference | Reference | 342 | Reference | Reference | 435 | Reference | Reference |
| 2 | 1689 | $1.67\ 10^{-11}$ | 1.09 [1.06-1.12] | 161 | 0.38 | 1.03 [0.96-1.10] | 241 | 0.06 | 1.36 [0.99-1.87] |
| 2,4 | 598 | $3.48\ 10^{-14}$ | 1.16 [1.12-1.20] | 56 | $1.00\ 10^{-03}$ | 1.18 [1.07-1.30] | 67 | 0.03 | 1.80 [1.06-3.05] |
| 2,8 | 63 | $2.82\ 10^{-06}$ | 1.30 [1.16-1.45] | 3 | 0.50 | 0.87 [0.59-1.30] | 13 | 0.02 | 6.22 [1.34-28.6] |

**Supplementary Table 4: SNPs at chromosome 3p25.1 and 22q11.23 associated regions with p-values ≤ 5 x 10-8 and/or in LD (r2 > 0.7) with the lead SNP** *The indel variant haven't been analyzed in the imputed data to prevent putative imputation mistakes

| RS_number | Coordinates (GRCh37.p13) | Alleles | MAF | Distance (kb) | D' | $r^2$ | Correlated_Alleles | p-value discovery GWAS* |
|---|---|---|---|---|---|---|---|---|
| rs62232870 | chr3:14257709 | (G/A) | 0.1869 | 0 | 1.0 | 1.0 | G=G,A=A | 8,74E-11 |
| rs113608926 | chr3:14296105 | (-/TG) | 0.2121 | 38396 | 0.8971 | 0.6869 | G=-,A=TG | 3,402E-08 |
| rs13090226 | chr3:14261998 | (T/C) | 0.3131 | 4289 | 1.0 | 0.5041 | G=T,A=C | 1,75E-04 |
| rs12630964 | chr3:14263206 | (G/A) | 0.298 | 5497 | 1.0 | 0.5414 | G=G,A=A | 6,86E-06 |
| rs12639212 | chr3:14267211 | (A/G) | 0.3081 | 9502 | 1.0 | 0.5161 | G=A,A=G | 7,48E-06 |
| rs55834511 | chr3:14273414 | (G/C) | 0.2071 | 15705 | 0.8977 | 0.7092 | G=G,A=C | 1,175E-09 |
| rs55647831 | chr3:14273464 | (C/T) | 0.2071 | 15755 | 0.8977 | 0.7092 | G=C,A=T | 1,153E-09 |
| rs56242556 | chr3:14273472 | (C/T) | 0.2071 | 15763 | 0.8977 | 0.7092 | G=C,A=T | 1,152E-09 |
| rs56281979 | chr3:14274293 | (C/T) | 0.2071 | 16584 | 0.8977 | 0.7092 | G=C,A=T | 1,567E-09 |
| rs6442433 | chr3:14275759 | (C/G) | 0.2071 | 18050 | 0.8977 | 0.7092 | G=C,A=G | 1,245E-09 |
| rs7612736 | chr3:14276238 | (G/A) | 0.2071 | 18529 | 0.8977 | 0.7092 | G=G,A=A | 1,629E-09 |
| rs62231949 | chr3:14276494 | (G/A) | 0.2071 | 18785 | 0.8977 | 0.7092 | G=G,A=A | 1,203E-09 |
| rs17226476 | chr3:14276982 | (G/A) | 0.2071 | 19273 | 0.8977 | 0.7092 | G=G,A=A | 1,192E-09 |
| rs17305149 | chr3:14277297 | (A/G) | 0.2071 | 19588 | 0.8977 | 0.7092 | G=A,A=G | 2,595E-09 |
| rs4685090 | chr3:14280870 | (G/T) | 0.2071 | 23161 | 0.8977 | 0.7092 | G=G,A=T | 2,408E-09 |
| rs62231950 | chr3:14281051 | (G/T) | 0.2071 | 23342 | 0.8977 | 0.7092 | G=G,A=T | 3,137E-09 |
| rs62231953 | chr3:14296180 | (T/A) | 0.2121 | 38471 | 0.8971 | 0.6869 | G=T,A=A | 3,407E-08 |
| rs62231954 | chr3:14296311 | (C/T) | 0.2121 | 38602 | 0.8971 | 0.6869 | G=C,A=T | 3,431E-08 |
| rs116103636 | chr3:14296781 | (G/A) | 0.2121 | 39072 | 0.8971 | 0.6869 | G=G,A=A | 3,455E-08 |
| rs62231955 | chr3:14297159 | (C/T) | 0.2121 | 39450 | 0.8971 | 0.6869 | G=C,A=T | 3,778E-08 |
| rs62231956 | chr3:14297290 | (G/A) | 0.2121 | 39581 | 0.8971 | 0.6869 | G=G,A=A | 3,712E-08 |
| rs9843704 | chr3:14297304 | (C/T) | 0.2121 | 39595 | 0.8971 | 0.6869 | G=C,A=T | 4,053E-08 |

| | | | | | | | | |
|---|---|---|---|---|---|---|---|---|
| rs62231957 | chr3:14297855 | (G/A) | 0.2121 | 40146 | 0.8971 | 0.6869 | G=G,A=A | 3,871E-08 |
| rs62231958 | chr3:14298347 | (G/C) | 0.2121 | 40638 | 0.8971 | 0.6869 | G=G,A=C | 4,46E-08 |
| rs62231959 | chr3:14298407 | (A/G) | 0.2121 | 40698 | 0.8971 | 0.6869 | G=A,A=G | 4,51E-08 |
| rs55824951 | chr3:14298510 | (G/C) | 0.2121 | 40801 | 0.8971 | 0.6869 | G=G,A=C | 4,779E-08 |
| rs4684185 | chr3:14272914 | (C/T) | 0.3485 | 0 | 1.0 | 1.0 | C=C,T=T | 8,386E-09 |
| rs73028853 | chr3:14273117 | (C/T) | 0.3485 | 203 | 1.0 | 1.0 | C=C,T=T | 9,528E-09 |
| rs6807275 | chr3:14274451 | (G/A) | 0.3485 | 1537 | 1.0 | 1.0 | C=G,T=A | 1,056E-08 |
| rs11709201 | chr3:14276263 | (A/G) | 0.3485 | 3349 | 1.0 | 1.0 | C=A,T=G | 8,716E-09 |
| rs55691578 | chr3:14277060 | (A/C) | 0.3485 | 4146 | 1.0 | 1.0 | C=A,T=C | 8,82E-09 |
| rs900173 | chr3:14278376 | (T/C) | 0.3485 | 5462 | 1.0 | 1.0 | C=T,T=C | 1,502E-08 |
| rs900175 | chr3:14280232 | (T/G) | 0.3485 | 7318 | 1.0 | 1.0 | C=T,T=G | 1,957E-08 |
| rs900174 | chr3:14280136 | (G/A) | 0.3485 | 7222 | 1.0 | 1.0 | C=G,T=A | 1,965E-08 |
| rs4684186 | chr3:14280458 | (G/T) | 0.3485 | 7544 | 1.0 | 1.0 | C=G,T=T | 2,62E-08 |
| rs4685091 | chr3:14281431 | (A/G) | 0.3485 | 8517 | 1.0 | 1.0 | C=A,T=G | 1,384E-08 |
| rs6442434 | chr3:14282894 | (A/G) | 0.3485 | 9980 | 1.0 | 1.0 | C=A,T=G | 1,903E-08 |
| rs7640329 | chr3:14283776 | (A/G) | 0.3485 | 10862 | 1.0 | 1.0 | C=A,T=G | 1,902E-08 |
| rs7618619 | chr3:14283861 | (G/T) | 0.3485 | 10947 | 1.0 | 1.0 | C=G,T=T | 1,903E-08 |
| rs73031103 | chr3:14284662 | (A/G) | 0.3485 | 11748 | 1.0 | 1.0 | C=A,T=G | 2,021E-08 |
| rs6442435 | chr3:14288558 | (C/T) | 0.3485 | 15644 | 1.0 | 1.0 | C=C,T=T | 4,774E-08 |
| rs73031110 | chr3:14289114 | (G/A) | 0.3485 | 16200 | 1.0 | 1.0 | C=G,T=A | 3,918E-08 |
| rs200775399 | chr3:14291003 | (TTTT/-) | 0.3485 | 18089 | 1.0 | 1.0 | C=TTTT,T=- | NA |
| rs1871854 | chr3:14291091 | (C/T) | 0.3485 | 18177 | 1.0 | 1.0 | C=C,T=T | 4,15E-08 |
| rs11715059 | chr3:14292138 | (A/G) | 0.3485 | 19224 | 1.0 | 1.0 | C=A,T=G | 3,968E-08 |
| rs11710541 | chr3:14291679 | (T/C) | 0.3485 | 18765 | 1.0 | 1.0 | C=T,T=C | 3,985E-08 |
| rs55735564 | chr3:14292498 | (C/T) | 0.3485 | 19584 | 1.0 | 1.0 | C=C,T=T | 4,324E-08 |
| rs12630973 | chr3:14293372 | (G/T) | 0.3485 | 20458 | 1.0 | 1.0 | C=G,T=T | 4,47E-08 |
| rs12631029 | chr3:14293622 | (G/A) | 0.3485 | 20708 | 1.0 | 1.0 | C=G,T=A | 4,583E-08 |
| rs73028848 | chr3:14272760 | (A/G) | 0.3535 | -154 | 1.0 | 0.9781 | C=A,T=G | 9,235E-09 |

| | | | | | | | | |
|---|---|---|---|---|---|---|---|---|
| rs73028849 | chr3:14272766 | (G/C) | 0.3535 | -148 | 1.0 | 0.9781 | C=G,T=C | 9,528E-09 |
| rs113308768 | chr3:14296182 | (-/TG) | 0.3333 | 23268 | 1.0 | 0.9348 | C=TG,T=- | NA |
| rs59625485 | chr3:14296437 | (G/T) | 0.3333 | 23523 | 1.0 | 0.9348 | C=T,T=G | 8,433E-08 |
| rs10865722 | chr3:14306782 | (G/T) | 0.3283 | 33868 | 1.0 | 0.9137 | C=G,T=T | 5,002E-08 |
| rs11721007 | chr3:14260417 | (G/A) | 0.3586 | -12497 | 0.9548 | 0.8722 | C=G,T=A | 1,02E-06 |
| rs7284877 | chr22:24155111 | (G/C) | 0.1465 | 0 | 1.0 | 1.0 | G=G,C=C | 3,346E-08 |
| rs9608201 | chr22:24166788 | (G/A) | 0.1465 | 11677 | 1.0 | 1.0 | G=G,C=A | 3,831E-08 |
| rs2186370 | chr22:24171305 | (A/G) | 0.1465 | 16194 | 1.0 | 1.0 | G=A,C=G | 3,838E-08 |
| rs2267039 | chr22:24169751 | (C/T) | 0.1465 | 14640 | 1.0 | 1.0 | G=C,C=T | 3,853E-08 |
| rs5760054 | chr22:24161717 | (C/T) | 0.1465 | 6606 | 1.0 | 1.0 | G=C,C=T | 3,878E-08 |
| rs2070458 | chr22:24159307 | (A/T) | 0.1465 | 4196 | 1.0 | 1.0 | G=A,C=T | 3,91E-08 |
| rs2267032 | chr22:24148273 | (A/G) | 0.1465 | -6838 | 1.0 | 1.0 | G=A,C=G | 4,23E-08 |
| rs5760032 | chr22:24145727 | (C/T) | 0.1465 | -9384 | 1.0 | 1.0 | G=C,C=T | 4,362E-08 |
| rs5760061 | chr22:24178279 | (G/A) | 0.1465 | 23168 | 1.0 | 1.0 | G=G,C=A | 4,568E-08 |
| rs60463265 | chr22:24160324 | (GTGTGTGTGTGT/-) | 0.1364 | 5213 | 1.0 | 0.9201 | G=GTGTGTGTGTGT,C=- | NA |
| rs2267038 | chr22:24169196 | (G/C) | 0.1364 | 14085 | 1.0 | 0.9201 | G=G,C=C | 3,742E-08 |
| rs6003909 | chr22:24181652 | (A/G) | 0.1263 | 26541 | 0.9531 | 0.7651 | G=A,C=G | 6,263E-08 |
| rs131445 | chr22:24111044 | (C/A) | 0.1616 | -44067 | 0.9177 | 0.7497 | G=C,C=A | 2,90E-06 |
| rs80135378 | chr22:24162363 | (A/T) | 0.1111 | 7252 | 1.0 | 0.7284 | G=T,C=A | 1,08E-05 |
| rs9612464 | chr22:24163666 | (G/C) | 0.1111 | 8555 | 1.0 | 0.7284 | G=C,C=G | 1,00E-05 |
| rs17003930 | chr22:24146055 | (T/G) | 0.1111 | -9056 | 1.0 | 0.7284 | G=G,C=T | 1,89E-05 |
| rs6003895 | chr22:24166540 | (T/C) | 0.1111 | 11429 | 1.0 | 0.7284 | G=C,C=T | 9,62E-06 |
| rs8140489 | chr22:24168335 | (A/G) | 0.1111 | 13224 | 1.0 | 0.7284 | G=G,C=A | 9,50E-06 |
| rs6003898 | chr22:24171404 | (A/G) | 0.1111 | 16293 | 1.0 | 0.7284 | G=G,C=A | 9,19E-06 |
| rs6003900 | chr22:24173022 | (G/A) | 0.1111 | 17911 | 1.0 | 0.7284 | G=A,C=G | 8,84E-06 |
| rs6003904 | chr22:24173394 | (A/G) | 0.1111 | 18283 | 1.0 | 0.7284 | G=G,C=A | 8,81E-06 |
| rs6003905 | chr22:24173633 | (G/A) | 0.1111 | 18522 | 1.0 | 0.7284 | G=A,C=G | 8,08E-06 |

| | | | | | | | | |
|---|---|---|---|---|---|---|---|---|
| rs17003983 | chr22:24173718 | (A/G) | 0.1111 | 18607 | 1.0 | 0.7284 | G=G,C=A | 7,10E-06 |
| rs8138435 | chr22:24173919 | (T/G) | 0.1111 | 18808 | 1.0 | 0.7284 | G=G,C=T | 8,79E-06 |
| rs8138827 | chr22:24174145 | (T/C) | 0.1111 | 19034 | 1.0 | 0.7284 | G=C,C=T | 8,79E-06 |
| rs2267045 | chr22:24175583 | (A/C) | 0.1111 | 20472 | 1.0 | 0.7284 | G=C,C=A | 8,49E-06 |
| rs1128127 | chr22:24179132 | (G/A) | 0.1111 | 24021 | 1.0 | 0.7284 | G=A,C=G | 1,44E-05 |
| rs9612485 | chr22:24179502 | (G/A) | 0.1111 | 24391 | 1.0 | 0.7284 | G=A,C=G | 1,46E-05 |
| rs11703971 | chr22:24179648 | (T/C) | 0.1111 | 24537 | 1.0 | 0.7284 | G=C,C=T | 1,84E-05 |

**Supplementary Table 5: List of positional candidate genes in the associated loci**

| Chromosome | Gene Symbol | Gene Name | Position (start-end) hg19 | Functions |
|---|---|---|---|---|
| 3 | *CHCHD4* | Coiled-coil-helix-coiled-coil-helix domain containing 4 | 14153577-14166371, complement | Component of human mitochondrial intermembrane space with import and/or protein folding function. Protein familywith 6 highly conserved cysteine residues constituting a -CXC-CX(9)C-CX(9)C- motif in C terminus |
| 3 | *TMEM43* | Transmembrane protein 43 | 14166440-14185180 | Transmembrane protein participating to nuclear envelope structure maintenance. Defects in this gene are implied in familial arrhythmogenic right ventricular cardiomyopathy type 5 (ARVC5), an inherited disorder, often involving both ventricles, characterized by ventricular tachycardia, heart failure, sudden cardiac death, and fibrofatty replacement of cardiomyocytes. |
| 3 | *XPC* | XPC complex subunit, DNA damage recognition and repair factor | 14186647-14220172, complement | Key component of the XPC complex, which plays an important role in the early steps of global genome nucleotide excision repair (NER). The encoded protein is important for damage sensing and DNA binding, and shows a preference for single-stranded DNA. Mutations in this gene or some other NER components can result in Xeroderma pigmentosum |
| 3 | *LSM3* | U4/U6-U5 snRNP complex subunit LSM3 | 14220228-14239869 | Plays a role in pre-mRNA splicing as component of the U4/U6-U5 tri-snRNP complex involved in spliceosome assembly and as component of the precatalytic spliceosome (spliceosome B complex) |
| 3 | *SLC6A6* | Solute carrier family 6 member 6 | 14444076-14530857 | Multi-pass membrane protein, sodium-dependent taurine and beta-alanine transporter |
| 3 | *GRIP2* | Glutamate receptor interacting protein 2 | 14530396-14583588, complement | May play a role as a localized scaffold for a multiprotein signaling complex assembly and as mediator of its binding partners' trafficking at specific subcellular location in neurons |
| 22 | *RAB36* | RAB36, member RAS oncogene family | 23487513-23506531 | Protein with GTPase activity that may have a role in protein transport and vesicular trafficking. Implied in rhabdoid and bladder cancer |
| 22 | *BCR* | BCR activator of RhoGEF and GTPase | 23522402-23660224 | Encode a protein with two opposing regulatory activities toward small GTP-binding proteins. The C-terminus is a GTPase-activating protein (GAP) domain which stimulates GTP hydrolysis by RAC1, RAC2 and CDC42, accelerates GTP hydrolysis rate of RAC1 or |

| | | | | |
|---|---|---|---|---|
| | | | | CDC42 thus leading to down-regulation of the active GTP-bound form. The central Dbl homology (DH) domain functions as guanine nucleotide exchange factor (GEF) that modulates the GTPases CDC42, RHOA and RAC1 and promotes the conversion of CDC42, RHOA and RAC1 from the GDP-bound to the GTP-bound form. The amino terminus contains an intrinsic kinase activity. Functions as an important negative regulator of neuronal RAC1 activity (By similarity). Regulates macrophage functions such as CSF1-directed motility and phagocytosis through the modulation of RAC1 activity |
| 22 | *FBXW4P1* | F-box and WD repeat domain containing 4 pseudogene 1 | 23604954-23607192 | Pseudogen |
| 22 | *ZDHHC8P1* | ZDHHC8 pseudogene 1 | 23732792-23744799, complement | Pseudogen |
| 22 | *IGLL1* | Immunoglobulin lambda like polypeptide 1 | 23915312-23922619, complement | Member of the immunoglobulin gene superfamily that encodes one of the surrogate light chain subunits. Mutations can result in B cell deficiency and agammaglobulinemia, an autosomal recessive disease |
| 22 | *DRICH1* | Aspartate rich 1 | 23950639-23974508, complement | DRICH1 aspartate rich 1 |
| 22 | *RGL4* | Ral guanine nucleotide dissociation stimulator like 4 | 24033048-24041444 | Encodes a protein similar to guanine nucleotide exchange factor Ral guanine dissociation stimulator. Increased expression leads to translocation of the encoded protein to the cell membrane which can activate several pathways, including the Ras-Raf-MEK-ERK cascade |
| 22 | *ZNF70* | Zinc finger protein 70 | 24083772-24093279, complement | May have a role in transcriptional regulation |
| 22 | *VPREB3* | V-set pre-B cell surrogate light chain 3 | 24094930-24096630, complement | The encoded protein associates with the Ig-mu chain to form a molecular complex expressed on the surface of pre-B-cells. May have a role in B-cell maturation and assembly of the pre-B cell receptor (pre-BCR). Expression of this gene has been observed in some lymphomas |

| | | | | |
|---|---|---|---|---|
| 22 | *C22orf15* | Chromosome 22 open reading frame 15 | 24102622-24108050 | |
| 22 | *CHCHD10* | Coiled-coil-helix-coiled-coil-helix domain containing 10 | 24108021-24110141, complement | Encodes a mitochondrial protein enriched at cristae junctions in the intermembrane space. May play a role in cristae morphology maintenance or oxidative phosphorylation. Mutations in this gene cause frontotemporal dementia and/or amyotrophic lateral sclerosis-2. |
| 22 | *MMP11* | Matrix metallopeptidase 11 | 24115036-24126503 | The encoded enzyme is intracellularly activated by furin within the constitutive secretory pathway. In contrast to other MMP's, this enzyme cleaves alpha 1-proteinase inhibitor but weakly degrades structural proteins of the extracellular matrix. May play an important role in the progression of epithelial malignancies. |
| 22 | *SMARCB1* | SWI/SNF related, matrix associated, actin dependent regulator of chromatin, subfamily b, member 1 | 24129150-24176705 | Core component of the BAF (hSWI/SNF) complex. This ATP-dependent chromatin-remodeling complex plays important roles in cell proliferation and differentiation, in cellular antiviral activities and inhibition of tumor formation. This gene is a tumor suppressor. Mutations have been associated with malignant rhabdoid tumors |
| 22 | *DERL3* | Derlin 3 | 24176690-24181430, complement | Functional component of endoplasmic reticulum-associated degradation (ERAD) for misfolded lumenal glycoprotein |
| 22 | *SLC2A11* | Solute carrier family 2 member 11 | 24199059-24227738 | Glucose and fructose transporter |
| 22 | *MIF* | Macrophage migration inhibitory factor | 24236565-24237409 | Encodes a pro-inflammatory cytokine involved in cell-mediated immunity, immunoregulation, innate immune response to bacterial pathogens and inflammation. It plays a role in the regulation of macrophage function in host defense through the suppression of anti-inflammatory effects of glucocorticoids. This lymphokine and the JAB1 protein form a complex in the cytosol near the peripheral plasma membrane, which may indicate an additional role in integrin signaling pathways. |
| 22 | *GSTT2* | Glutathione S-transferase theta 2 (gene/pseudogene) | 24322291-24326106 | Encodes a member of a protein superfamily that catalyzes conjugation of reduced glutathione to a variety of electrophilic and hydrophobic compounds that have a sulfatase activity. May play a role in human carcinogenesis |

| | | | | |
|---|---|---|---|---|
| 22 | *DDTL* | D-dopachrome tautomerase like | 24308652-24314748 | May have lyase activity |
| 22 | *DDT* | D-dopachrome tautomerase | 24313554-24322019, complement | Converts D-dopachrome into 5,6-dihydroxyindole. Related to migration inhibitory factor (MIF) in terms of sequence, enzyme activity, and gene structure |
| 22 | *GSTT1* | Glutathione S-transferase theta 1 | 24376135-24384284, complement | Encodes a member of a protein superfamily that catalyzes conjugation of reduced glutathione to a variety of electrophilic and hydrophobic compounds. *GSTT1* is haplotype-specific and absent from 38% of the population. May play a role in human carcinogenesis. |
| 22 | *CABIN1* | Calcineurin-binding protein 1 | 24407642-24574596 | The protein encoded by this gene binds specifically to the activated form of calcineurin and inhibits calcineurin-mediated signal transduction which may negatively regulates T-cell receptor signaling. May act as a negative regulator of p53/TP53 by keeping p53 in an inactive state on chromatin at promoters of a subset of it's target genes. The encoded protein is found in the nucleus and contains a leucine zipper domain as well as several PEST motifs, sequences which confer targeted degradation to those proteins which contain them. May be required for replication-independent chromatin assembly. |

**Supplementary figure 6. 4C interaction p-Value at chr3p25.1.** DNA sequence segments location is based on GRCh37.p13

| Chromosome | Start position | End position | P-value |
|---|---|---|---|
| 3 | 14292587 | 14309550 | 2.98E-63 |
| 3 | 14293744 | 14309662 | 2.98E-63 |
| 3 | 14295215 | 14310198 | 2.98E-63 |
| 3 | 14292061 | 14309220 | 5.33E-60 |
| 3 | 14295267 | 14311311 | 5.33E-60 |
| 3 | 14295565 | 14311423 | 5.33E-60 |
| 3 | 14295703 | 14311895 | 5.33E-60 |
| 3 | 14291542 | 14309107 | 6.14E-57 |
| 3 | 14296297 | 14312015 | 6.14E-57 |
| 3 | 14297247 | 14312031 | 6.14E-57 |
| 3 | 14297965 | 14313663 | 6.14E-57 |
| 3 | 14298063 | 14313844 | 6.14E-57 |
| 3 | 14497311 | 14510622 | 6.14E-57 |
| 3 | 14497399 | 14513390 | 6.14E-57 |
| 3 | 14498474 | 14515805 | 6.14E-57 |
| 3 | 14498955 | 14516319 | 6.14E-57 |
| 3 | 14259756 | 14284820 | 5.13E-54 |
| 3 | 14260538 | 14286068 | 5.13E-54 |
| 3 | 14290828 | 14308459 | 5.13E-54 |
| 3 | 14298084 | 14314339 | 5.13E-54 |
| 3 | 14298275 | 14316846 | 5.13E-54 |
| 3 | 14299064 | 14316897 | 5.13E-54 |
| 3 | 14433210 | 14449429 | 5.13E-54 |
| 3 | 14434236 | 14449767 | 5.13E-54 |
| 3 | 14435021 | 14450226 | 5.13E-54 |

| | | | |
|---|---|---|---|
| 3 | 14435057 | 14450260 | 5.13E-54 |
| 3 | 14435093 | 14450278 | 5.13E-54 |
| 3 | 14494787 | 14507146 | 5.13E-54 |
| 3 | 14495113 | 14507198 | 5.13E-54 |
| 3 | 14495397 | 14507927 | 5.13E-54 |
| 3 | 14495955 | 14509111 | 5.13E-54 |
| 3 | 14495961 | 14510286 | 5.13E-54 |
| 3 | 14496662 | 14510445 | 5.13E-54 |
| 3 | 14497670 | 14514285 | 5.13E-54 |
| 3 | 14497858 | 14515690 | 5.13E-54 |
| 3 | 14500584 | 14517013 | 5.13E-54 |
| 3 | 14218000 | 14243878 | 3.31E-51 |
| 3 | 14219560 | 14244281 | 3.31E-51 |
| 3 | 14219752 | 14244829 | 3.31E-51 |
| 3 | 14220807 | 14245013 | 3.31E-51 |
| 3 | 14220987 | 14245333 | 3.31E-51 |
| 3 | 14221266 | 14245461 | 3.31E-51 |
| 3 | 14255977 | 14279672 | 3.31E-51 |
| 3 | 14259277 | 14284514 | 3.31E-51 |
| 3 | 14261092 | 14286336 | 3.31E-51 |
| 3 | 14262602 | 14286375 | 3.31E-51 |
| 3 | 14263562 | 14286430 | 3.31E-51 |
| 3 | 14264775 | 14286684 | 3.31E-51 |
| 3 | 14268256 | 14288199 | 3.31E-51 |
| 3 | 14270675 | 14288406 | 3.31E-51 |
| 3 | 14272823 | 14288454 | 3.31E-51 |
| 3 | 14273650 | 14288462 | 3.31E-51 |
| 3 | 14283914 | 14299461 | 3.31E-51 |
| 3 | 14283985 | 14300336 | 3.31E-51 |

| | | | |
|---|---|---|---|
| 3 | 14284514 | 14300394 | 3.31E-51 |
| 3 | 14286375 | 14301802 | 3.31E-51 |
| 3 | 14286430 | 14302145 | 3.31E-51 |
| 3 | 14286684 | 14302465 | 3.31E-51 |
| 3 | 14288199 | 14303544 | 3.31E-51 |
| 3 | 14288406 | 14303578 | 3.31E-51 |
| 3 | 14288454 | 14304203 | 3.31E-51 |
| 3 | 14288462 | 14304383 | 3.31E-51 |
| 3 | 14290179 | 14305059 | 3.31E-51 |
| 3 | 14299261 | 14317050 | 3.31E-51 |
| 3 | 14299461 | 14317208 | 3.31E-51 |
| 3 | 14300336 | 14318013 | 3.31E-51 |
| 3 | 14300394 | 14318094 | 3.31E-51 |
| 3 | 14301237 | 14321484 | 3.31E-51 |
| 3 | 14301488 | 14322371 | 3.31E-51 |
| 3 | 14301802 | 14324700 | 3.31E-51 |
| 3 | 14302145 | 14324790 | 3.31E-51 |
| 3 | 14302465 | 14326120 | 3.31E-51 |
| 3 | 14303544 | 14326297 | 3.31E-51 |
| 3 | 14433071 | 14449255 | 3.31E-51 |
| 3 | 14434698 | 14449777 | 3.31E-51 |
| 3 | 14434986 | 14449801 | 3.31E-51 |
| 3 | 14435206 | 14450722 | 3.31E-51 |
| 3 | 14435308 | 14450788 | 3.31E-51 |
| 3 | 14435540 | 14451167 | 3.31E-51 |
| 3 | 14439915 | 14456821 | 3.31E-51 |
| 3 | 14440048 | 14457032 | 3.31E-51 |
| 3 | 14491219 | 14504701 | 3.31E-51 |
| 3 | 14491943 | 14506115 | 3.31E-51 |

| | | | |
|---|---|---|---|
| 3 | 14492591 | 14506340 | 3.31E-51 |
| 3 | 14492689 | 14506633 | 3.31E-51 |
| 3 | 14494220 | 14506828 | 3.31E-51 |
| 3 | 14500718 | 14517358 | 3.31E-51 |
| 3 | 14217876 | 14243445 | 1.72E-48 |
| 3 | 14221486 | 14245957 | 1.72E-48 |
| 3 | 14223813 | 14247929 | 1.72E-48 |
| 3 | 14223875 | 14247991 | 1.72E-48 |
| 3 | 14223942 | 14248515 | 1.72E-48 |
| 3 | 14254767 | 14279666 | 1.72E-48 |
| 3 | 14256333 | 14280084 | 1.72E-48 |
| 3 | 14258228 | 14280757 | 1.72E-48 |
| 3 | 14258442 | 14280962 | 1.72E-48 |
| 3 | 14258613 | 14283985 | 1.72E-48 |
| 3 | 14273954 | 14290179 | 1.72E-48 |
| 3 | 14275196 | 14290828 | 1.72E-48 |
| 3 | 14280757 | 14298275 | 1.72E-48 |
| 3 | 14280962 | 14299064 | 1.72E-48 |
| 3 | 14282103 | 14299261 | 1.72E-48 |
| 3 | 14284820 | 14301196 | 1.72E-48 |
| 3 | 14286068 | 14301237 | 1.72E-48 |
| 3 | 14286336 | 14301488 | 1.72E-48 |
| 3 | 14301196 | 14320376 | 1.72E-48 |
| 3 | 14303578 | 14326951 | 1.72E-48 |
| 3 | 14304203 | 14327123 | 1.72E-48 |
| 3 | 14304383 | 14327551 | 1.72E-48 |
| 3 | 14432630 | 14449082 | 1.72E-48 |
| 3 | 14437423 | 14451228 | 1.72E-48 |
| 3 | 14437731 | 14451307 | 1.72E-48 |

| | | | |
|---|---|---|---|
| 3 | 14439698 | 14452731 | 1.72E-48 |
| 3 | 14439903 | 14454302 | 1.72E-48 |
| 3 | 14441915 | 14457166 | 1.72E-48 |
| 3 | 14442019 | 14457355 | 1.72E-48 |
| 3 | 14490342 | 14504121 | 1.72E-48 |
| 3 | 14500883 | 14517799 | 1.72E-48 |
| 3 | 14500957 | 14518072 | 1.72E-48 |
| 3 | 14501472 | 14518295 | 1.72E-48 |
| 3 | 14502202 | 14518576 | 1.72E-48 |
| 3 | 14217397 | 14243120 | 7.36E-46 |
| 3 | 14217848 | 14243201 | 7.36E-46 |
| 3 | 14222030 | 14246035 | 7.36E-46 |
| 3 | 14224669 | 14248839 | 7.36E-46 |
| 3 | 14225463 | 14248920 | 7.36E-46 |
| 3 | 14225902 | 14248949 | 7.36E-46 |
| 3 | 14237794 | 14250063 | 7.36E-46 |
| 3 | 14238152 | 14251195 | 7.36E-46 |
| 3 | 14239029 | 14251386 | 7.36E-46 |
| 3 | 14254704 | 14279430 | 7.36E-46 |
| 3 | 14258495 | 14282103 | 7.36E-46 |
| 3 | 14258607 | 14283914 | 7.36E-46 |
| 3 | 14275310 | 14291542 | 7.36E-46 |
| 3 | 14275461 | 14292587 | 7.36E-46 |
| 3 | 14275752 | 14293744 | 7.36E-46 |
| 3 | 14276003 | 14295215 | 7.36E-46 |
| 3 | 14276631 | 14295267 | 7.36E-46 |
| 3 | 14277749 | 14295565 | 7.36E-46 |
| 3 | 14278910 | 14295703 | 7.36E-46 |
| 3 | 14279278 | 14296297 | 7.36E-46 |

| | | | |
|---|---|---|---|
| 3 | 14279430 | 14297247 | 7.36E-46 |
| 3 | 14279666 | 14297965 | 7.36E-46 |
| 3 | 14279672 | 14298063 | 7.36E-46 |
| 3 | 14280084 | 14298084 | 7.36E-46 |
| 3 | 14305059 | 14328530 | 7.36E-46 |
| 3 | 14432352 | 14448799 | 7.36E-46 |
| 3 | 14442536 | 14457883 | 7.36E-46 |
| 3 | 14490276 | 14503666 | 7.36E-46 |
| 3 | 14502560 | 14518736 | 7.36E-46 |
| 3 | 14503017 | 14519114 | 7.36E-46 |
| 3 | 14503194 | 14520241 | 7.36E-46 |
| 3 | 14503305 | 14520506 | 7.36E-46 |
| 3 | 14503325 | 14520675 | 7.36E-46 |
| 3 | 14503334 | 14521007 | 7.36E-46 |
| 3 | 14215640 | 14242075 | 2.65E-43 |
| 3 | 14215870 | 14242617 | 2.65E-43 |
| 3 | 14227273 | 14249215 | 2.65E-43 |
| 3 | 14227405 | 14249235 | 2.65E-43 |
| 3 | 14236158 | 14249364 | 2.65E-43 |
| 3 | 14236388 | 14249556 | 2.65E-43 |
| 3 | 14238557 | 14251203 | 2.65E-43 |
| 3 | 14239065 | 14252172 | 2.65E-43 |
| 3 | 14254232 | 14279278 | 2.65E-43 |
| 3 | 14275320 | 14292061 | 2.65E-43 |
| 3 | 14308459 | 14328787 | 2.65E-43 |
| 3 | 14309107 | 14328961 | 2.65E-43 |
| 3 | 14432337 | 14447220 | 2.65E-43 |
| 3 | 14442796 | 14457922 | 2.65E-43 |
| 3 | 14489451 | 14503334 | 2.65E-43 |

| | | | |
|---|---|---|---|
| 3 | 14503666 | 14521390 | 2.65E-43 |
| 3 | 14504121 | 14523027 | 2.65E-43 |
| 3 | 14504701 | 14523427 | 2.65E-43 |
| 3 | 14506115 | 14523824 | 2.65E-43 |
| 3 | 14518072 | 14533650 | 2.65E-43 |
| 3 | 14214841 | 14240668 | 8.16E-41 |
| 3 | 14239416 | 14252346 | 8.16E-41 |
| 3 | 14254075 | 14278910 | 8.16E-41 |
| 3 | 14309220 | 14329305 | 8.16E-41 |
| 3 | 14431441 | 14445893 | 8.16E-41 |
| 3 | 14443186 | 14458126 | 8.16E-41 |
| 3 | 14476829 | 14495961 | 8.16E-41 |
| 3 | 14487343 | 14503325 | 8.16E-41 |
| 3 | 14506340 | 14523856 | 8.16E-41 |
| 3 | 14517799 | 14531797 | 8.16E-41 |
| 3 | 14518295 | 14534150 | 8.16E-41 |
| 3 | 14518736 | 14535131 | 8.16E-41 |
| 3 | 14519114 | 14535207 | 8.16E-41 |
| 3 | 14161810 | 14175877 | 2.16E-38 |
| 3 | 14214259 | 14240620 | 2.16E-38 |
| 3 | 14240620 | 14254075 | 2.16E-38 |
| 3 | 14252346 | 14277749 | 2.16E-38 |
| 3 | 14309550 | 14329777 | 2.16E-38 |
| 3 | 14309662 | 14330096 | 2.16E-38 |
| 3 | 14310198 | 14331204 | 2.16E-38 |
| 3 | 14430594 | 14445721 | 2.16E-38 |
| 3 | 14443461 | 14458231 | 2.16E-38 |
| 3 | 14443664 | 14459347 | 2.16E-38 |
| 3 | 14445461 | 14459412 | 2.16E-38 |

| | | | |
|---|---|---|---|
| 3 | 14472508 | 14494787 | 2.16E-38 |
| 3 | 14472627 | 14495113 | 2.16E-38 |
| 3 | 14473108 | 14495397 | 2.16E-38 |
| 3 | 14476763 | 14495955 | 2.16E-38 |
| 3 | 14477339 | 14496662 | 2.16E-38 |
| 3 | 14486382 | 14503194 | 2.16E-38 |
| 3 | 14487292 | 14503305 | 2.16E-38 |
| 3 | 14506633 | 14523959 | 2.16E-38 |
| 3 | 14506828 | 14524922 | 2.16E-38 |
| 3 | 14507146 | 14525165 | 2.16E-38 |
| 3 | 14507198 | 14525749 | 2.16E-38 |
| 3 | 14507927 | 14526246 | 2.16E-38 |
| 3 | 14509111 | 14526893 | 2.16E-38 |
| 3 | 14510286 | 14526912 | 2.16E-38 |
| 3 | 14510445 | 14527345 | 2.16E-38 |
| 3 | 14510622 | 14528020 | 2.16E-38 |
| 3 | 14514285 | 14529605 | 2.16E-38 |
| 3 | 14515690 | 14529659 | 2.16E-38 |
| 3 | 14515805 | 14529813 | 2.16E-38 |
| 3 | 14516319 | 14530279 | 2.16E-38 |
| 3 | 14517013 | 14530650 | 2.16E-38 |
| 3 | 14517358 | 14531146 | 2.16E-38 |
| 3 | 14518576 | 14534477 | 2.16E-38 |
| 3 | 14520241 | 14535254 | 2.16E-38 |
| 3 | 14524922 | 14543183 | 2.16E-38 |
| 3 | 14525165 | 14543456 | 2.16E-38 |
| 3 | 14583210 | 14597251 | 2.16E-38 |
| 3 | 14583525 | 14598749 | 2.16E-38 |
| 3 | 14159619 | 14173473 | 4.99E-36 |

| | | | |
|---|---|---|---|
| 3 | 14159629 | 14173604 | 4.99E-36 |
| 3 | 14159816 | 14173760 | 4.99E-36 |
| 3 | 14160692 | 14173949 | 4.99E-36 |
| 3 | 14160814 | 14174210 | 4.99E-36 |
| 3 | 14160884 | 14175413 | 4.99E-36 |
| 3 | 14162967 | 14176443 | 4.99E-36 |
| 3 | 14211865 | 14238152 | 4.99E-36 |
| 3 | 14213888 | 14239416 | 4.99E-36 |
| 3 | 14240668 | 14254232 | 4.99E-36 |
| 3 | 14248920 | 14270675 | 4.99E-36 |
| 3 | 14248949 | 14272823 | 4.99E-36 |
| 3 | 14249215 | 14273650 | 4.99E-36 |
| 3 | 14249235 | 14273954 | 4.99E-36 |
| 3 | 14249364 | 14275196 | 4.99E-36 |
| 3 | 14249556 | 14275310 | 4.99E-36 |
| 3 | 14251386 | 14276003 | 4.99E-36 |
| 3 | 14252172 | 14276631 | 4.99E-36 |
| 3 | 14311311 | 14332360 | 4.99E-36 |
| 3 | 14311423 | 14333990 | 4.99E-36 |
| 3 | 14321484 | 14344758 | 4.99E-36 |
| 3 | 14430584 | 14445673 | 4.99E-36 |
| 3 | 14445673 | 14459607 | 4.99E-36 |
| 3 | 14464815 | 14481483 | 4.99E-36 |
| 3 | 14465425 | 14482124 | 4.99E-36 |
| 3 | 14466483 | 14484339 | 4.99E-36 |
| 3 | 14467574 | 14485286 | 4.99E-36 |
| 3 | 14467741 | 14486382 | 4.99E-36 |
| 3 | 14467902 | 14487292 | 4.99E-36 |
| 3 | 14468833 | 14487343 | 4.99E-36 |

| | | | |
|---|---|---|---|
| 3 | 14471561 | 14492689 | 4.99E-36 |
| 3 | 14472401 | 14494220 | 4.99E-36 |
| 3 | 14478152 | 14497311 | 4.99E-36 |
| 3 | 14478481 | 14497399 | 4.99E-36 |
| 3 | 14479071 | 14497670 | 4.99E-36 |
| 3 | 14480186 | 14500584 | 4.99E-36 |
| 3 | 14480961 | 14500718 | 4.99E-36 |
| 3 | 14481483 | 14500883 | 4.99E-36 |
| 3 | 14482124 | 14500957 | 4.99E-36 |
| 3 | 14483781 | 14502202 | 4.99E-36 |
| 3 | 14484339 | 14502560 | 4.99E-36 |
| 3 | 14485286 | 14503017 | 4.99E-36 |
| 3 | 14513390 | 14529083 | 4.99E-36 |
| 3 | 14520506 | 14539024 | 4.99E-36 |
| 3 | 14520675 | 14539178 | 4.99E-36 |
| 3 | 14523959 | 14542539 | 4.99E-36 |
| 3 | 14525749 | 14543483 | 4.99E-36 |
| 3 | 14526246 | 14544257 | 4.99E-36 |
| 3 | 14582656 | 14596484 | 4.99E-36 |
| 3 | 14583614 | 14599339 | 4.99E-36 |
| 3 | 14585061 | 14599434 | 4.99E-36 |
| 3 | 14585078 | 14600215 | 4.99E-36 |
| 3 | 14585379 | 14600481 | 4.99E-36 |
| 3 | 13930894 | 13945508 | 1.01E-33 |
| 3 | 13931245 | 13945630 | 1.01E-33 |
| 3 | 14159552 | 14173314 | 1.01E-33 |
| 3 | 14163195 | 14177023 | 1.01E-33 |
| 3 | 14164466 | 14178033 | 1.01E-33 |
| 3 | 14191658 | 14206306 | 1.01E-33 |

| | | | |
|---|---|---|---|
| 3 | 14191946 | 14206773 | 1.01E-33 |
| 3 | 14210441 | 14227405 | 1.01E-33 |
| 3 | 14210537 | 14236158 | 1.01E-33 |
| 3 | 14210950 | 14236388 | 1.01E-33 |
| 3 | 14211062 | 14237794 | 1.01E-33 |
| 3 | 14212244 | 14238557 | 1.01E-33 |
| 3 | 14212295 | 14239029 | 1.01E-33 |
| 3 | 14213396 | 14239065 | 1.01E-33 |
| 3 | 14242075 | 14254704 | 1.01E-33 |
| 3 | 14247929 | 14262602 | 1.01E-33 |
| 3 | 14247991 | 14263562 | 1.01E-33 |
| 3 | 14248515 | 14264775 | 1.01E-33 |
| 3 | 14248839 | 14268256 | 1.01E-33 |
| 3 | 14250063 | 14275320 | 1.01E-33 |
| 3 | 14251195 | 14275461 | 1.01E-33 |
| 3 | 14251203 | 14275752 | 1.01E-33 |
| 3 | 14311895 | 14336257 | 1.01E-33 |
| 3 | 14312031 | 14338619 | 1.01E-33 |
| 3 | 14317208 | 14342521 | 1.01E-33 |
| 3 | 14318013 | 14342734 | 1.01E-33 |
| 3 | 14318094 | 14343342 | 1.01E-33 |
| 3 | 14320376 | 14343703 | 1.01E-33 |
| 3 | 14322371 | 14345107 | 1.01E-33 |
| 3 | 14370499 | 14390222 | 1.01E-33 |
| 3 | 14373320 | 14391146 | 1.01E-33 |
| 3 | 14378027 | 14394287 | 1.01E-33 |
| 3 | 14378667 | 14395078 | 1.01E-33 |
| 3 | 14379483 | 14395209 | 1.01E-33 |
| 3 | 14382424 | 14395955 | 1.01E-33 |

| | | | |
|---|---|---|---|
| 3 | 14382444 | 14397397 | 1.01E-33 |
| 3 | 14382816 | 14397500 | 1.01E-33 |
| 3 | 14383217 | 14399032 | 1.01E-33 |
| 3 | 14383295 | 14399256 | 1.01E-33 |
| 3 | 14393255 | 14408386 | 1.01E-33 |
| 3 | 14394057 | 14409629 | 1.01E-33 |
| 3 | 14430578 | 14445461 | 1.01E-33 |
| 3 | 14445721 | 14461440 | 1.01E-33 |
| 3 | 14445893 | 14462233 | 1.01E-33 |
| 3 | 14463994 | 14480961 | 1.01E-33 |
| 3 | 14465691 | 14483344 | 1.01E-33 |
| 3 | 14466104 | 14483781 | 1.01E-33 |
| 3 | 14469044 | 14489451 | 1.01E-33 |
| 3 | 14469469 | 14490276 | 1.01E-33 |
| 3 | 14469645 | 14490342 | 1.01E-33 |
| 3 | 14469821 | 14491219 | 1.01E-33 |
| 3 | 14470922 | 14491943 | 1.01E-33 |
| 3 | 14471313 | 14492591 | 1.01E-33 |
| 3 | 14479216 | 14497858 | 1.01E-33 |
| 3 | 14479758 | 14498474 | 1.01E-33 |
| 3 | 14480140 | 14498955 | 1.01E-33 |
| 3 | 14483344 | 14501472 | 1.01E-33 |
| 3 | 14521007 | 14539247 | 1.01E-33 |
| 3 | 14521390 | 14539651 | 1.01E-33 |
| 3 | 14523027 | 14539963 | 1.01E-33 |
| 3 | 14523427 | 14540606 | 1.01E-33 |
| 3 | 14523824 | 14540770 | 1.01E-33 |
| 3 | 14523856 | 14542054 | 1.01E-33 |
| 3 | 14526893 | 14544511 | 1.01E-33 |

| | | | |
|---|---|---|---|
| 3 | 14526912 | 14544522 | 1.01E-33 |
| 3 | 14527345 | 14544674 | 1.01E-33 |
| 3 | 14540606 | 14561297 | 1.01E-33 |
| 3 | 14580137 | 14593616 | 1.01E-33 |
| 3 | 14581245 | 14594409 | 1.01E-33 |
| 3 | 14581728 | 14595811 | 1.01E-33 |
| 3 | 14582506 | 14595904 | 1.01E-33 |
| 3 | 14585690 | 14600722 | 1.01E-33 |
| 3 | 13930492 | 13945483 | 1.79E-31 |
| 3 | 13931553 | 13945912 | 1.79E-31 |
| 3 | 13931562 | 13945944 | 1.79E-31 |
| 3 | 13931996 | 13946058 | 1.79E-31 |
| 3 | 13932641 | 13946259 | 1.79E-31 |
| 3 | 13932646 | 13946539 | 1.79E-31 |
| 3 | 14159351 | 14173092 | 1.79E-31 |
| 3 | 14163506 | 14177485 | 1.79E-31 |
| 3 | 14163733 | 14177632 | 1.79E-31 |
| 3 | 14164500 | 14178099 | 1.79E-31 |
| 3 | 14164662 | 14179219 | 1.79E-31 |
| 3 | 14165614 | 14179610 | 1.79E-31 |
| 3 | 14166979 | 14180329 | 1.79E-31 |
| 3 | 14167117 | 14180486 | 1.79E-31 |
| 3 | 14181871 | 14200532 | 1.79E-31 |
| 3 | 14182321 | 14201174 | 1.79E-31 |
| 3 | 14182863 | 14201804 | 1.79E-31 |
| 3 | 14182973 | 14202567 | 1.79E-31 |
| 3 | 14183034 | 14202666 | 1.79E-31 |
| 3 | 14183339 | 14202680 | 1.79E-31 |
| 3 | 14183737 | 14203074 | 1.79E-31 |

| | | | |
|---|---|---|---|
| 3 | 14186001 | 14203787 | 1.79E-31 |
| 3 | 14189033 | 14206296 | 1.79E-31 |
| 3 | 14192064 | 14207025 | 1.79E-31 |
| 3 | 14209829 | 14227273 | 1.79E-31 |
| 3 | 14242617 | 14254767 | 1.79E-31 |
| 3 | 14243120 | 14255977 | 1.79E-31 |
| 3 | 14243201 | 14256333 | 1.79E-31 |
| 3 | 14243445 | 14258228 | 1.79E-31 |
| 3 | 14243878 | 14258442 | 1.79E-31 |
| 3 | 14244281 | 14258495 | 1.79E-31 |
| 3 | 14246035 | 14261092 | 1.79E-31 |
| 3 | 14312015 | 14338005 | 1.79E-31 |
| 3 | 14313663 | 14339559 | 1.79E-31 |
| 3 | 14316846 | 14340946 | 1.79E-31 |
| 3 | 14316897 | 14341113 | 1.79E-31 |
| 3 | 14317050 | 14341489 | 1.79E-31 |
| 3 | 14324700 | 14345435 | 1.79E-31 |
| 3 | 14367262 | 14389849 | 1.79E-31 |
| 3 | 14367327 | 14390065 | 1.79E-31 |
| 3 | 14371873 | 14391011 | 1.79E-31 |
| 3 | 14375279 | 14391885 | 1.79E-31 |
| 3 | 14376155 | 14393255 | 1.79E-31 |
| 3 | 14376385 | 14393586 | 1.79E-31 |
| 3 | 14377920 | 14394057 | 1.79E-31 |
| 3 | 14382109 | 14395902 | 1.79E-31 |
| 3 | 14383982 | 14399752 | 1.79E-31 |
| 3 | 14384276 | 14399800 | 1.79E-31 |
| 3 | 14384282 | 14400440 | 1.79E-31 |
| 3 | 14384662 | 14401459 | 1.79E-31 |

| | | | |
|---|---|---|---|
| 3 | 14385064 | 14401590 | 1.79E-31 |
| 3 | 14385358 | 14401699 | 1.79E-31 |
| 3 | 14386222 | 14402966 | 1.79E-31 |
| 3 | 14391146 | 14407547 | 1.79E-31 |
| 3 | 14391885 | 14407833 | 1.79E-31 |
| 3 | 14393586 | 14408990 | 1.79E-31 |
| 3 | 14394287 | 14411952 | 1.79E-31 |
| 3 | 14404639 | 14419930 | 1.79E-31 |
| 3 | 14404733 | 14420321 | 1.79E-31 |
| 3 | 14406060 | 14420635 | 1.79E-31 |
| 3 | 14406152 | 14421085 | 1.79E-31 |
| 3 | 14428996 | 14442796 | 1.79E-31 |
| 3 | 14429222 | 14443186 | 1.79E-31 |
| 3 | 14429532 | 14443461 | 1.79E-31 |
| 3 | 14430142 | 14443664 | 1.79E-31 |
| 3 | 14447220 | 14463676 | 1.79E-31 |
| 3 | 14461440 | 14479758 | 1.79E-31 |
| 3 | 14462233 | 14480140 | 1.79E-31 |
| 3 | 14463676 | 14480186 | 1.79E-31 |
| 3 | 14528020 | 14545510 | 1.79E-31 |
| 3 | 14529083 | 14546639 | 1.79E-31 |
| 3 | 14529605 | 14548688 | 1.79E-31 |
| 3 | 14529813 | 14549368 | 1.79E-31 |
| 3 | 14539178 | 14558643 | 1.79E-31 |
| 3 | 14539651 | 14560092 | 1.79E-31 |
| 3 | 14539963 | 14561266 | 1.79E-31 |
| 3 | 14540770 | 14561601 | 1.79E-31 |
| 3 | 14542054 | 14561666 | 1.79E-31 |
| 3 | 14567048 | 14587447 | 1.79E-31 |

| | | | |
|---|---|---|---|
| 3 | 14567066 | 14587979 | 1.79E-31 |
| 3 | 14567205 | 14588635 | 1.79E-31 |
| 3 | 14567737 | 14588731 | 1.79E-31 |
| 3 | 14571597 | 14588932 | 1.79E-31 |
| 3 | 14576989 | 14591538 | 1.79E-31 |
| 3 | 14580058 | 14593498 | 1.79E-31 |
| 3 | 14581441 | 14594586 | 1.79E-31 |
| 3 | 14587288 | 14600987 | 1.79E-31 |
| 3 | 13927131 | 13940314 | 2.80E-29 |
| 3 | 13927457 | 13940694 | 2.80E-29 |
| 3 | 13930385 | 13945189 | 2.80E-29 |
| 3 | 13932962 | 13948142 | 2.80E-29 |
| 3 | 13933720 | 13948540 | 2.80E-29 |
| 3 | 13934297 | 13948874 | 2.80E-29 |
| 3 | 13935013 | 13949950 | 2.80E-29 |
| 3 | 13936876 | 13953091 | 2.80E-29 |
| 3 | 13937384 | 13953461 | 2.80E-29 |
| 3 | 14158598 | 14171311 | 2.80E-29 |
| 3 | 14164535 | 14178604 | 2.80E-29 |
| 3 | 14168070 | 14180784 | 2.80E-29 |
| 3 | 14181855 | 14200359 | 2.80E-29 |
| 3 | 14183965 | 14203198 | 2.80E-29 |
| 3 | 14184586 | 14203367 | 2.80E-29 |
| 3 | 14184800 | 14203732 | 2.80E-29 |
| 3 | 14186870 | 14204189 | 2.80E-29 |
| 3 | 14187047 | 14205247 | 2.80E-29 |
| 3 | 14188055 | 14206109 | 2.80E-29 |
| 3 | 14192588 | 14207438 | 2.80E-29 |
| 3 | 14209000 | 14225902 | 2.80E-29 |

| | | | |
|---|---|---|---|
| 3 | 14244829 | 14258607 | 2.80E-29 |
| 3 | 14245013 | 14258613 | 2.80E-29 |
| 3 | 14245957 | 14260538 | 2.80E-29 |
| 3 | 14313844 | 14340492 | 2.80E-29 |
| 3 | 14314339 | 14340705 | 2.80E-29 |
| 3 | 14324790 | 14345822 | 2.80E-29 |
| 3 | 14326120 | 14347159 | 2.80E-29 |
| 3 | 14327123 | 14348095 | 2.80E-29 |
| 3 | 14364331 | 14387516 | 2.80E-29 |
| 3 | 14364625 | 14388746 | 2.80E-29 |
| 3 | 14366067 | 14389469 | 2.80E-29 |
| 3 | 14366800 | 14389492 | 2.80E-29 |
| 3 | 14366884 | 14389713 | 2.80E-29 |
| 3 | 14387290 | 14403228 | 2.80E-29 |
| 3 | 14389713 | 14406060 | 2.80E-29 |
| 3 | 14389849 | 14406152 | 2.80E-29 |
| 3 | 14390065 | 14406217 | 2.80E-29 |
| 3 | 14390222 | 14407154 | 2.80E-29 |
| 3 | 14391011 | 14407332 | 2.80E-29 |
| 3 | 14395078 | 14412985 | 2.80E-29 |
| 3 | 14402966 | 14418959 | 2.80E-29 |
| 3 | 14403228 | 14419904 | 2.80E-29 |
| 3 | 14405202 | 14420339 | 2.80E-29 |
| 3 | 14405703 | 14420562 | 2.80E-29 |
| 3 | 14406217 | 14421171 | 2.80E-29 |
| 3 | 14427673 | 14442019 | 2.80E-29 |
| 3 | 14428984 | 14442536 | 2.80E-29 |
| 3 | 14448799 | 14463994 | 2.80E-29 |
| 3 | 14449082 | 14464815 | 2.80E-29 |

| | | | |
|---|---|---|---|
| 3 | 14449255 | 14465425 | 2.80E-29 |
| 3 | 14449429 | 14465691 | 2.80E-29 |
| 3 | 14459347 | 14478481 | 2.80E-29 |
| 3 | 14459412 | 14479071 | 2.80E-29 |
| 3 | 14459607 | 14479216 | 2.80E-29 |
| 3 | 14529659 | 14549088 | 2.80E-29 |
| 3 | 14530279 | 14549375 | 2.80E-29 |
| 3 | 14530650 | 14551758 | 2.80E-29 |
| 3 | 14539024 | 14557853 | 2.80E-29 |
| 3 | 14539247 | 14559493 | 2.80E-29 |
| 3 | 14542539 | 14561977 | 2.80E-29 |
| 3 | 14564849 | 14581728 | 2.80E-29 |
| 3 | 14566323 | 14587288 | 2.80E-29 |
| 3 | 14571712 | 14589498 | 2.80E-29 |
| 3 | 14573167 | 14590116 | 2.80E-29 |
| 3 | 14573597 | 14590567 | 2.80E-29 |
| 3 | 14574296 | 14591262 | 2.80E-29 |
| 3 | 14576814 | 14591389 | 2.80E-29 |
| 3 | 14577557 | 14592342 | 2.80E-29 |
| 3 | 14579153 | 14592741 | 2.80E-29 |
| 3 | 14579813 | 14592830 | 2.80E-29 |
| 3 | 14587447 | 14601122 | 2.80E-29 |
| 3 | 13927121 | 13939849 | 3.87E-27 |
| 3 | 13928316 | 13941092 | 3.87E-27 |
| 3 | 13928507 | 13941605 | 3.87E-27 |
| 3 | 13930108 | 13942680 | 3.87E-27 |
| 3 | 13937577 | 13954810 | 3.87E-27 |
| 3 | 14158019 | 14171218 | 3.87E-27 |
| 3 | 14158345 | 14171256 | 3.87E-27 |

| | | | |
|---|---|---|---|
| 3 | 14168191 | 14181077 | 3.87E-27 |
| 3 | 14181408 | 14200023 | 3.87E-27 |
| 3 | 14193151 | 14208177 | 3.87E-27 |
| 3 | 14208243 | 14225463 | 3.87E-27 |
| 3 | 14245333 | 14259277 | 3.87E-27 |
| 3 | 14245461 | 14259756 | 3.87E-27 |
| 3 | 14326297 | 14347576 | 3.87E-27 |
| 3 | 14326951 | 14347582 | 3.87E-27 |
| 3 | 14327551 | 14348300 | 3.87E-27 |
| 3 | 14340705 | 14353846 | 3.87E-27 |
| 3 | 14340946 | 14354359 | 3.87E-27 |
| 3 | 14341113 | 14354927 | 3.87E-27 |
| 3 | 14341489 | 14355252 | 3.87E-27 |
| 3 | 14342521 | 14356433 | 3.87E-27 |
| 3 | 14353469 | 14376385 | 3.87E-27 |
| 3 | 14353662 | 14377920 | 3.87E-27 |
| 3 | 14353846 | 14378027 | 3.87E-27 |
| 3 | 14354359 | 14378667 | 3.87E-27 |
| 3 | 14354927 | 14379483 | 3.87E-27 |
| 3 | 14359812 | 14383982 | 3.87E-27 |
| 3 | 14364195 | 14387290 | 3.87E-27 |
| 3 | 14387516 | 14404639 | 3.87E-27 |
| 3 | 14388746 | 14404733 | 3.87E-27 |
| 3 | 14389469 | 14405202 | 3.87E-27 |
| 3 | 14389492 | 14405703 | 3.87E-27 |
| 3 | 14395209 | 14413259 | 3.87E-27 |
| 3 | 14395902 | 14413606 | 3.87E-27 |
| 3 | 14395955 | 14413699 | 3.87E-27 |
| 3 | 14397397 | 14414060 | 3.87E-27 |

| | | | |
|---|---|---|---|
| 3 | 14399800 | 14417056 | 3.87E-27 |
| 3 | 14401459 | 14417384 | 3.87E-27 |
| 3 | 14401699 | 14418566 | 3.87E-27 |
| 3 | 14407154 | 14421338 | 3.87E-27 |
| 3 | 14407332 | 14422244 | 3.87E-27 |
| 3 | 14407547 | 14422808 | 3.87E-27 |
| 3 | 14426688 | 14440048 | 3.87E-27 |
| 3 | 14427056 | 14441915 | 3.87E-27 |
| 3 | 14449767 | 14466104 | 3.87E-27 |
| 3 | 14449801 | 14467574 | 3.87E-27 |
| 3 | 14458231 | 14478152 | 3.87E-27 |
| 3 | 14531146 | 14551837 | 3.87E-27 |
| 3 | 14531797 | 14552959 | 3.87E-27 |
| 3 | 14533650 | 14553242 | 3.87E-27 |
| 3 | 14535131 | 14556174 | 3.87E-27 |
| 3 | 14535207 | 14556296 | 3.87E-27 |
| 3 | 14535254 | 14556507 | 3.87E-27 |
| 3 | 14543183 | 14562282 | 3.87E-27 |
| 3 | 14549368 | 14565813 | 3.87E-27 |
| 3 | 14564114 | 14581245 | 3.87E-27 |
| 3 | 14564405 | 14581441 | 3.87E-27 |
| 3 | 14565186 | 14582506 | 3.87E-27 |
| 3 | 14566146 | 14585690 | 3.87E-27 |
| 3 | 14572005 | 14589848 | 3.87E-27 |
| 3 | 14573072 | 14590003 | 3.87E-27 |
| 3 | 14587979 | 14601183 | 3.87E-27 |
| 3 | 14616608 | 14636045 | 3.87E-27 |
| 3 | 14617653 | 14638595 | 3.87E-27 |
| 3 | 14625606 | 14642339 | 3.87E-27 |

| | | | |
|---|---|---|---|
| 3 | 14631811 | 14645036 | 3.87E-27 |
| 3 | 14632368 | 14645223 | 3.87E-27 |
| 3 | 14632644 | 14645718 | 3.87E-27 |
| 3 | 14653682 | 14664800 | 3.87E-27 |
| 3 | 14654903 | 14665375 | 3.87E-27 |
| 3 | 14656756 | 14669819 | 3.87E-27 |
| 3 | 14681550 | 14698619 | 3.87E-27 |
| 3 | 13926765 | 13939585 | 4.74E-25 |
| 3 | 13930087 | 13942028 | 4.74E-25 |
| 3 | 13937931 | 13957037 | 4.74E-25 |
| 3 | 14156941 | 14169724 | 4.74E-25 |
| 3 | 14156984 | 14169775 | 4.74E-25 |
| 3 | 14157992 | 14169959 | 4.74E-25 |
| 3 | 14168713 | 14181097 | 4.74E-25 |
| 3 | 14181097 | 14198965 | 4.74E-25 |
| 3 | 14193439 | 14208243 | 4.74E-25 |
| 3 | 14198965 | 14212244 | 4.74E-25 |
| 3 | 14200023 | 14212295 | 4.74E-25 |
| 3 | 14208177 | 14224669 | 4.74E-25 |
| 3 | 14328530 | 14349004 | 4.74E-25 |
| 3 | 14328787 | 14349777 | 4.74E-25 |
| 3 | 14328961 | 14349821 | 4.74E-25 |
| 3 | 14329305 | 14350241 | 4.74E-25 |
| 3 | 14329777 | 14350636 | 4.74E-25 |
| 3 | 14330096 | 14352025 | 4.74E-25 |
| 3 | 14336257 | 14352388 | 4.74E-25 |
| 3 | 14338005 | 14352837 | 4.74E-25 |
| 3 | 14338619 | 14352927 | 4.74E-25 |
| 3 | 14340492 | 14353662 | 4.74E-25 |

| | | | |
|---|---|---|---|
| 3 | 14342734 | 14357193 | 4.74E-25 |
| 3 | 14343342 | 14357207 | 4.74E-25 |
| 3 | 14352185 | 14370499 | 4.74E-25 |
| 3 | 14352379 | 14371873 | 4.74E-25 |
| 3 | 14352388 | 14373320 | 4.74E-25 |
| 3 | 14352837 | 14375279 | 4.74E-25 |
| 3 | 14352927 | 14376155 | 4.74E-25 |
| 3 | 14355252 | 14382109 | 4.74E-25 |
| 3 | 14356433 | 14382424 | 4.74E-25 |
| 3 | 14358350 | 14383295 | 4.74E-25 |
| 3 | 14360241 | 14384276 | 4.74E-25 |
| 3 | 14360292 | 14384282 | 4.74E-25 |
| 3 | 14360830 | 14384662 | 4.74E-25 |
| 3 | 14361072 | 14385064 | 4.74E-25 |
| 3 | 14362529 | 14385358 | 4.74E-25 |
| 3 | 14362794 | 14386222 | 4.74E-25 |
| 3 | 14397500 | 14414097 | 4.74E-25 |
| 3 | 14399032 | 14414276 | 4.74E-25 |
| 3 | 14399256 | 14414791 | 4.74E-25 |
| 3 | 14399752 | 14416746 | 4.74E-25 |
| 3 | 14400440 | 14417218 | 4.74E-25 |
| 3 | 14401590 | 14417871 | 4.74E-25 |
| 3 | 14407833 | 14423679 | 4.74E-25 |
| 3 | 14416746 | 14429222 | 4.74E-25 |
| 3 | 14417056 | 14429532 | 4.74E-25 |
| 3 | 14426640 | 14439915 | 4.74E-25 |
| 3 | 14449777 | 14466483 | 4.74E-25 |
| 3 | 14450226 | 14467741 | 4.74E-25 |
| 3 | 14450278 | 14468833 | 4.74E-25 |

| | | | |
|---|---|---|---|
| 3 | 14451307 | 14470922 | 4.74E-25 |
| 3 | 14458126 | 14477339 | 4.74E-25 |
| 3 | 14534150 | 14554785 | 4.74E-25 |
| 3 | 14534477 | 14556129 | 4.74E-25 |
| 3 | 14543456 | 14562486 | 4.74E-25 |
| 3 | 14543483 | 14562972 | 4.74E-25 |
| 3 | 14544257 | 14563075 | 4.74E-25 |
| 3 | 14549088 | 14565370 | 4.74E-25 |
| 3 | 14549375 | 14565970 | 4.74E-25 |
| 3 | 14551758 | 14566000 | 4.74E-25 |
| 3 | 14556174 | 14567066 | 4.74E-25 |
| 3 | 14556296 | 14567205 | 4.74E-25 |
| 3 | 14557853 | 14571597 | 4.74E-25 |
| 3 | 14558643 | 14571712 | 4.74E-25 |
| 3 | 14559493 | 14572005 | 4.74E-25 |
| 3 | 14560092 | 14573072 | 4.74E-25 |
| 3 | 14561266 | 14573167 | 4.74E-25 |
| 3 | 14561297 | 14573597 | 4.74E-25 |
| 3 | 14562486 | 14579153 | 4.74E-25 |
| 3 | 14562972 | 14579813 | 4.74E-25 |
| 3 | 14564024 | 14580137 | 4.74E-25 |
| 3 | 14565298 | 14582656 | 4.74E-25 |
| 3 | 14566138 | 14585379 | 4.74E-25 |
| 3 | 14588635 | 14601193 | 4.74E-25 |
| 3 | 14615739 | 14635850 | 4.74E-25 |
| 3 | 14616940 | 14637086 | 4.74E-25 |
| 3 | 14617588 | 14638561 | 4.74E-25 |
| 3 | 14618071 | 14638768 | 4.74E-25 |
| 3 | 14618672 | 14639336 | 4.74E-25 |

| | | | |
|---|---|---|---|
| 3 | 14619668 | 14639953 | 4.74E-25 |
| 3 | 14620773 | 14640185 | 4.74E-25 |
| 3 | 14622669 | 14640239 | 4.74E-25 |
| 3 | 14624370 | 14641355 | 4.74E-25 |
| 3 | 14625243 | 14641622 | 4.74E-25 |
| 3 | 14625531 | 14641731 | 4.74E-25 |
| 3 | 14626262 | 14642807 | 4.74E-25 |
| 3 | 14626283 | 14643176 | 4.74E-25 |
| 3 | 14628401 | 14644251 | 4.74E-25 |
| 3 | 14632716 | 14646206 | 4.74E-25 |
| 3 | 14632816 | 14646500 | 4.74E-25 |
| 3 | 14652846 | 14664181 | 4.74E-25 |
| 3 | 14653668 | 14664659 | 4.74E-25 |
| 3 | 14655029 | 14666063 | 4.74E-25 |
| 3 | 14655071 | 14666493 | 4.74E-25 |
| 3 | 14656345 | 14669651 | 4.74E-25 |
| 3 | 14656947 | 14670254 | 4.74E-25 |
| 3 | 14680176 | 14696948 | 4.74E-25 |
| 3 | 14680944 | 14698550 | 4.74E-25 |
| 3 | 14682613 | 14699063 | 4.74E-25 |
| 3 | 14684401 | 14700067 | 4.74E-25 |
| 3 | 13926534 | 13939481 | 5.15E-23 |
| 3 | 13938056 | 13957234 | 5.15E-23 |
| 3 | 13938061 | 13957569 | 5.15E-23 |
| 3 | 14156806 | 14168723 | 5.15E-23 |
| 3 | 14156913 | 14169059 | 5.15E-23 |
| 3 | 14168723 | 14181408 | 5.15E-23 |
| 3 | 14169724 | 14181871 | 5.15E-23 |
| 3 | 14180784 | 14198065 | 5.15E-23 |

| | | | |
|---|---|---|---|
| 3 | 14181077 | 14198173 | 5.15E-23 |
| 3 | 14193629 | 14209000 | 5.15E-23 |
| 3 | 14193728 | 14210441 | 5.15E-23 |
| 3 | 14195235 | 14210950 | 5.15E-23 |
| 3 | 14198065 | 14211062 | 5.15E-23 |
| 3 | 14198173 | 14211865 | 5.15E-23 |
| 3 | 14200359 | 14213396 | 5.15E-23 |
| 3 | 14207438 | 14223942 | 5.15E-23 |
| 3 | 14331204 | 14352047 | 5.15E-23 |
| 3 | 14333990 | 14352379 | 5.15E-23 |
| 3 | 14339559 | 14353469 | 5.15E-23 |
| 3 | 14343703 | 14357355 | 5.15E-23 |
| 3 | 14345435 | 14360241 | 5.15E-23 |
| 3 | 14345822 | 14360292 | 5.15E-23 |
| 3 | 14347159 | 14360830 | 5.15E-23 |
| 3 | 14347582 | 14362529 | 5.15E-23 |
| 3 | 14350241 | 14366800 | 5.15E-23 |
| 3 | 14352025 | 14367262 | 5.15E-23 |
| 3 | 14352047 | 14367327 | 5.15E-23 |
| 3 | 14357193 | 14382444 | 5.15E-23 |
| 3 | 14357207 | 14382816 | 5.15E-23 |
| 3 | 14357355 | 14383217 | 5.15E-23 |
| 3 | 14408386 | 14424303 | 5.15E-23 |
| 3 | 14408990 | 14424510 | 5.15E-23 |
| 3 | 14409629 | 14424883 | 5.15E-23 |
| 3 | 14414791 | 14428996 | 5.15E-23 |
| 3 | 14417218 | 14430142 | 5.15E-23 |
| 3 | 14417384 | 14430578 | 5.15E-23 |
| 3 | 14425074 | 14437731 | 5.15E-23 |

| | | | |
|---|---|---|---|
| 3 | 14425850 | 14439698 | 5.15E-23 |
| 3 | 14426581 | 14439903 | 5.15E-23 |
| 3 | 14450260 | 14467902 | 5.15E-23 |
| 3 | 14450722 | 14469044 | 5.15E-23 |
| 3 | 14450788 | 14469469 | 5.15E-23 |
| 3 | 14451167 | 14469645 | 5.15E-23 |
| 3 | 14451228 | 14469821 | 5.15E-23 |
| 3 | 14452731 | 14471313 | 5.15E-23 |
| 3 | 14457355 | 14473108 | 5.15E-23 |
| 3 | 14457922 | 14476829 | 5.15E-23 |
| 3 | 14544511 | 14564024 | 5.15E-23 |
| 3 | 14544522 | 14564114 | 5.15E-23 |
| 3 | 14546639 | 14565186 | 5.15E-23 |
| 3 | 14548688 | 14565298 | 5.15E-23 |
| 3 | 14551837 | 14566041 | 5.15E-23 |
| 3 | 14552959 | 14566138 | 5.15E-23 |
| 3 | 14553242 | 14566146 | 5.15E-23 |
| 3 | 14556129 | 14567048 | 5.15E-23 |
| 3 | 14556507 | 14567737 | 5.15E-23 |
| 3 | 14561601 | 14574296 | 5.15E-23 |
| 3 | 14561666 | 14576814 | 5.15E-23 |
| 3 | 14561977 | 14576989 | 5.15E-23 |
| 3 | 14562282 | 14577557 | 5.15E-23 |
| 3 | 14563075 | 14580058 | 5.15E-23 |
| 3 | 14565370 | 14583210 | 5.15E-23 |
| 3 | 14565813 | 14583525 | 5.15E-23 |
| 3 | 14565970 | 14583614 | 5.15E-23 |
| 3 | 14566000 | 14585061 | 5.15E-23 |
| 3 | 14566041 | 14585078 | 5.15E-23 |

| | | | |
|---|---|---|---|
| 3 | 14588731 | 14601708 | 5.15E-23 |
| 3 | 14589848 | 14602664 | 5.15E-23 |
| 3 | 14590003 | 14602775 | 5.15E-23 |
| 3 | 14592741 | 14604579 | 5.15E-23 |
| 3 | 14592830 | 14604631 | 5.15E-23 |
| 3 | 14595811 | 14607617 | 5.15E-23 |
| 3 | 14595904 | 14607645 | 5.15E-23 |
| 3 | 14596484 | 14607760 | 5.15E-23 |
| 3 | 14603091 | 14614515 | 5.15E-23 |
| 3 | 14615480 | 14634892 | 5.15E-23 |
| 3 | 14620984 | 14640224 | 5.15E-23 |
| 3 | 14623806 | 14640462 | 5.15E-23 |
| 3 | 14623938 | 14640767 | 5.15E-23 |
| 3 | 14633478 | 14646682 | 5.15E-23 |
| 3 | 14651036 | 14663627 | 5.15E-23 |
| 3 | 14653647 | 14664543 | 5.15E-23 |
| 3 | 14655480 | 14668925 | 5.15E-23 |
| 3 | 14655604 | 14669351 | 5.15E-23 |
| 3 | 14655702 | 14669527 | 5.15E-23 |
| 3 | 14657659 | 14672194 | 5.15E-23 |
| 3 | 14657903 | 14672416 | 5.15E-23 |
| 3 | 14658001 | 14672658 | 5.15E-23 |
| 3 | 14658273 | 14672864 | 5.15E-23 |
| 3 | 14658434 | 14672947 | 5.15E-23 |
| 3 | 14659786 | 14673967 | 5.15E-23 |
| 3 | 14660033 | 14674027 | 5.15E-23 |
| 3 | 14672947 | 14686890 | 5.15E-23 |
| 3 | 14673967 | 14687468 | 5.15E-23 |
| 3 | 14674027 | 14688169 | 5.15E-23 |

| | | | |
|---|---|---|---|
| 3 | 14674050 | 14688771 | 5.15E-23 |
| 3 | 14674971 | 14689352 | 5.15E-23 |
| 3 | 14675218 | 14689382 | 5.15E-23 |
| 3 | 14676141 | 14689973 | 5.15E-23 |
| 3 | 14676400 | 14690998 | 5.15E-23 |
| 3 | 14676974 | 14691785 | 5.15E-23 |
| 3 | 14677138 | 14692031 | 5.15E-23 |
| 3 | 14678247 | 14693546 | 5.15E-23 |
| 3 | 14678470 | 14693627 | 5.15E-23 |
| 3 | 14678536 | 14695357 | 5.15E-23 |
| 3 | 14678838 | 14695942 | 5.15E-23 |
| 3 | 14679878 | 14696052 | 5.15E-23 |
| 3 | 14682664 | 14699291 | 5.15E-23 |
| 3 | 14683356 | 14699477 | 5.15E-23 |
| 3 | 14683489 | 14699861 | 5.15E-23 |
| 3 | 14684426 | 14700668 | 5.15E-23 |
| 3 | 14690998 | 14708565 | 5.15E-23 |
| 3 | 13926472 | 13939359 | 4.95E-21 |
| 3 | 13939087 | 13957866 | 4.95E-21 |
| 3 | 13942680 | 13969159 | 4.95E-21 |
| 3 | 14156445 | 14168191 | 4.95E-21 |
| 3 | 14156453 | 14168713 | 4.95E-21 |
| 3 | 14169059 | 14181855 | 4.95E-21 |
| 3 | 14169775 | 14182321 | 4.95E-21 |
| 3 | 14178604 | 14193629 | 4.95E-21 |
| 3 | 14179219 | 14193722 | 4.95E-21 |
| 3 | 14179610 | 14193728 | 4.95E-21 |
| 3 | 14180329 | 14194313 | 4.95E-21 |
| 3 | 14180486 | 14195235 | 4.95E-21 |

| | | | |
|---|---|---|---|
| 3 | 14193722 | 14209829 | 4.95E-21 |
| 3 | 14194313 | 14210537 | 4.95E-21 |
| 3 | 14200532 | 14213888 | 4.95E-21 |
| 3 | 14202680 | 14217397 | 4.95E-21 |
| 3 | 14203074 | 14217848 | 4.95E-21 |
| 3 | 14203367 | 14218000 | 4.95E-21 |
| 3 | 14206296 | 14221486 | 4.95E-21 |
| 3 | 14207025 | 14223875 | 4.95E-21 |
| 3 | 14332360 | 14352185 | 4.95E-21 |
| 3 | 14344758 | 14358350 | 4.95E-21 |
| 3 | 14345107 | 14359812 | 4.95E-21 |
| 3 | 14347576 | 14361072 | 4.95E-21 |
| 3 | 14348095 | 14362794 | 4.95E-21 |
| 3 | 14349004 | 14364331 | 4.95E-21 |
| 3 | 14349777 | 14364625 | 4.95E-21 |
| 3 | 14349821 | 14366067 | 4.95E-21 |
| 3 | 14350636 | 14366884 | 4.95E-21 |
| 3 | 14411952 | 14425074 | 4.95E-21 |
| 3 | 14412985 | 14425850 | 4.95E-21 |
| 3 | 14413259 | 14426581 | 4.95E-21 |
| 3 | 14413606 | 14426640 | 4.95E-21 |
| 3 | 14413699 | 14426688 | 4.95E-21 |
| 3 | 14414060 | 14427056 | 4.95E-21 |
| 3 | 14414097 | 14427673 | 4.95E-21 |
| 3 | 14414276 | 14428984 | 4.95E-21 |
| 3 | 14417871 | 14430584 | 4.95E-21 |
| 3 | 14418566 | 14430594 | 4.95E-21 |
| 3 | 14424510 | 14435540 | 4.95E-21 |
| 3 | 14424883 | 14437423 | 4.95E-21 |

| | | | |
|---|---|---|---|
| 3 | 14454302 | 14471561 | 4.95E-21 |
| 3 | 14456821 | 14472401 | 4.95E-21 |
| 3 | 14457032 | 14472508 | 4.95E-21 |
| 3 | 14457166 | 14472627 | 4.95E-21 |
| 3 | 14457883 | 14476763 | 4.95E-21 |
| 3 | 14544674 | 14564405 | 4.95E-21 |
| 3 | 14545510 | 14564849 | 4.95E-21 |
| 3 | 14554785 | 14566323 | 4.95E-21 |
| 3 | 14588932 | 14601753 | 4.95E-21 |
| 3 | 14589498 | 14602551 | 4.95E-21 |
| 3 | 14590116 | 14603080 | 4.95E-21 |
| 3 | 14590567 | 14603091 | 4.95E-21 |
| 3 | 14591262 | 14603286 | 4.95E-21 |
| 3 | 14591538 | 14603927 | 4.95E-21 |
| 3 | 14592342 | 14604193 | 4.95E-21 |
| 3 | 14593498 | 14605065 | 4.95E-21 |
| 3 | 14593616 | 14606230 | 4.95E-21 |
| 3 | 14594409 | 14606333 | 4.95E-21 |
| 3 | 14594586 | 14606637 | 4.95E-21 |
| 3 | 14597251 | 14607963 | 4.95E-21 |
| 3 | 14603080 | 14613734 | 4.95E-21 |
| 3 | 14603286 | 14614626 | 4.95E-21 |
| 3 | 14615100 | 14634338 | 4.95E-21 |
| 3 | 14615202 | 14634521 | 4.95E-21 |
| 3 | 14633791 | 14647162 | 4.95E-21 |
| 3 | 14634319 | 14647881 | 4.95E-21 |
| 3 | 14641731 | 14656947 | 4.95E-21 |
| 3 | 14642339 | 14657659 | 4.95E-21 |
| 3 | 14646682 | 14660864 | 4.95E-21 |

| | | | |
|---|---|---|---|
| 3 | 14647162 | 14660928 | 4.95E-21 |
| 3 | 14647881 | 14661098 | 4.95E-21 |
| 3 | 14648465 | 14661468 | 4.95E-21 |
| 3 | 14648977 | 14661625 | 4.95E-21 |
| 3 | 14650260 | 14662314 | 4.95E-21 |
| 3 | 14650313 | 14663486 | 4.95E-21 |
| 3 | 14660047 | 14674050 | 4.95E-21 |
| 3 | 14662314 | 14678470 | 4.95E-21 |
| 3 | 14663486 | 14678536 | 4.95E-21 |
| 3 | 14663627 | 14678838 | 4.95E-21 |
| 3 | 14672864 | 14686823 | 4.95E-21 |
| 3 | 14684666 | 14701140 | 4.95E-21 |
| 3 | 14689973 | 14706723 | 4.95E-21 |
| 3 | 14691785 | 14708890 | 4.95E-21 |
| 3 | 14692031 | 14708904 | 4.95E-21 |
| 3 | 14693546 | 14709424 | 4.95E-21 |
| 3 | 14695357 | 14709565 | 4.95E-21 |
| 3 | 13926336 | 13939241 | 4.21E-19 |
| 3 | 13939241 | 13960629 | 4.21E-19 |
| 3 | 13939359 | 13962248 | 4.21E-19 |
| 3 | 13942028 | 13968881 | 4.21E-19 |
| 3 | 13945189 | 13969248 | 4.21E-19 |
| 3 | 14150556 | 14163195 | 4.21E-19 |
| 3 | 14152097 | 14163506 | 4.21E-19 |
| 3 | 14154255 | 14164466 | 4.21E-19 |
| 3 | 14154397 | 14164500 | 4.21E-19 |
| 3 | 14155713 | 14167117 | 4.21E-19 |
| 3 | 14155826 | 14168070 | 4.21E-19 |
| 3 | 14169959 | 14182863 | 4.21E-19 |

| | | | |
|---|---|---|---|
| 3 | 14171218 | 14182973 | 4.21E-19 |
| 3 | 14171311 | 14183339 | 4.21E-19 |
| 3 | 14173092 | 14183737 | 4.21E-19 |
| 3 | 14173314 | 14183965 | 4.21E-19 |
| 3 | 14173473 | 14184586 | 4.21E-19 |
| 3 | 14173949 | 14186870 | 4.21E-19 |
| 3 | 14177632 | 14192588 | 4.21E-19 |
| 3 | 14178033 | 14193151 | 4.21E-19 |
| 3 | 14178099 | 14193439 | 4.21E-19 |
| 3 | 14201174 | 14214259 | 4.21E-19 |
| 3 | 14202567 | 14215640 | 4.21E-19 |
| 3 | 14202666 | 14215870 | 4.21E-19 |
| 3 | 14203198 | 14217876 | 4.21E-19 |
| 3 | 14203732 | 14219560 | 4.21E-19 |
| 3 | 14203787 | 14219752 | 4.21E-19 |
| 3 | 14204189 | 14220807 | 4.21E-19 |
| 3 | 14205247 | 14220987 | 4.21E-19 |
| 3 | 14206109 | 14221266 | 4.21E-19 |
| 3 | 14206306 | 14222030 | 4.21E-19 |
| 3 | 14206773 | 14223813 | 4.21E-19 |
| 3 | 14348300 | 14364195 | 4.21E-19 |
| 3 | 14418959 | 14431441 | 4.21E-19 |
| 3 | 14424303 | 14435308 | 4.21E-19 |
| 3 | 14591389 | 14603445 | 4.21E-19 |
| 3 | 14598749 | 14608103 | 4.21E-19 |
| 3 | 14599339 | 14608477 | 4.21E-19 |
| 3 | 14599434 | 14608859 | 4.21E-19 |
| 3 | 14602664 | 14613033 | 4.21E-19 |
| 3 | 14602775 | 14613295 | 4.21E-19 |

| | | | |
|---|---|---|---|
| 3 | 14603445 | 14614929 | 4.21E-19 |
| 3 | 14603927 | 14615100 | 4.21E-19 |
| 3 | 14613033 | 14632644 | 4.21E-19 |
| 3 | 14614929 | 14634319 | 4.21E-19 |
| 3 | 14634338 | 14648465 | 4.21E-19 |
| 3 | 14634521 | 14648977 | 4.21E-19 |
| 3 | 14634892 | 14649376 | 4.21E-19 |
| 3 | 14640239 | 14655480 | 4.21E-19 |
| 3 | 14640462 | 14655604 | 4.21E-19 |
| 3 | 14640767 | 14655702 | 4.21E-19 |
| 3 | 14641355 | 14656345 | 4.21E-19 |
| 3 | 14641622 | 14656756 | 4.21E-19 |
| 3 | 14642807 | 14657903 | 4.21E-19 |
| 3 | 14643176 | 14658001 | 4.21E-19 |
| 3 | 14644251 | 14658273 | 4.21E-19 |
| 3 | 14645036 | 14658434 | 4.21E-19 |
| 3 | 14646206 | 14660047 | 4.21E-19 |
| 3 | 14646500 | 14660778 | 4.21E-19 |
| 3 | 14649376 | 14661872 | 4.21E-19 |
| 3 | 14660778 | 14674971 | 4.21E-19 |
| 3 | 14660864 | 14675218 | 4.21E-19 |
| 3 | 14660928 | 14676141 | 4.21E-19 |
| 3 | 14661098 | 14676400 | 4.21E-19 |
| 3 | 14661468 | 14676974 | 4.21E-19 |
| 3 | 14661625 | 14677138 | 4.21E-19 |
| 3 | 14661872 | 14678247 | 4.21E-19 |
| 3 | 14664181 | 14679878 | 4.21E-19 |
| 3 | 14672416 | 14686368 | 4.21E-19 |
| 3 | 14672658 | 14686625 | 4.21E-19 |

| | | | |
|---|---|---|---|
| 3 | 14685836 | 14701373 | 4.21E-19 |
| 3 | 14686213 | 14702625 | 4.21E-19 |
| 3 | 14686320 | 14702685 | 4.21E-19 |
| 3 | 14689352 | 14706305 | 4.21E-19 |
| 3 | 14689382 | 14706596 | 4.21E-19 |
| 3 | 14693627 | 14709458 | 4.21E-19 |
| 3 | 14695942 | 14710290 | 4.21E-19 |
| 3 | 14696052 | 14710474 | 4.21E-19 |
| 3 | 14696948 | 14711951 | 4.21E-19 |
| 3 | 13924656 | 13937384 | 3.16E-17 |
| 3 | 13925064 | 13937577 | 3.16E-17 |
| 3 | 13925104 | 13937931 | 3.16E-17 |
| 3 | 13925956 | 13938061 | 3.16E-17 |
| 3 | 13926068 | 13939087 | 3.16E-17 |
| 3 | 13939481 | 13962683 | 3.16E-17 |
| 3 | 13941605 | 13968510 | 3.16E-17 |
| 3 | 13945483 | 13969494 | 3.16E-17 |
| 3 | 13991351 | 14017423 | 3.16E-17 |
| 3 | 13991762 | 14017464 | 3.16E-17 |
| 3 | 14149858 | 14162967 | 3.16E-17 |
| 3 | 14153120 | 14163733 | 3.16E-17 |
| 3 | 14155000 | 14164535 | 3.16E-17 |
| 3 | 14155064 | 14164662 | 3.16E-17 |
| 3 | 14155099 | 14165614 | 3.16E-17 |
| 3 | 14155705 | 14166979 | 3.16E-17 |
| 3 | 14171256 | 14183034 | 3.16E-17 |
| 3 | 14173604 | 14184800 | 3.16E-17 |
| 3 | 14173760 | 14186001 | 3.16E-17 |
| 3 | 14174210 | 14187047 | 3.16E-17 |

| | | | |
|---|---|---|---|
| 3 | 14175413 | 14188055 | 3.16E-17 |
| 3 | 14177485 | 14192064 | 3.16E-17 |
| 3 | 14201804 | 14214841 | 3.16E-17 |
| 3 | 14419904 | 14432337 | 3.16E-17 |
| 3 | 14419930 | 14432352 | 3.16E-17 |
| 3 | 14420562 | 14433210 | 3.16E-17 |
| 3 | 14420635 | 14434236 | 3.16E-17 |
| 3 | 14421085 | 14434698 | 3.16E-17 |
| 3 | 14422244 | 14435057 | 3.16E-17 |
| 3 | 14422808 | 14435093 | 3.16E-17 |
| 3 | 14423679 | 14435206 | 3.16E-17 |
| 3 | 14600215 | 14609417 | 3.16E-17 |
| 3 | 14600481 | 14609695 | 3.16E-17 |
| 3 | 14600722 | 14610158 | 3.16E-17 |
| 3 | 14600987 | 14610911 | 3.16E-17 |
| 3 | 14601122 | 14611049 | 3.16E-17 |
| 3 | 14601183 | 14611059 | 3.16E-17 |
| 3 | 14601193 | 14611798 | 3.16E-17 |
| 3 | 14601708 | 14612130 | 3.16E-17 |
| 3 | 14601753 | 14612850 | 3.16E-17 |
| 3 | 14602551 | 14612987 | 3.16E-17 |
| 3 | 14604193 | 14615202 | 3.16E-17 |
| 3 | 14606230 | 14616940 | 3.16E-17 |
| 3 | 14612850 | 14631811 | 3.16E-17 |
| 3 | 14612987 | 14632368 | 3.16E-17 |
| 3 | 14613295 | 14632716 | 3.16E-17 |
| 3 | 14613734 | 14632816 | 3.16E-17 |
| 3 | 14614515 | 14633478 | 3.16E-17 |
| 3 | 14614626 | 14633791 | 3.16E-17 |

| | | | |
|---|---|---|---|
| 3 | 14635850 | 14650260 | 3.16E-17 |
| 3 | 14640224 | 14655071 | 3.16E-17 |
| 3 | 14645223 | 14659786 | 3.16E-17 |
| 3 | 14645718 | 14660033 | 3.16E-17 |
| 3 | 14664543 | 14680176 | 3.16E-17 |
| 3 | 14664659 | 14680944 | 3.16E-17 |
| 3 | 14664800 | 14681550 | 3.16E-17 |
| 3 | 14665375 | 14682613 | 3.16E-17 |
| 3 | 14666063 | 14682664 | 3.16E-17 |
| 3 | 14666493 | 14683356 | 3.16E-17 |
| 3 | 14669351 | 14684401 | 3.16E-17 |
| 3 | 14669527 | 14684426 | 3.16E-17 |
| 3 | 14669651 | 14684666 | 3.16E-17 |
| 3 | 14669819 | 14685836 | 3.16E-17 |
| 3 | 14672194 | 14686320 | 3.16E-17 |
| 3 | 14686368 | 14703083 | 3.16E-17 |
| 3 | 14686823 | 14703746 | 3.16E-17 |
| 3 | 14688771 | 14706289 | 3.16E-17 |
| 3 | 14698550 | 14711995 | 3.16E-17 |
| 3 | 14698619 | 14712441 | 3.16E-17 |
| 3 | 14699291 | 14712703 | 3.16E-17 |
| 3 | 14699477 | 14713100 | 3.16E-17 |
| 3 | 13734743 | 13751134 | 2.09E-15 |
| 3 | 13734815 | 13751278 | 2.09E-15 |
| 3 | 13735113 | 13751300 | 2.09E-15 |
| 3 | 13737141 | 13751758 | 2.09E-15 |
| 3 | 13737685 | 13751778 | 2.09E-15 |
| 3 | 13822726 | 13836335 | 2.09E-15 |
| 3 | 13822776 | 13836503 | 2.09E-15 |

| | | | |
|---|---|---|---|
| 3 | 13823137 | 13837234 | 2.09E-15 |
| 3 | 13823947 | 13837573 | 2.09E-15 |
| 3 | 13824064 | 13838115 | 2.09E-15 |
| 3 | 13824265 | 13838186 | 2.09E-15 |
| 3 | 13824275 | 13838413 | 2.09E-15 |
| 3 | 13824664 | 13838667 | 2.09E-15 |
| 3 | 13825727 | 13839045 | 2.09E-15 |
| 3 | 13876499 | 13896769 | 2.09E-15 |
| 3 | 13887812 | 13906038 | 2.09E-15 |
| 3 | 13887890 | 13906977 | 2.09E-15 |
| 3 | 13916797 | 13927457 | 2.09E-15 |
| 3 | 13917650 | 13928507 | 2.09E-15 |
| 3 | 13917681 | 13930087 | 2.09E-15 |
| 3 | 13917810 | 13930108 | 2.09E-15 |
| 3 | 13923256 | 13933720 | 2.09E-15 |
| 3 | 13923347 | 13934297 | 2.09E-15 |
| 3 | 13924216 | 13935013 | 2.09E-15 |
| 3 | 13924589 | 13936876 | 2.09E-15 |
| 3 | 13925283 | 13938056 | 2.09E-15 |
| 3 | 13939585 | 13963451 | 2.09E-15 |
| 3 | 13939849 | 13966580 | 2.09E-15 |
| 3 | 13940314 | 13967728 | 2.09E-15 |
| 3 | 13940694 | 13968226 | 2.09E-15 |
| 3 | 13941092 | 13968432 | 2.09E-15 |
| 3 | 13945508 | 13969965 | 2.09E-15 |
| 3 | 13990736 | 14015186 | 2.09E-15 |
| 3 | 13992372 | 14017968 | 2.09E-15 |
| 3 | 13992379 | 14018759 | 2.09E-15 |
| 3 | 13992480 | 14019252 | 2.09E-15 |

| | | | |
|---|---|---|---|
| 3 | 14090912 | 14123078 | 2.09E-15 |
| 3 | 14093969 | 14124753 | 2.09E-15 |
| 3 | 14094056 | 14124835 | 2.09E-15 |
| 3 | 14149062 | 14161810 | 2.09E-15 |
| 3 | 14175877 | 14189033 | 2.09E-15 |
| 3 | 14177023 | 14191946 | 2.09E-15 |
| 3 | 14420321 | 14432630 | 2.09E-15 |
| 3 | 14420339 | 14433071 | 2.09E-15 |
| 3 | 14421171 | 14434986 | 2.09E-15 |
| 3 | 14421338 | 14435021 | 2.09E-15 |
| 3 | 14604579 | 14615480 | 2.09E-15 |
| 3 | 14605065 | 14616608 | 2.09E-15 |
| 3 | 14606333 | 14617588 | 2.09E-15 |
| 3 | 14606637 | 14617653 | 2.09E-15 |
| 3 | 14607617 | 14618071 | 2.09E-15 |
| 3 | 14611049 | 14625606 | 2.09E-15 |
| 3 | 14611059 | 14626262 | 2.09E-15 |
| 3 | 14611798 | 14626283 | 2.09E-15 |
| 3 | 14612130 | 14628401 | 2.09E-15 |
| 3 | 14636045 | 14650313 | 2.09E-15 |
| 3 | 14638561 | 14652846 | 2.09E-15 |
| 3 | 14640185 | 14655029 | 2.09E-15 |
| 3 | 14668925 | 14683489 | 2.09E-15 |
| 3 | 14670254 | 14686213 | 2.09E-15 |
| 3 | 14686625 | 14703185 | 2.09E-15 |
| 3 | 14686890 | 14703781 | 2.09E-15 |
| 3 | 14688169 | 14705175 | 2.09E-15 |
| 3 | 14699063 | 14712459 | 2.09E-15 |
| 3 | 14699861 | 14713489 | 2.09E-15 |

| | | | |
|---|---|---|---|
| 3 | 14700067 | 14713933 | 2.09E-15 |
| 3 | 14703746 | 14717440 | 2.09E-15 |
| 3 | 13734731 | 13750978 | 1.20E-13 |
| 3 | 13737991 | 13752673 | 1.20E-13 |
| 3 | 13741563 | 13755127 | 1.20E-13 |
| 3 | 13741963 | 13757358 | 1.20E-13 |
| 3 | 13742019 | 13757721 | 1.20E-13 |
| 3 | 13822284 | 13836121 | 1.20E-13 |
| 3 | 13826239 | 13839282 | 1.20E-13 |
| 3 | 13826424 | 13839828 | 1.20E-13 |
| 3 | 13869881 | 13888020 | 1.20E-13 |
| 3 | 13871706 | 13891143 | 1.20E-13 |
| 3 | 13872080 | 13892806 | 1.20E-13 |
| 3 | 13872601 | 13894872 | 1.20E-13 |
| 3 | 13872725 | 13895019 | 1.20E-13 |
| 3 | 13873675 | 13896086 | 1.20E-13 |
| 3 | 13874069 | 13896566 | 1.20E-13 |
| 3 | 13878753 | 13898665 | 1.20E-13 |
| 3 | 13879151 | 13898880 | 1.20E-13 |
| 3 | 13886510 | 13905365 | 1.20E-13 |
| 3 | 13887362 | 13905890 | 1.20E-13 |
| 3 | 13888020 | 13906997 | 1.20E-13 |
| 3 | 13914791 | 13925956 | 1.20E-13 |
| 3 | 13914983 | 13926068 | 1.20E-13 |
| 3 | 13915478 | 13926336 | 1.20E-13 |
| 3 | 13915502 | 13926472 | 1.20E-13 |
| 3 | 13915938 | 13926534 | 1.20E-13 |
| 3 | 13916094 | 13926765 | 1.20E-13 |
| 3 | 13916592 | 13927121 | 1.20E-13 |

| | | | |
|---|---|---|---|
| 3 | 13916622 | 13927131 | 1.20E-13 |
| 3 | 13917605 | 13928316 | 1.20E-13 |
| 3 | 13918163 | 13930385 | 1.20E-13 |
| 3 | 13918472 | 13930894 | 1.20E-13 |
| 3 | 13919122 | 13931245 | 1.20E-13 |
| 3 | 13919320 | 13931553 | 1.20E-13 |
| 3 | 13920312 | 13931562 | 1.20E-13 |
| 3 | 13922133 | 13932646 | 1.20E-13 |
| 3 | 13923075 | 13932962 | 1.20E-13 |
| 3 | 13945630 | 13971596 | 1.20E-13 |
| 3 | 13978079 | 14007800 | 1.20E-13 |
| 3 | 13981062 | 14010019 | 1.20E-13 |
| 3 | 13982133 | 14011580 | 1.20E-13 |
| 3 | 13983040 | 14012450 | 1.20E-13 |
| 3 | 13984873 | 14012623 | 1.20E-13 |
| 3 | 13990380 | 14014124 | 1.20E-13 |
| 3 | 13993825 | 14019547 | 1.20E-13 |
| 3 | 13994391 | 14020138 | 1.20E-13 |
| 3 | 14083809 | 14101765 | 1.20E-13 |
| 3 | 14085457 | 14102704 | 1.20E-13 |
| 3 | 14086332 | 14103612 | 1.20E-13 |
| 3 | 14090260 | 14120642 | 1.20E-13 |
| 3 | 14090851 | 14121852 | 1.20E-13 |
| 3 | 14091408 | 14123668 | 1.20E-13 |
| 3 | 14092088 | 14124039 | 1.20E-13 |
| 3 | 14093171 | 14124289 | 1.20E-13 |
| 3 | 14093868 | 14124575 | 1.20E-13 |
| 3 | 14094170 | 14125001 | 1.20E-13 |
| 3 | 14095644 | 14125500 | 1.20E-13 |

| | | | |
|---|---|---|---|
| 3 | 14098668 | 14126067 | 1.20E-13 |
| 3 | 14129988 | 14154397 | 1.20E-13 |
| 3 | 14130311 | 14155000 | 1.20E-13 |
| 3 | 14130335 | 14155064 | 1.20E-13 |
| 3 | 14130971 | 14155099 | 1.20E-13 |
| 3 | 14132297 | 14156445 | 1.20E-13 |
| 3 | 14135722 | 14156941 | 1.20E-13 |
| 3 | 14136082 | 14156984 | 1.20E-13 |
| 3 | 14136742 | 14157992 | 1.20E-13 |
| 3 | 14140067 | 14159619 | 1.20E-13 |
| 3 | 14145834 | 14159816 | 1.20E-13 |
| 3 | 14146449 | 14160692 | 1.20E-13 |
| 3 | 14147387 | 14160814 | 1.20E-13 |
| 3 | 14147768 | 14160884 | 1.20E-13 |
| 3 | 14176443 | 14191658 | 1.20E-13 |
| 3 | 14604631 | 14615739 | 1.20E-13 |
| 3 | 14607645 | 14618672 | 1.20E-13 |
| 3 | 14608477 | 14622669 | 1.20E-13 |
| 3 | 14608859 | 14623806 | 1.20E-13 |
| 3 | 14609417 | 14623938 | 1.20E-13 |
| 3 | 14609695 | 14624370 | 1.20E-13 |
| 3 | 14610158 | 14625243 | 1.20E-13 |
| 3 | 14610911 | 14625531 | 1.20E-13 |
| 3 | 14637086 | 14651036 | 1.20E-13 |
| 3 | 14638595 | 14653647 | 1.20E-13 |
| 3 | 14639336 | 14653682 | 1.20E-13 |
| 3 | 14639953 | 14654903 | 1.20E-13 |
| 3 | 14687468 | 14704442 | 1.20E-13 |
| 3 | 14700668 | 14714356 | 1.20E-13 |

| | | | |
|---|---|---|---|
| 3 | 14701140 | 14715380 | 1.20E-13 |
| 3 | 14701373 | 14715816 | 1.20E-13 |
| 3 | 14702625 | 14716128 | 1.20E-13 |
| 3 | 14702685 | 14716340 | 1.20E-13 |
| 3 | 14703083 | 14716472 | 1.20E-13 |
| 3 | 14703185 | 14716772 | 1.20E-13 |
| 3 | 14703781 | 14717904 | 1.20E-13 |
| 3 | 14704442 | 14718102 | 1.20E-13 |
| 3 | 14705175 | 14718396 | 1.20E-13 |
| 3 | 14706289 | 14719326 | 1.20E-13 |
| 3 | 14706305 | 14719510 | 1.20E-13 |
| 3 | 14706596 | 14719826 | 1.20E-13 |
| 3 | 14706723 | 14720443 | 1.20E-13 |
| 3 | 14712703 | 14725843 | 1.20E-13 |
| 3 | 14713489 | 14726883 | 1.20E-13 |
| 3 | 14713933 | 14728463 | 1.20E-13 |
| 3 | 14714356 | 14728605 | 1.20E-13 |
| 3 | 14715380 | 14728909 | 1.20E-13 |
| 3 | 14715816 | 14728953 | 1.20E-13 |
| 3 | 14716128 | 14729015 | 1.20E-13 |
| 3 | 14716340 | 14729063 | 1.20E-13 |
| 3 | 14716472 | 14729384 | 1.20E-13 |
| 3 | 14716772 | 14729562 | 1.20E-13 |
| 3 | 14717440 | 14730573 | 1.20E-13 |
| 3 | 14924472 | 14941393 | 1.20E-13 |
| 3 | 14925557 | 14942876 | 1.20E-13 |
| 3 | 14925564 | 14942906 | 1.20E-13 |
| 3 | 14925576 | 14943006 | 1.20E-13 |
| 3 | 14925600 | 14943051 | 1.20E-13 |

| | | | |
|---|---|---|---|
| 3 | 14926042 | 14946078 | 1.20E-13 |
| 3 | 14956317 | 14973319 | 1.20E-13 |
| 3 | 14958686 | 14973409 | 1.20E-13 |
| 3 | 13406039 | 13421871 | 6.05E-12 |
| 3 | 13406995 | 13421905 | 6.05E-12 |
| 3 | 13407476 | 13422044 | 6.05E-12 |
| 3 | 13407998 | 13422370 | 6.05E-12 |
| 3 | 13408005 | 13423218 | 6.05E-12 |
| 3 | 13409248 | 13423298 | 6.05E-12 |
| 3 | 13410173 | 13423325 | 6.05E-12 |
| 3 | 13410417 | 13423674 | 6.05E-12 |
| 3 | 13410617 | 13423721 | 6.05E-12 |
| 3 | 13411069 | 13424598 | 6.05E-12 |
| 3 | 13411458 | 13425089 | 6.05E-12 |
| 3 | 13411588 | 13425834 | 6.05E-12 |
| 3 | 13413726 | 13429988 | 6.05E-12 |
| 3 | 13413982 | 13430881 | 6.05E-12 |
| 3 | 13414362 | 13431557 | 6.05E-12 |
| 3 | 13415213 | 13432104 | 6.05E-12 |
| 3 | 13699937 | 13719494 | 6.05E-12 |
| 3 | 13703879 | 13722645 | 6.05E-12 |
| 3 | 13708681 | 13728620 | 6.05E-12 |
| 3 | 13708847 | 13729026 | 6.05E-12 |
| 3 | 13708857 | 13730989 | 6.05E-12 |
| 3 | 13708948 | 13731024 | 6.05E-12 |
| 3 | 13709248 | 13731257 | 6.05E-12 |
| 3 | 13715417 | 13734190 | 6.05E-12 |
| 3 | 13732723 | 13748712 | 6.05E-12 |
| 3 | 13733664 | 13748797 | 6.05E-12 |

| | | | |
|---|---|---|---|
| 3 | 13734363 | 13750495 | 6.05E-12 |
| 3 | 13734658 | 13750638 | 6.05E-12 |
| 3 | 13734711 | 13750799 | 6.05E-12 |
| 3 | 13738144 | 13753900 | 6.05E-12 |
| 3 | 13739243 | 13753979 | 6.05E-12 |
| 3 | 13739269 | 13754099 | 6.05E-12 |
| 3 | 13740234 | 13754606 | 6.05E-12 |
| 3 | 13741159 | 13754706 | 6.05E-12 |
| 3 | 13741549 | 13754865 | 6.05E-12 |
| 3 | 13742085 | 13757735 | 6.05E-12 |
| 3 | 13744960 | 13757765 | 6.05E-12 |
| 3 | 13746227 | 13758126 | 6.05E-12 |
| 3 | 13746426 | 13758230 | 6.05E-12 |
| 3 | 13746783 | 13758535 | 6.05E-12 |
| 3 | 13746934 | 13758541 | 6.05E-12 |
| 3 | 13748529 | 13759080 | 6.05E-12 |
| 3 | 13771236 | 13797641 | 6.05E-12 |
| 3 | 13819593 | 13830811 | 6.05E-12 |
| 3 | 13820033 | 13831850 | 6.05E-12 |
| 3 | 13820051 | 13833033 | 6.05E-12 |
| 3 | 13820164 | 13833383 | 6.05E-12 |
| 3 | 13820616 | 13834235 | 6.05E-12 |
| 3 | 13821206 | 13834682 | 6.05E-12 |
| 3 | 13821228 | 13834815 | 6.05E-12 |
| 3 | 13821363 | 13835626 | 6.05E-12 |
| 3 | 13821369 | 13835788 | 6.05E-12 |
| 3 | 13822094 | 13835794 | 6.05E-12 |
| 3 | 13826481 | 13840222 | 6.05E-12 |
| 3 | 13827157 | 13842619 | 6.05E-12 |

| | | | |
|---|---|---|---|
| 3 | 13842619 | 13855561 | 6.05E-12 |
| 3 | 13843224 | 13857423 | 6.05E-12 |
| 3 | 13843858 | 13858080 | 6.05E-12 |
| 3 | 13843942 | 13858090 | 6.05E-12 |
| 3 | 13844170 | 13858514 | 6.05E-12 |
| 3 | 13869367 | 13887890 | 6.05E-12 |
| 3 | 13871179 | 13889640 | 6.05E-12 |
| 3 | 13871692 | 13890744 | 6.05E-12 |
| 3 | 13873054 | 13895234 | 6.05E-12 |
| 3 | 13880335 | 13899029 | 6.05E-12 |
| 3 | 13881351 | 13900362 | 6.05E-12 |
| 3 | 13881461 | 13900689 | 6.05E-12 |
| 3 | 13881715 | 13901436 | 6.05E-12 |
| 3 | 13883023 | 13902739 | 6.05E-12 |
| 3 | 13883261 | 13902988 | 6.05E-12 |
| 3 | 13884036 | 13903371 | 6.05E-12 |
| 3 | 13884248 | 13903779 | 6.05E-12 |
| 3 | 13884969 | 13904643 | 6.05E-12 |
| 3 | 13885800 | 13904803 | 6.05E-12 |
| 3 | 13889640 | 13907328 | 6.05E-12 |
| 3 | 13890744 | 13907336 | 6.05E-12 |
| 3 | 13891143 | 13908874 | 6.05E-12 |
| 3 | 13911533 | 13925104 | 6.05E-12 |
| 3 | 13914239 | 13925283 | 6.05E-12 |
| 3 | 13918267 | 13930492 | 6.05E-12 |
| 3 | 13921248 | 13931996 | 6.05E-12 |
| 3 | 13921357 | 13932641 | 6.05E-12 |
| 3 | 13945912 | 13971611 | 6.05E-12 |
| 3 | 13945944 | 13971768 | 6.05E-12 |

| | | | |
|---|---|---|---|
| 3 | 13946058 | 13972407 | 6.05E-12 |
| 3 | 13946259 | 13973425 | 6.05E-12 |
| 3 | 13946539 | 13973463 | 6.05E-12 |
| 3 | 13948142 | 13973502 | 6.05E-12 |
| 3 | 13948540 | 13973844 | 6.05E-12 |
| 3 | 13948874 | 13974350 | 6.05E-12 |
| 3 | 13977684 | 14007371 | 6.05E-12 |
| 3 | 13994727 | 14020522 | 6.05E-12 |
| 3 | 13996191 | 14021888 | 6.05E-12 |
| 3 | 13996302 | 14022974 | 6.05E-12 |
| 3 | 13996903 | 14023823 | 6.05E-12 |
| 3 | 13997916 | 14027994 | 6.05E-12 |
| 3 | 13998701 | 14030276 | 6.05E-12 |
| 3 | 14001830 | 14031207 | 6.05E-12 |
| 3 | 14002188 | 14034312 | 6.05E-12 |
| 3 | 14006488 | 14045401 | 6.05E-12 |
| 3 | 14006822 | 14045620 | 6.05E-12 |
| 3 | 14077596 | 14098668 | 6.05E-12 |
| 3 | 14077965 | 14100070 | 6.05E-12 |
| 3 | 14080396 | 14100156 | 6.05E-12 |
| 3 | 14081023 | 14100658 | 6.05E-12 |
| 3 | 14082316 | 14101643 | 6.05E-12 |
| 3 | 14086914 | 14104566 | 6.05E-12 |
| 3 | 14088117 | 14107315 | 6.05E-12 |
| 3 | 14088201 | 14107658 | 6.05E-12 |
| 3 | 14088484 | 14110744 | 6.05E-12 |
| 3 | 14089770 | 14110962 | 6.05E-12 |
| 3 | 14090206 | 14112158 | 6.05E-12 |
| 3 | 14090227 | 14114167 | 6.05E-12 |

| | | | |
|---|---|---|---|
| 3 | 14090816 | 14120951 | 6.05E-12 |
| 3 | 14100070 | 14126810 | 6.05E-12 |
| 3 | 14100156 | 14128983 | 6.05E-12 |
| 3 | 14100658 | 14129988 | 6.05E-12 |
| 3 | 14101643 | 14130311 | 6.05E-12 |
| 3 | 14103612 | 14131209 | 6.05E-12 |
| 3 | 14105371 | 14132297 | 6.05E-12 |
| 3 | 14106759 | 14134040 | 6.05E-12 |
| 3 | 14107315 | 14135133 | 6.05E-12 |
| 3 | 14128983 | 14154255 | 6.05E-12 |
| 3 | 14131209 | 14155705 | 6.05E-12 |
| 3 | 14131287 | 14155713 | 6.05E-12 |
| 3 | 14132130 | 14155826 | 6.05E-12 |
| 3 | 14134040 | 14156453 | 6.05E-12 |
| 3 | 14135133 | 14156806 | 6.05E-12 |
| 3 | 14135277 | 14156913 | 6.05E-12 |
| 3 | 14136902 | 14158019 | 6.05E-12 |
| 3 | 14138029 | 14158345 | 6.05E-12 |
| 3 | 14138064 | 14158598 | 6.05E-12 |
| 3 | 14138356 | 14159351 | 6.05E-12 |
| 3 | 14138946 | 14159552 | 6.05E-12 |
| 3 | 14144943 | 14159629 | 6.05E-12 |
| 3 | 14607760 | 14619668 | 6.05E-12 |
| 3 | 14607963 | 14620773 | 6.05E-12 |
| 3 | 14608103 | 14620984 | 6.05E-12 |
| 3 | 14638768 | 14653668 | 6.05E-12 |
| 3 | 14708565 | 14720491 | 6.05E-12 |
| 3 | 14709424 | 14721934 | 6.05E-12 |
| 3 | 14709458 | 14722347 | 6.05E-12 |

| | | | |
|---|---|---|---|
| 3 | 14709565 | 14722352 | 6.05E-12 |
| 3 | 14710474 | 14723583 | 6.05E-12 |
| 3 | 14712459 | 14725443 | 6.05E-12 |
| 3 | 14713100 | 14726855 | 6.05E-12 |
| 3 | 14717904 | 14730761 | 6.05E-12 |
| 3 | 14718102 | 14731365 | 6.05E-12 |
| 3 | 14718396 | 14731556 | 6.05E-12 |
| 3 | 14719326 | 14731768 | 6.05E-12 |
| 3 | 14719510 | 14732557 | 6.05E-12 |
| 3 | 14721934 | 14736161 | 6.05E-12 |
| 3 | 14723583 | 14738618 | 6.05E-12 |
| 3 | 14923879 | 14941359 | 6.05E-12 |
| 3 | 14926260 | 14946363 | 6.05E-12 |
| 3 | 14927246 | 14946810 | 6.05E-12 |
| 3 | 14927572 | 14946919 | 6.05E-12 |
| 3 | 14928569 | 14947005 | 6.05E-12 |
| 3 | 14928995 | 14947018 | 6.05E-12 |
| 3 | 14929346 | 14947505 | 6.05E-12 |
| 3 | 14929667 | 14948757 | 6.05E-12 |
| 3 | 14929734 | 14948993 | 6.05E-12 |
| 3 | 14950195 | 14967443 | 6.05E-12 |
| 3 | 14951124 | 14967717 | 6.05E-12 |
| 3 | 14951995 | 14968571 | 6.05E-12 |
| 3 | 14953958 | 14970888 | 6.05E-12 |
| 3 | 14954623 | 14971184 | 6.05E-12 |
| 3 | 14954868 | 14972525 | 6.05E-12 |
| 3 | 14955080 | 14972814 | 6.05E-12 |
| 3 | 14955188 | 14973195 | 6.05E-12 |
| 3 | 14959437 | 14973431 | 6.05E-12 |

| | | | |
|---|---|---|---|
| 3 | 14959473 | 14974644 | 6.05E-12 |
| 3 | 13396937 | 13411588 | 2.62E-10 |
| 3 | 13397808 | 13412362 | 2.62E-10 |
| 3 | 13398242 | 13412712 | 2.62E-10 |
| 3 | 13398895 | 13412808 | 2.62E-10 |
| 3 | 13405583 | 13420565 | 2.62E-10 |
| 3 | 13405852 | 13421204 | 2.62E-10 |
| 3 | 13412316 | 13426213 | 2.62E-10 |
| 3 | 13412362 | 13426931 | 2.62E-10 |
| 3 | 13412808 | 13428442 | 2.62E-10 |
| 3 | 13413163 | 13429328 | 2.62E-10 |
| 3 | 13413446 | 13429354 | 2.62E-10 |
| 3 | 13413524 | 13429367 | 2.62E-10 |
| 3 | 13415266 | 13432551 | 2.62E-10 |
| 3 | 13566069 | 13589160 | 2.62E-10 |
| 3 | 13684091 | 13703879 | 2.62E-10 |
| 3 | 13695109 | 13718167 | 2.62E-10 |
| 3 | 13696409 | 13718726 | 2.62E-10 |
| 3 | 13696655 | 13718949 | 2.62E-10 |
| 3 | 13697780 | 13718997 | 2.62E-10 |
| 3 | 13698854 | 13719239 | 2.62E-10 |
| 3 | 13699978 | 13719510 | 2.62E-10 |
| 3 | 13701443 | 13720106 | 2.62E-10 |
| 3 | 13702080 | 13721229 | 2.62E-10 |
| 3 | 13702298 | 13721461 | 2.62E-10 |
| 3 | 13703114 | 13722220 | 2.62E-10 |
| 3 | 13703497 | 13722593 | 2.62E-10 |
| 3 | 13705246 | 13724315 | 2.62E-10 |
| 3 | 13706536 | 13725397 | 2.62E-10 |

| | | | |
|---|---|---|---|
| 3 | 13707611 | 13726947 | 2.62E-10 |
| 3 | 13709301 | 13731442 | 2.62E-10 |
| 3 | 13710013 | 13731523 | 2.62E-10 |
| 3 | 13710687 | 13731582 | 2.62E-10 |
| 3 | 13711986 | 13731684 | 2.62E-10 |
| 3 | 13715039 | 13733664 | 2.62E-10 |
| 3 | 13718167 | 13734286 | 2.62E-10 |
| 3 | 13722593 | 13738144 | 2.62E-10 |
| 3 | 13732676 | 13748529 | 2.62E-10 |
| 3 | 13734190 | 13748872 | 2.62E-10 |
| 3 | 13734286 | 13750339 | 2.62E-10 |
| 3 | 13745718 | 13758044 | 2.62E-10 |
| 3 | 13748712 | 13759137 | 2.62E-10 |
| 3 | 13767703 | 13795059 | 2.62E-10 |
| 3 | 13769077 | 13795257 | 2.62E-10 |
| 3 | 13770068 | 13796050 | 2.62E-10 |
| 3 | 13771035 | 13797427 | 2.62E-10 |
| 3 | 13771524 | 13798175 | 2.62E-10 |
| 3 | 13771604 | 13798302 | 2.62E-10 |
| 3 | 13783908 | 13804504 | 2.62E-10 |
| 3 | 13784236 | 13804734 | 2.62E-10 |
| 3 | 13784823 | 13805211 | 2.62E-10 |
| 3 | 13786281 | 13806768 | 2.62E-10 |
| 3 | 13787536 | 13809309 | 2.62E-10 |
| 3 | 13787547 | 13809323 | 2.62E-10 |
| 3 | 13788988 | 13811457 | 2.62E-10 |
| 3 | 13790461 | 13811552 | 2.62E-10 |
| 3 | 13790515 | 13812208 | 2.62E-10 |
| 3 | 13791258 | 13812668 | 2.62E-10 |

| | | | |
|---|---|---|---|
| 3 | 13791505 | 13813343 | 2.62E-10 |
| 3 | 13791730 | 13814807 | 2.62E-10 |
| 3 | 13791995 | 13814890 | 2.62E-10 |
| 3 | 13792890 | 13815096 | 2.62E-10 |
| 3 | 13794700 | 13815365 | 2.62E-10 |
| 3 | 13795007 | 13815946 | 2.62E-10 |
| 3 | 13818407 | 13830029 | 2.62E-10 |
| 3 | 13818439 | 13830097 | 2.62E-10 |
| 3 | 13819363 | 13830655 | 2.62E-10 |
| 3 | 13827951 | 13843224 | 2.62E-10 |
| 3 | 13835794 | 13850257 | 2.62E-10 |
| 3 | 13836121 | 13850286 | 2.62E-10 |
| 3 | 13838667 | 13853786 | 2.62E-10 |
| 3 | 13839045 | 13853950 | 2.62E-10 |
| 3 | 13839282 | 13854561 | 2.62E-10 |
| 3 | 13839828 | 13854607 | 2.62E-10 |
| 3 | 13840222 | 13854833 | 2.62E-10 |
| 3 | 13844274 | 13858787 | 2.62E-10 |
| 3 | 13844628 | 13860732 | 2.62E-10 |
| 3 | 13844755 | 13860921 | 2.62E-10 |
| 3 | 13869363 | 13887812 | 2.62E-10 |
| 3 | 13881997 | 13902372 | 2.62E-10 |
| 3 | 13882005 | 13902468 | 2.62E-10 |
| 3 | 13882187 | 13902669 | 2.62E-10 |
| 3 | 13882781 | 13902734 | 2.62E-10 |
| 3 | 13892806 | 13908951 | 2.62E-10 |
| 3 | 13894872 | 13909089 | 2.62E-10 |
| 3 | 13895019 | 13909529 | 2.62E-10 |
| 3 | 13895234 | 13911533 | 2.62E-10 |

| | | | |
|---|---|---|---|
| 3 | 13896086 | 13914239 | 2.62E-10 |
| 3 | 13905365 | 13920312 | 2.62E-10 |
| 3 | 13909529 | 13925064 | 2.62E-10 |
| 3 | 13949950 | 13976449 | 2.62E-10 |
| 3 | 13953091 | 13976726 | 2.62E-10 |
| 3 | 13953461 | 13977684 | 2.62E-10 |
| 3 | 13976449 | 14006822 | 2.62E-10 |
| 3 | 13976726 | 14007250 | 2.62E-10 |
| 3 | 14002750 | 14036703 | 2.62E-10 |
| 3 | 14004463 | 14041253 | 2.62E-10 |
| 3 | 14005678 | 14041604 | 2.62E-10 |
| 3 | 14005941 | 14042368 | 2.62E-10 |
| 3 | 14006465 | 14043343 | 2.62E-10 |
| 3 | 14007250 | 14045694 | 2.62E-10 |
| 3 | 14007371 | 14046970 | 2.62E-10 |
| 3 | 14075444 | 14090912 | 2.62E-10 |
| 3 | 14075688 | 14091408 | 2.62E-10 |
| 3 | 14076494 | 14092088 | 2.62E-10 |
| 3 | 14076799 | 14094056 | 2.62E-10 |
| 3 | 14076855 | 14094170 | 2.62E-10 |
| 3 | 14077341 | 14095644 | 2.62E-10 |
| 3 | 14086962 | 14104725 | 2.62E-10 |
| 3 | 14087176 | 14105371 | 2.62E-10 |
| 3 | 14087567 | 14106759 | 2.62E-10 |
| 3 | 14088424 | 14108773 | 2.62E-10 |
| 3 | 14101765 | 14130335 | 2.62E-10 |
| 3 | 14102704 | 14130971 | 2.62E-10 |
| 3 | 14104566 | 14131287 | 2.62E-10 |
| 3 | 14104725 | 14132130 | 2.62E-10 |

| | | | |
|---|---|---|---|
| 3 | 14107658 | 14135277 | 2.62E-10 |
| 3 | 14108773 | 14135722 | 2.62E-10 |
| 3 | 14110744 | 14136082 | 2.62E-10 |
| 3 | 14110962 | 14136742 | 2.62E-10 |
| 3 | 14112158 | 14136902 | 2.62E-10 |
| 3 | 14114167 | 14138029 | 2.62E-10 |
| 3 | 14120642 | 14138064 | 2.62E-10 |
| 3 | 14126067 | 14152097 | 2.62E-10 |
| 3 | 14126810 | 14153120 | 2.62E-10 |
| 3 | 14708890 | 14720789 | 2.62E-10 |
| 3 | 14708904 | 14720859 | 2.62E-10 |
| 3 | 14710290 | 14723270 | 2.62E-10 |
| 3 | 14711951 | 14723809 | 2.62E-10 |
| 3 | 14711995 | 14723824 | 2.62E-10 |
| 3 | 14712441 | 14724313 | 2.62E-10 |
| 3 | 14719826 | 14732866 | 2.62E-10 |
| 3 | 14720789 | 14734078 | 2.62E-10 |
| 3 | 14720859 | 14735570 | 2.62E-10 |
| 3 | 14722347 | 14736925 | 2.62E-10 |
| 3 | 14722352 | 14737074 | 2.62E-10 |
| 3 | 14723270 | 14738605 | 2.62E-10 |
| 3 | 14723809 | 14738849 | 2.62E-10 |
| 3 | 14723824 | 14743086 | 2.62E-10 |
| 3 | 14724313 | 14743110 | 2.62E-10 |
| 3 | 14725443 | 14743149 | 2.62E-10 |
| 3 | 14729384 | 14748076 | 2.62E-10 |
| 3 | 14732557 | 14751134 | 2.62E-10 |
| 3 | 14916377 | 14932588 | 2.62E-10 |
| 3 | 14917001 | 14933482 | 2.62E-10 |

| | | | |
|---|---|---|---|
| 3 | 14919181 | 14934730 | 2.62E-10 |
| 3 | 14919620 | 14936259 | 2.62E-10 |
| 3 | 14919731 | 14937187 | 2.62E-10 |
| 3 | 14920090 | 14937293 | 2.62E-10 |
| 3 | 14920589 | 14937317 | 2.62E-10 |
| 3 | 14923694 | 14940551 | 2.62E-10 |
| 3 | 14930267 | 14949201 | 2.62E-10 |
| 3 | 14931599 | 14949550 | 2.62E-10 |
| 3 | 14946810 | 14963286 | 2.62E-10 |
| 3 | 14946919 | 14964225 | 2.62E-10 |
| 3 | 14947005 | 14964432 | 2.62E-10 |
| 3 | 14950002 | 14967198 | 2.62E-10 |
| 3 | 14952139 | 14968674 | 2.62E-10 |
| 3 | 14952568 | 14969337 | 2.62E-10 |
| 3 | 14953290 | 14970601 | 2.62E-10 |
| 3 | 14959850 | 14975473 | 2.62E-10 |
| 3 | 14964871 | 14981092 | 2.62E-10 |
| 3 | 14965905 | 14981390 | 2.62E-10 |
| 3 | 14965931 | 14981434 | 2.62E-10 |
| 3 | 14966180 | 14981449 | 2.62E-10 |
| 3 | 14966805 | 14982973 | 2.62E-10 |
| 3 | 14966880 | 14982997 | 2.62E-10 |
| 3 | 15124225 | 15145557 | 2.62E-10 |
| 3 | 15124952 | 15146980 | 2.62E-10 |
| 3 | 12984444 | 13001747 | 9.71E-09 |
| 3 | 13315135 | 13328415 | 9.71E-09 |
| 3 | 13315211 | 13329154 | 9.71E-09 |
| 3 | 13315283 | 13329829 | 9.71E-09 |
| 3 | 13315476 | 13331085 | 9.71E-09 |

| | | | |
|---|---|---|---|
| 3 | 13315794 | 13331806 | 9.71E-09 |
| 3 | 13316793 | 13331895 | 9.71E-09 |
| 3 | 13318813 | 13335749 | 9.71E-09 |
| 3 | 13319280 | 13336296 | 9.71E-09 |
| 3 | 13395310 | 13410173 | 9.71E-09 |
| 3 | 13396378 | 13410417 | 9.71E-09 |
| 3 | 13396744 | 13410617 | 9.71E-09 |
| 3 | 13396846 | 13411069 | 9.71E-09 |
| 3 | 13396894 | 13411458 | 9.71E-09 |
| 3 | 13397710 | 13412316 | 9.71E-09 |
| 3 | 13401803 | 13413163 | 9.71E-09 |
| 3 | 13405245 | 13419581 | 9.71E-09 |
| 3 | 13412712 | 13427797 | 9.71E-09 |
| 3 | 13416481 | 13433408 | 9.71E-09 |
| 3 | 13417076 | 13433635 | 9.71E-09 |
| 3 | 13417233 | 13433726 | 9.71E-09 |
| 3 | 13418111 | 13433884 | 9.71E-09 |
| 3 | 13519991 | 13539973 | 9.71E-09 |
| 3 | 13520014 | 13540140 | 9.71E-09 |
| 3 | 13564794 | 13587088 | 9.71E-09 |
| 3 | 13565177 | 13587898 | 9.71E-09 |
| 3 | 13565206 | 13588339 | 9.71E-09 |
| 3 | 13565600 | 13589102 | 9.71E-09 |
| 3 | 13567414 | 13589299 | 9.71E-09 |
| 3 | 13567428 | 13589488 | 9.71E-09 |
| 3 | 13567506 | 13590007 | 9.71E-09 |
| 3 | 13568345 | 13590044 | 9.71E-09 |
| 3 | 13568901 | 13590322 | 9.71E-09 |
| 3 | 13569077 | 13591917 | 9.71E-09 |

| | | | |
|---|---|---|---|
| 3 | 13572079 | 13592503 | 9.71E-09 |
| 3 | 13572559 | 13593047 | 9.71E-09 |
| 3 | 13678161 | 13699937 | 9.71E-09 |
| 3 | 13678446 | 13699978 | 9.71E-09 |
| 3 | 13678476 | 13701443 | 9.71E-09 |
| 3 | 13678974 | 13702080 | 9.71E-09 |
| 3 | 13680079 | 13702298 | 9.71E-09 |
| 3 | 13680926 | 13703114 | 9.71E-09 |
| 3 | 13682118 | 13703497 | 9.71E-09 |
| 3 | 13684445 | 13705246 | 9.71E-09 |
| 3 | 13685615 | 13706536 | 9.71E-09 |
| 3 | 13686209 | 13707611 | 9.71E-09 |
| 3 | 13687738 | 13709248 | 9.71E-09 |
| 3 | 13688805 | 13709301 | 9.71E-09 |
| 3 | 13689116 | 13710013 | 9.71E-09 |
| 3 | 13690214 | 13711986 | 9.71E-09 |
| 3 | 13690487 | 13712181 | 9.71E-09 |
| 3 | 13691932 | 13712197 | 9.71E-09 |
| 3 | 13693125 | 13713679 | 9.71E-09 |
| 3 | 13693331 | 13714157 | 9.71E-09 |
| 3 | 13693891 | 13715417 | 9.71E-09 |
| 3 | 13712181 | 13732009 | 9.71E-09 |
| 3 | 13712197 | 13732637 | 9.71E-09 |
| 3 | 13713679 | 13732676 | 9.71E-09 |
| 3 | 13714157 | 13732723 | 9.71E-09 |
| 3 | 13718726 | 13734363 | 9.71E-09 |
| 3 | 13718949 | 13734658 | 9.71E-09 |
| 3 | 13722220 | 13737991 | 9.71E-09 |
| 3 | 13722645 | 13739243 | 9.71E-09 |

| | | | |
|---|---|---|---|
| 3 | 13731442 | 13744960 | 9.71E-09 |
| 3 | 13731523 | 13745718 | 9.71E-09 |
| 3 | 13731582 | 13746227 | 9.71E-09 |
| 3 | 13731684 | 13746426 | 9.71E-09 |
| 3 | 13732009 | 13746783 | 9.71E-09 |
| 3 | 13732637 | 13746934 | 9.71E-09 |
| 3 | 13748797 | 13759557 | 9.71E-09 |
| 3 | 13748872 | 13760419 | 9.71E-09 |
| 3 | 13750339 | 13761588 | 9.71E-09 |
| 3 | 13767576 | 13795007 | 9.71E-09 |
| 3 | 13773192 | 13799265 | 9.71E-09 |
| 3 | 13775412 | 13799531 | 9.71E-09 |
| 3 | 13779477 | 13802506 | 9.71E-09 |
| 3 | 13780673 | 13803141 | 9.71E-09 |
| 3 | 13781645 | 13803176 | 9.71E-09 |
| 3 | 13781932 | 13803625 | 9.71E-09 |
| 3 | 13782379 | 13804277 | 9.71E-09 |
| 3 | 13785628 | 13805991 | 9.71E-09 |
| 3 | 13795059 | 13816526 | 9.71E-09 |
| 3 | 13817544 | 13829177 | 9.71E-09 |
| 3 | 13817872 | 13829366 | 9.71E-09 |
| 3 | 13819308 | 13830636 | 9.71E-09 |
| 3 | 13829177 | 13843858 | 9.71E-09 |
| 3 | 13830029 | 13844170 | 9.71E-09 |
| 3 | 13830097 | 13844274 | 9.71E-09 |
| 3 | 13830636 | 13844398 | 9.71E-09 |
| 3 | 13830655 | 13844628 | 9.71E-09 |
| 3 | 13830811 | 13844755 | 9.71E-09 |
| 3 | 13834235 | 13845418 | 9.71E-09 |

| | | | |
|---|---|---|---|
| 3 | 13834682 | 13847683 | 9.71E-09 |
| 3 | 13834815 | 13847727 | 9.71E-09 |
| 3 | 13835626 | 13849186 | 9.71E-09 |
| 3 | 13835788 | 13849405 | 9.71E-09 |
| 3 | 13836335 | 13850361 | 9.71E-09 |
| 3 | 13837573 | 13852279 | 9.71E-09 |
| 3 | 13838115 | 13852340 | 9.71E-09 |
| 3 | 13838186 | 13852781 | 9.71E-09 |
| 3 | 13838413 | 13852966 | 9.71E-09 |
| 3 | 13844398 | 13860073 | 9.71E-09 |
| 3 | 13844905 | 13861246 | 9.71E-09 |
| 3 | 13845200 | 13863195 | 9.71E-09 |
| 3 | 13845332 | 13863368 | 9.71E-09 |
| 3 | 13845418 | 13864019 | 9.71E-09 |
| 3 | 13863195 | 13882005 | 9.71E-09 |
| 3 | 13863368 | 13882187 | 9.71E-09 |
| 3 | 13864019 | 13882781 | 9.71E-09 |
| 3 | 13864500 | 13883023 | 9.71E-09 |
| 3 | 13865804 | 13883261 | 9.71E-09 |
| 3 | 13865823 | 13884036 | 9.71E-09 |
| 3 | 13866382 | 13884248 | 9.71E-09 |
| 3 | 13867170 | 13884969 | 9.71E-09 |
| 3 | 13867376 | 13885800 | 9.71E-09 |
| 3 | 13868543 | 13886510 | 9.71E-09 |
| 3 | 13869147 | 13887362 | 9.71E-09 |
| 3 | 13896566 | 13914791 | 9.71E-09 |
| 3 | 13896769 | 13914983 | 9.71E-09 |
| 3 | 13903779 | 13918472 | 9.71E-09 |
| 3 | 13904643 | 13919122 | 9.71E-09 |

| | | | |
|---|---|---|---|
| 3 | 13904803 | 13919320 | 9.71E-09 |
| 3 | 13905890 | 13921248 | 9.71E-09 |
| 3 | 13906038 | 13921357 | 9.71E-09 |
| 3 | 13909089 | 13924656 | 9.71E-09 |
| 3 | 13954810 | 13978079 | 9.71E-09 |
| 3 | 13957037 | 13981062 | 9.71E-09 |
| 3 | 13957234 | 13982133 | 9.71E-09 |
| 3 | 13957569 | 13983040 | 9.71E-09 |
| 3 | 13957866 | 13984873 | 9.71E-09 |
| 3 | 13960629 | 13990380 | 9.71E-09 |
| 3 | 13962248 | 13990736 | 9.71E-09 |
| 3 | 13963451 | 13991762 | 9.71E-09 |
| 3 | 13966580 | 13992372 | 9.71E-09 |
| 3 | 13973502 | 14005941 | 9.71E-09 |
| 3 | 13973844 | 14006465 | 9.71E-09 |
| 3 | 13974350 | 14006488 | 9.71E-09 |
| 3 | 14007800 | 14047098 | 9.71E-09 |
| 3 | 14065635 | 14088201 | 9.71E-09 |
| 3 | 14065644 | 14088424 | 9.71E-09 |
| 3 | 14066352 | 14088484 | 9.71E-09 |
| 3 | 14072522 | 14089770 | 9.71E-09 |
| 3 | 14073361 | 14090260 | 9.71E-09 |
| 3 | 14075240 | 14090816 | 9.71E-09 |
| 3 | 14075383 | 14090851 | 9.71E-09 |
| 3 | 14076663 | 14093171 | 9.71E-09 |
| 3 | 14076768 | 14093868 | 9.71E-09 |
| 3 | 14076792 | 14093969 | 9.71E-09 |
| 3 | 14120951 | 14138356 | 9.71E-09 |
| 3 | 14121852 | 14138946 | 9.71E-09 |

| | | | |
|---|---|---|---|
| 3 | 14123078 | 14140067 | 9.71E-09 |
| 3 | 14124289 | 14146449 | 9.71E-09 |
| 3 | 14124575 | 14147387 | 9.71E-09 |
| 3 | 14124753 | 14147768 | 9.71E-09 |
| 3 | 14125500 | 14150556 | 9.71E-09 |
| 3 | 14720443 | 14732879 | 9.71E-09 |
| 3 | 14720491 | 14733789 | 9.71E-09 |
| 3 | 14725843 | 14745895 | 9.71E-09 |
| 3 | 14729015 | 14747547 | 9.71E-09 |
| 3 | 14729063 | 14747894 | 9.71E-09 |
| 3 | 14729562 | 14748092 | 9.71E-09 |
| 3 | 14730573 | 14748133 | 9.71E-09 |
| 3 | 14730761 | 14748229 | 9.71E-09 |
| 3 | 14731365 | 14748342 | 9.71E-09 |
| 3 | 14731556 | 14750112 | 9.71E-09 |
| 3 | 14731768 | 14750718 | 9.71E-09 |
| 3 | 14732866 | 14751961 | 9.71E-09 |
| 3 | 14732879 | 14752286 | 9.71E-09 |
| 3 | 14733789 | 14755085 | 9.71E-09 |
| 3 | 14734078 | 14755547 | 9.71E-09 |
| 3 | 14751134 | 14770012 | 9.71E-09 |
| 3 | 14875044 | 14893207 | 9.71E-09 |
| 3 | 14875164 | 14893735 | 9.71E-09 |
| 3 | 14875221 | 14895845 | 9.71E-09 |
| 3 | 14876313 | 14896142 | 9.71E-09 |
| 3 | 14876421 | 14896312 | 9.71E-09 |
| 3 | 14877547 | 14896317 | 9.71E-09 |
| 3 | 14879115 | 14896503 | 9.71E-09 |
| 3 | 14879681 | 14897539 | 9.71E-09 |

| | | | |
|---|---|---|---|
| 3 | 14883446 | 14904869 | 9.71E-09 |
| 3 | 14883810 | 14905319 | 9.71E-09 |
| 3 | 14883879 | 14905835 | 9.71E-09 |
| 3 | 14916289 | 14931599 | 9.71E-09 |
| 3 | 14917877 | 14934016 | 9.71E-09 |
| 3 | 14919121 | 14934498 | 9.71E-09 |
| 3 | 14920854 | 14939350 | 9.71E-09 |
| 3 | 14920994 | 14939501 | 9.71E-09 |
| 3 | 14921273 | 14939937 | 9.71E-09 |
| 3 | 14923670 | 14940352 | 9.71E-09 |
| 3 | 14932588 | 14949797 | 9.71E-09 |
| 3 | 14934730 | 14951995 | 9.71E-09 |
| 3 | 14942906 | 14959850 | 9.71E-09 |
| 3 | 14943006 | 14961549 | 9.71E-09 |
| 3 | 14943051 | 14961821 | 9.71E-09 |
| 3 | 14946078 | 14962213 | 9.71E-09 |
| 3 | 14946363 | 14962560 | 9.71E-09 |
| 3 | 14947018 | 14964555 | 9.71E-09 |
| 3 | 14947505 | 14964871 | 9.71E-09 |
| 3 | 14948757 | 14965905 | 9.71E-09 |
| 3 | 14949797 | 14966880 | 9.71E-09 |
| 3 | 14961549 | 14976819 | 9.71E-09 |
| 3 | 14961821 | 14978067 | 9.71E-09 |
| 3 | 14962213 | 14978671 | 9.71E-09 |
| 3 | 14962560 | 14978813 | 9.71E-09 |
| 3 | 14963286 | 14979220 | 9.71E-09 |
| 3 | 14964225 | 14979323 | 9.71E-09 |
| 3 | 14964432 | 14980876 | 9.71E-09 |
| 3 | 14964555 | 14981019 | 9.71E-09 |

| | | | |
|---|---|---|---|
| 3 | 14967198 | 14983303 | 9.71E-09 |
| 3 | 14970601 | 14987725 | 9.71E-09 |
| 3 | 14970888 | 14988187 | 9.71E-09 |
| 3 | 14972525 | 14988445 | 9.71E-09 |
| 3 | 14972814 | 14990910 | 9.71E-09 |
| 3 | 14973195 | 14993187 | 9.71E-09 |
| 3 | 14973319 | 14994055 | 9.71E-09 |
| 3 | 15014464 | 15035173 | 9.71E-09 |
| 3 | 15015104 | 15037320 | 9.71E-09 |
| 3 | 15015473 | 15039052 | 9.71E-09 |
| 3 | 15015543 | 15039076 | 9.71E-09 |
| 3 | 15018420 | 15040879 | 9.71E-09 |
| 3 | 15019158 | 15040992 | 9.71E-09 |
| 3 | 15122514 | 15144520 | 9.71E-09 |
| 3 | 15123219 | 15145035 | 9.71E-09 |
| 3 | 15123385 | 15145325 | 9.71E-09 |
| 3 | 15123970 | 15145368 | 9.71E-09 |
| 3 | 15124999 | 15148090 | 9.71E-09 |
| 3 | 15126020 | 15148940 | 9.71E-09 |
| 3 | 15126168 | 15149022 | 9.71E-09 |
| 3 | 15252850 | 15276478 | 9.71E-09 |
| 3 | 15253651 | 15276710 | 9.71E-09 |
| 3 | 15254793 | 15276795 | 9.71E-09 |

**Supplementary Table 7A: Expression levels of candidate genes at the 3p25.1 locus in different cardiac tissues**

| | GTex Portal[1] | | Heinig et al. [2] | | | |
|---|---|---|---|---|---|---|
| | Heart Left Ventricle | Heart Atrial Appendage | Heart donors | DCM patients | Fold-change | P[3] |
| Gene | | | | | | |
| *CHCHD4* | 14.24 | 13.90 | 1.14 | 1.12 | -0.02 | 0.03 |
| *TMEM43* | 23.33 | 58.76 | 1.39 | 1.41 | 0.02 | 0.22 |
| *XPC* | 6.88 | 12.70 | 0.75 | 0.83 | 0.08 | $8.3\ 10^{-15}$ |
| *LSM3* | 12.19 | 16.41 | 1.07 | 1.03 | -0.04 | $7.6\ 10^{-8}$ |
| *SLC6A6* | 7.45 | 21.27 | 0.94 | 1.13 | 0.19 | $6.9\ 10^{-6}$ |
| *GRIP2* | 9.99 | 4.95 | 0.86 | 0.94 | 0.08 | 0.03 |

[1] Expression values are shown in TPM (transcript per million) calculated from a gene model with isoforms collapsed to a single gene. No other normalization steps have been performed

[2] In Heinig et al, left ventricle gene expression levels were expressed as RPKM values

[3] Adjusted p-value for multiple testing using Benjamini & Hochberg's method

**Supplementary Table 7B: Expression levels of candidate genes at the 22q11.231 locus in different cardiac tissues**

| | GTex Portal[1] | | Heinig et al. [2] | | | |
|---|---|---|---|---|---|---|
| | Heart Left Ventricle | Heart Atrial Appendage | Heart donors | DCM patients | Fold-change | P[3] |
| Gene | | | | | | |
| *RAB36* | 0.37 | 1.15 | NA | NA | NA | NA |
| *BCR* | 5.37 | 7.65 | 0.63 | 0.62 | -0.01 | 0.52 |
| *FBXW4P1* | 0.33 | 0.89 | NA | NA | NA | NA |
| *ZDHHC8P1* | 0.03 | 0.2 | NA | NA | NA | NA |
| *IGLL1* | 0 | 0 | NA | NA | NA | NA |
| *DRICH1* | 0.19 | 0.37 | NA | NA | NA | NA |
| *RGL4* | 0.46 | 0.88 | NA | NA | NA | NA |
| *ZNF70* | 0.819 | 1.58 | 0.20 | 0.30 | 0.1 | $2.6\ 10^{-19}$ |
| *VPREB3* | 0 | 0 | NA | NA | NA | NA |
| *C22orf15* | 0 | 0 | NA | NA | NA | NA |
| *CHCHD10* | 245.6 | 221.80 | 2.36 | 2.20 | -0.16 | $7.8\ 10^{-22}$ |
| *MMP11* | 0 | 0 | 0.40 | 0.32 | -0.08 | $2.1\ 10^{-5}$ |
| *SMARCB1* | 18.82 | 26.44 | 1.23 | 1.19 | -0.04 | $1.2\ 10^{-5}$ |
| *DERL3* | 0 | 3.486 | 0.26 | 0.21 | -0.05 | $2.8\ 10^{-5}$ |
| *SLC2A11* | 6.27 | 6.479 | 0.66 | 0.74 | 0.078 | $3.6\ 10^{-4}$ |
| *MIF* | 1.48 | 1.04 | 1.01 | 0.94 | -0.07 | $1.8\ 10^{-6}$ |
| *GSTT2* | 1.09 | 2.94 | 0.39 | 0.37 | -0.02 | 0.44 |
| *DDTL* | 2.56 | 2.5 | 0.61 | 0.54 | -0.07 | $1.5\ 10^{-7}$ |
| *DDT* | 24.18 | 28.48 | 1.25 | 1.15 | -0.1 | $7.5\ 10^{-12}$ |
| *GSTT1* | 25.77 | 43.65 | 0.89 | 0.89 | 0.00 | 0.51 |
| *CABIN1* | 9.37 | 13.73 | 0.89 | 0.91 | 0.02 | $5.0\ 10^{-3}$ |

[1] Expression values are shown in TPM (transcript per million) calculated from a gene model with isoforms collapsed to a single gene. No other normalization steps have been performed

[2] In Heinig et al, left ventricle gene expression levels were expressed as RPKM values

[3] Adjusted p-value for multiple testing using Benjamini & Hochberg's method

**Supplementary table 8. 4C interaction p-Value at chr22q11.23**. DNA sequence segments location is based on GRCh37.p13

| Chromosome | Start position | End position | P-value |
|---|---|---|---|
| chr22 | 24132984 | 24159004 | 9.33E-55 |
| chr22 | 24132217 | 24158653 | 2.70E-52 |
| chr22 | 24132988 | 24159063 | 2.70E-52 |
| chr22 | 24133200 | 24159514 | 2.70E-52 |
| chr22 | 24146194 | 24170051 | 2.70E-52 |
| chr22 | 24146281 | 24170985 | 2.70E-52 |
| chr22 | 24148050 | 24171300 | 2.70E-52 |
| chr22 | 24148471 | 24171587 | 2.70E-52 |
| chr22 | 24148713 | 24172489 | 2.70E-52 |
| chr22 | 24149362 | 24172519 | 2.70E-52 |
| chr22 | 24149550 | 24173480 | 2.70E-52 |
| chr22 | 24153559 | 24175173 | 2.70E-52 |
| chr22 | 24153677 | 24175343 | 2.70E-52 |
| chr22 | 24154469 | 24175867 | 2.70E-52 |
| chr22 | 24154658 | 24176122 | 2.70E-52 |
| chr22 | 24154920 | 24176480 | 2.70E-52 |
| chr22 | 24154925 | 24177458 | 2.70E-52 |
| chr22 | 24155576 | 24177517 | 2.70E-52 |
| chr22 | 24157859 | 24179207 | 2.70E-52 |
| chr22 | 24158426 | 24179645 | 2.70E-52 |
| chr22 | 24158433 | 24180373 | 2.70E-52 |
| chr22 | 24159514 | 24182979 | 2.70E-52 |
| chr22 | 24161134 | 24183968 | 2.70E-52 |
| chr22 | 24131766 | 24158433 | 6.62E-50 |
| chr22 | 24133274 | 24159736 | 6.62E-50 |

| | | | |
|---|---|---|---|
| chr22 | 24133641 | 24161134 | 6.62E-50 |
| chr22 | 24134927 | 24161380 | 6.62E-50 |
| chr22 | 24135748 | 24161584 | 6.62E-50 |
| chr22 | 24145509 | 24168381 | 6.62E-50 |
| chr22 | 24145566 | 24169055 | 6.62E-50 |
| chr22 | 24145871 | 24169339 | 6.62E-50 |
| chr22 | 24156806 | 24177529 | 6.62E-50 |
| chr22 | 24158653 | 24181529 | 6.62E-50 |
| chr22 | 24159004 | 24182342 | 6.62E-50 |
| chr22 | 24159063 | 24182380 | 6.62E-50 |
| chr22 | 24159736 | 24183544 | 6.62E-50 |
| chr22 | 24161380 | 24185082 | 6.62E-50 |
| chr22 | 24165898 | 24192938 | 6.62E-50 |
| chr22 | 24166378 | 24193701 | 6.62E-50 |
| chr22 | 24166479 | 24193804 | 6.62E-50 |
| chr22 | 24166893 | 24194231 | 6.62E-50 |
| chr22 | 24167610 | 24194544 | 6.62E-50 |
| chr22 | 24168172 | 24194658 | 6.62E-50 |
| chr22 | 24168381 | 24198647 | 6.62E-50 |
| chr22 | 24130636 | 24157859 | 1.40E-47 |
| chr22 | 24131562 | 24158426 | 1.40E-47 |
| chr22 | 24138952 | 24161879 | 1.40E-47 |
| chr22 | 24140321 | 24161913 | 1.40E-47 |
| chr22 | 24140765 | 24164229 | 1.40E-47 |
| chr22 | 24142606 | 24165898 | 1.40E-47 |
| chr22 | 24142688 | 24166378 | 1.40E-47 |
| chr22 | 24143086 | 24166479 | 1.40E-47 |
| chr22 | 24143940 | 24166893 | 1.40E-47 |
| chr22 | 24144129 | 24167610 | 1.40E-47 |

| | | | |
|---|---|---|---|
| chr22 | 24144334 | 24168172 | 1.40E-47 |
| chr22 | 24161584 | 24187767 | 1.40E-47 |
| chr22 | 24161879 | 24187934 | 1.40E-47 |
| chr22 | 24161913 | 24188295 | 1.40E-47 |
| chr22 | 24164229 | 24190916 | 1.40E-47 |
| chr22 | 24164871 | 24191163 | 1.40E-47 |
| chr22 | 24169055 | 24199084 | 1.40E-47 |
| chr22 | 24123957 | 24149550 | 2.58E-45 |
| chr22 | 24124537 | 24153559 | 2.58E-45 |
| chr22 | 24124645 | 24153677 | 2.58E-45 |
| chr22 | 24125457 | 24154469 | 2.58E-45 |
| chr22 | 24126373 | 24154658 | 2.58E-45 |
| chr22 | 24127977 | 24154920 | 2.58E-45 |
| chr22 | 24129316 | 24154925 | 2.58E-45 |
| chr22 | 24129440 | 24155576 | 2.58E-45 |
| chr22 | 24130034 | 24156806 | 2.58E-45 |
| chr22 | 24142441 | 24164871 | 2.58E-45 |
| chr22 | 24169339 | 24200209 | 2.58E-45 |
| chr22 | 24170051 | 24200480 | 2.58E-45 |
| chr22 | 24170985 | 24200803 | 2.58E-45 |
| chr22 | 24123679 | 24149362 | 4.19E-43 |
| chr22 | 24171300 | 24200830 | 4.19E-43 |
| chr22 | 24171587 | 24201230 | 4.19E-43 |
| chr22 | 24172489 | 24204305 | 4.19E-43 |
| chr22 | 24172519 | 24204785 | 4.19E-43 |
| chr22 | 24121703 | 24146194 | 6.03E-41 |
| chr22 | 24122549 | 24146281 | 6.03E-41 |
| chr22 | 24122678 | 24148050 | 6.03E-41 |
| chr22 | 24122946 | 24148471 | 6.03E-41 |

| | | | |
|---|---|---|---|
| chr22 | 24123616 | 24148713 | 6.03E-41 |
| chr22 | 24173480 | 24206321 | 6.03E-41 |
| chr22 | 24120647 | 24145566 | 7.73E-39 |
| chr22 | 24120902 | 24145871 | 7.73E-39 |
| chr22 | 24175173 | 24206335 | 7.73E-39 |
| chr22 | 24120640 | 24145509 | 8.88E-37 |
| chr22 | 24175343 | 24206483 | 8.88E-37 |
| chr22 | 24120159 | 24144129 | 9.16E-35 |
| chr22 | 24120164 | 24144334 | 9.16E-35 |
| chr22 | 24175867 | 24207176 | 9.16E-35 |
| chr22 | 24118851 | 24143086 | 8.52E-33 |
| chr22 | 24120131 | 24143940 | 8.52E-33 |
| chr22 | 24176122 | 24207822 | 8.52E-33 |
| chr22 | 24036459 | 24056605 | 7.16E-31 |
| chr22 | 24037009 | 24056987 | 7.16E-31 |
| chr22 | 24037203 | 24057259 | 7.16E-31 |
| chr22 | 24118265 | 24142688 | 7.16E-31 |
| chr22 | 24176480 | 24207950 | 7.16E-31 |
| chr22 | 24036239 | 24056214 | 5.45E-29 |
| chr22 | 24037563 | 24057429 | 5.45E-29 |
| chr22 | 24038102 | 24059800 | 5.45E-29 |
| chr22 | 24038725 | 24061180 | 5.45E-29 |
| chr22 | 24038913 | 24063321 | 5.45E-29 |
| chr22 | 24039471 | 24063634 | 5.45E-29 |
| chr22 | 24039514 | 24064468 | 5.45E-29 |
| chr22 | 24041211 | 24064517 | 5.45E-29 |
| chr22 | 24116312 | 24142606 | 5.45E-29 |
| chr22 | 24177458 | 24208247 | 5.45E-29 |
| chr22 | 24177517 | 24209120 | 5.45E-29 |

| | | | |
|---|---|---|---|
| chr22 | 24177529 | 24209344 | 5.45E-29 |
| chr22 | 24179207 | 24210021 | 5.45E-29 |
| chr22 | 24179645 | 24210153 | 5.45E-29 |
| chr22 | 24033288 | 24052815 | 3.76E-27 |
| chr22 | 24034207 | 24054278 | 3.76E-27 |
| chr22 | 24034858 | 24054617 | 3.76E-27 |
| chr22 | 24035300 | 24054906 | 3.76E-27 |
| chr22 | 24036149 | 24056017 | 3.76E-27 |
| chr22 | 24037936 | 24058181 | 3.76E-27 |
| chr22 | 24037951 | 24059757 | 3.76E-27 |
| chr22 | 24038671 | 24060427 | 3.76E-27 |
| chr22 | 24041243 | 24064770 | 3.76E-27 |
| chr22 | 24043098 | 24070033 | 3.76E-27 |
| chr22 | 24097503 | 24125457 | 3.76E-27 |
| chr22 | 24098262 | 24126373 | 3.76E-27 |
| chr22 | 24099716 | 24127977 | 3.76E-27 |
| chr22 | 24100440 | 24129316 | 3.76E-27 |
| chr22 | 24101863 | 24129440 | 3.76E-27 |
| chr22 | 24105822 | 24130636 | 3.76E-27 |
| chr22 | 24106243 | 24131562 | 3.76E-27 |
| chr22 | 24106580 | 24131766 | 3.76E-27 |
| chr22 | 24108493 | 24133274 | 3.76E-27 |
| chr22 | 24108598 | 24133641 | 3.76E-27 |
| chr22 | 24113446 | 24135748 | 3.76E-27 |
| chr22 | 24113571 | 24138952 | 3.76E-27 |
| chr22 | 24114596 | 24140321 | 3.76E-27 |
| chr22 | 24115240 | 24140765 | 3.76E-27 |
| chr22 | 24115536 | 24142441 | 3.76E-27 |
| chr22 | 24180373 | 24210728 | 3.76E-27 |

| | | | |
|---|---|---|---|
| chr22 | 24182342 | 24212288 | 3.76E-27 |
| chr22 | 24182380 | 24214618 | 3.76E-27 |
| chr22 | 24223878 | 24244122 | 3.76E-27 |
| chr22 | 24225014 | 24245643 | 3.76E-27 |
| chr22 | 24226505 | 24246101 | 3.76E-27 |
| chr22 | 24226577 | 24246405 | 3.76E-27 |
| chr22 | 24226946 | 24246502 | 3.76E-27 |
| chr22 | 24227319 | 24246743 | 3.76E-27 |
| chr22 | 24235698 | 24252017 | 3.76E-27 |
| chr22 | 24237863 | 24253048 | 3.76E-27 |
| chr22 | 24238600 | 24253145 | 3.76E-27 |
| chr22 | 24238918 | 24253627 | 3.76E-27 |
| chr22 | 24032975 | 24052373 | 2.35E-25 |
| chr22 | 24041866 | 24064840 | 2.35E-25 |
| chr22 | 24043417 | 24070075 | 2.35E-25 |
| chr22 | 24044578 | 24072034 | 2.35E-25 |
| chr22 | 24045934 | 24072864 | 2.35E-25 |
| chr22 | 24096998 | 24124645 | 2.35E-25 |
| chr22 | 24105459 | 24130034 | 2.35E-25 |
| chr22 | 24107182 | 24132217 | 2.35E-25 |
| chr22 | 24107561 | 24132984 | 2.35E-25 |
| chr22 | 24107808 | 24132988 | 2.35E-25 |
| chr22 | 24108409 | 24133200 | 2.35E-25 |
| chr22 | 24112863 | 24134927 | 2.35E-25 |
| chr22 | 24181529 | 24210938 | 2.35E-25 |
| chr22 | 24182979 | 24214720 | 2.35E-25 |
| chr22 | 24188295 | 24217394 | 2.35E-25 |
| chr22 | 24190916 | 24217772 | 2.35E-25 |
| chr22 | 24191163 | 24218153 | 2.35E-25 |

| | | | |
|---|---|---|---|
| chr22 | 24223410 | 24244067 | 2.35E-25 |
| chr22 | 24223458 | 24244076 | 2.35E-25 |
| chr22 | 24225002 | 24245354 | 2.35E-25 |
| chr22 | 24228129 | 24247076 | 2.35E-25 |
| chr22 | 24233213 | 24251977 | 2.35E-25 |
| chr22 | 24237054 | 24252122 | 2.35E-25 |
| chr22 | 24237666 | 24252557 | 2.35E-25 |
| chr22 | 24239776 | 24253678 | 2.35E-25 |
| chr22 | 24239986 | 24253870 | 2.35E-25 |
| chr22 | 24240686 | 24257101 | 2.35E-25 |
| chr22 | 24241262 | 24257513 | 2.35E-25 |
| chr22 | 24241291 | 24257890 | 2.35E-25 |
| chr22 | 24242887 | 24259655 | 2.35E-25 |
| chr22 | 24243117 | 24259858 | 2.35E-25 |
| chr22 | 24243393 | 24260557 | 2.35E-25 |
| chr22 | 24032905 | 24051787 | 1.33E-23 |
| chr22 | 24046394 | 24073381 | 1.33E-23 |
| chr22 | 24046640 | 24073461 | 1.33E-23 |
| chr22 | 24047138 | 24075704 | 1.33E-23 |
| chr22 | 24094910 | 24122946 | 1.33E-23 |
| chr22 | 24095182 | 24123616 | 1.33E-23 |
| chr22 | 24095209 | 24123679 | 1.33E-23 |
| chr22 | 24095640 | 24123957 | 1.33E-23 |
| chr22 | 24095978 | 24124537 | 1.33E-23 |
| chr22 | 24183544 | 24215818 | 1.33E-23 |
| chr22 | 24183968 | 24216012 | 1.33E-23 |
| chr22 | 24185082 | 24216167 | 1.33E-23 |
| chr22 | 24187767 | 24216190 | 1.33E-23 |
| chr22 | 24187934 | 24217005 | 1.33E-23 |

| | | | |
|---|---|---|---|
| chr22 | 24192938 | 24218240 | 1.33E-23 |
| chr22 | 24193701 | 24218276 | 1.33E-23 |
| chr22 | 24223272 | 24243117 | 1.33E-23 |
| chr22 | 24223301 | 24243393 | 1.33E-23 |
| chr22 | 24223378 | 24243825 | 1.33E-23 |
| chr22 | 24229039 | 24247432 | 1.33E-23 |
| chr22 | 24232691 | 24251449 | 1.33E-23 |
| chr22 | 24232700 | 24251867 | 1.33E-23 |
| chr22 | 24239992 | 24255960 | 1.33E-23 |
| chr22 | 24240678 | 24257035 | 1.33E-23 |
| chr22 | 24243825 | 24262669 | 1.33E-23 |
| chr22 | 24032452 | 24050585 | 6.86E-22 |
| chr22 | 24032899 | 24050755 | 6.86E-22 |
| chr22 | 24046754 | 24074327 | 6.86E-22 |
| chr22 | 24047006 | 24074971 | 6.86E-22 |
| chr22 | 24047556 | 24076059 | 6.86E-22 |
| chr22 | 24093741 | 24122678 | 6.86E-22 |
| chr22 | 24193804 | 24218433 | 6.86E-22 |
| chr22 | 24217394 | 24231067 | 6.86E-22 |
| chr22 | 24217772 | 24231329 | 6.86E-22 |
| chr22 | 24218153 | 24231453 | 6.86E-22 |
| chr22 | 24218240 | 24232050 | 6.86E-22 |
| chr22 | 24218276 | 24232071 | 6.86E-22 |
| chr22 | 24218817 | 24232576 | 6.86E-22 |
| chr22 | 24219001 | 24232640 | 6.86E-22 |
| chr22 | 24219046 | 24232691 | 6.86E-22 |
| chr22 | 24221877 | 24241262 | 6.86E-22 |
| chr22 | 24222792 | 24241291 | 6.86E-22 |
| chr22 | 24222837 | 24242887 | 6.86E-22 |

| | | | |
|---|---|---|---|
| chr22 | 24231067 | 24247769 | 6.86E-22 |
| chr22 | 24231329 | 24247933 | 6.86E-22 |
| chr22 | 24232050 | 24249032 | 6.86E-22 |
| chr22 | 24232182 | 24249397 | 6.86E-22 |
| chr22 | 24232576 | 24250706 | 6.86E-22 |
| chr22 | 24232640 | 24250938 | 6.86E-22 |
| chr22 | 24244067 | 24262761 | 6.86E-22 |
| chr22 | 24031391 | 24049474 | 3.20E-20 |
| chr22 | 24048304 | 24077161 | 3.20E-20 |
| chr22 | 24048376 | 24077465 | 3.20E-20 |
| chr22 | 24049098 | 24078021 | 3.20E-20 |
| chr22 | 24049474 | 24078158 | 3.20E-20 |
| chr22 | 24092877 | 24122549 | 3.20E-20 |
| chr22 | 24194231 | 24218817 | 3.20E-20 |
| chr22 | 24210153 | 24223410 | 3.20E-20 |
| chr22 | 24210938 | 24223878 | 3.20E-20 |
| chr22 | 24212288 | 24225002 | 3.20E-20 |
| chr22 | 24217005 | 24229039 | 3.20E-20 |
| chr22 | 24218433 | 24232182 | 3.20E-20 |
| chr22 | 24219101 | 24232700 | 3.20E-20 |
| chr22 | 24219120 | 24233213 | 3.20E-20 |
| chr22 | 24219332 | 24235698 | 3.20E-20 |
| chr22 | 24221048 | 24239986 | 3.20E-20 |
| chr22 | 24221686 | 24240686 | 3.20E-20 |
| chr22 | 24231453 | 24248040 | 3.20E-20 |
| chr22 | 24232071 | 24249095 | 3.20E-20 |
| chr22 | 24244076 | 24262920 | 3.20E-20 |
| chr22 | 24029778 | 24048376 | 1.35E-18 |
| chr22 | 24030729 | 24049098 | 1.35E-18 |

| | | | |
|---|---|---|---|
| chr22 | 24050585 | 24078189 | 1.35E-18 |
| chr22 | 24087178 | 24118851 | 1.35E-18 |
| chr22 | 24087190 | 24120131 | 1.35E-18 |
| chr22 | 24087536 | 24120159 | 1.35E-18 |
| chr22 | 24092448 | 24121703 | 1.35E-18 |
| chr22 | 24194544 | 24219001 | 1.35E-18 |
| chr22 | 24194658 | 24219046 | 1.35E-18 |
| chr22 | 24200209 | 24219332 | 1.35E-18 |
| chr22 | 24200480 | 24219762 | 1.35E-18 |
| chr22 | 24200803 | 24220192 | 1.35E-18 |
| chr22 | 24201230 | 24220512 | 1.35E-18 |
| chr22 | 24204305 | 24220963 | 1.35E-18 |
| chr22 | 24209120 | 24223272 | 1.35E-18 |
| chr22 | 24209344 | 24223301 | 1.35E-18 |
| chr22 | 24210021 | 24223378 | 1.35E-18 |
| chr22 | 24210728 | 24223458 | 1.35E-18 |
| chr22 | 24214618 | 24225014 | 1.35E-18 |
| chr22 | 24216167 | 24227319 | 1.35E-18 |
| chr22 | 24216190 | 24228129 | 1.35E-18 |
| chr22 | 24219762 | 24237054 | 1.35E-18 |
| chr22 | 24220192 | 24237666 | 1.35E-18 |
| chr22 | 24220512 | 24238600 | 1.35E-18 |
| chr22 | 24220963 | 24238918 | 1.35E-18 |
| chr22 | 24221010 | 24239776 | 1.35E-18 |
| chr22 | 24221063 | 24239992 | 1.35E-18 |
| chr22 | 24221417 | 24240678 | 1.35E-18 |
| chr22 | 24244122 | 24265978 | 1.35E-18 |
| chr22 | 23938536 | 23960292 | 5.15E-17 |
| chr22 | 24029271 | 24048304 | 5.15E-17 |

| | | | |
|---|---|---|---|
| chr22 | 24050755 | 24078215 | 5.15E-17 |
| chr22 | 24086481 | 24114596 | 5.15E-17 |
| chr22 | 24086512 | 24115240 | 5.15E-17 |
| chr22 | 24086680 | 24115536 | 5.15E-17 |
| chr22 | 24086767 | 24116312 | 5.15E-17 |
| chr22 | 24086848 | 24118265 | 5.15E-17 |
| chr22 | 24088266 | 24120164 | 5.15E-17 |
| chr22 | 24088715 | 24120640 | 5.15E-17 |
| chr22 | 24089700 | 24120647 | 5.15E-17 |
| chr22 | 24091948 | 24120902 | 5.15E-17 |
| chr22 | 24198647 | 24219101 | 5.15E-17 |
| chr22 | 24199084 | 24219120 | 5.15E-17 |
| chr22 | 24200830 | 24220261 | 5.15E-17 |
| chr22 | 24204785 | 24221010 | 5.15E-17 |
| chr22 | 24206321 | 24221048 | 5.15E-17 |
| chr22 | 24206335 | 24221063 | 5.15E-17 |
| chr22 | 24206483 | 24221417 | 5.15E-17 |
| chr22 | 24207176 | 24221686 | 5.15E-17 |
| chr22 | 24207822 | 24221877 | 5.15E-17 |
| chr22 | 24207950 | 24222792 | 5.15E-17 |
| chr22 | 24208247 | 24222837 | 5.15E-17 |
| chr22 | 24214720 | 24226505 | 5.15E-17 |
| chr22 | 24215818 | 24226577 | 5.15E-17 |
| chr22 | 24216012 | 24226946 | 5.15E-17 |
| chr22 | 24220261 | 24237863 | 5.15E-17 |
| chr22 | 24245354 | 24266125 | 5.15E-17 |
| chr22 | 24245643 | 24267564 | 5.15E-17 |
| chr22 | 24246502 | 24270090 | 5.15E-17 |
| chr22 | 24246743 | 24270774 | 5.15E-17 |

| | | | |
|---|---|---|---|
| chr22 | 24247076 | 24271391 | 5.15E-17 |
| chr22 | 24247432 | 24271686 | 5.15E-17 |
| chr22 | 24247769 | 24271892 | 5.15E-17 |
| chr22 | 24247933 | 24272066 | 5.15E-17 |
| chr22 | 23869293 | 23897208 | 1.77E-15 |
| chr22 | 23929540 | 23956425 | 1.77E-15 |
| chr22 | 23930122 | 23956769 | 1.77E-15 |
| chr22 | 23934332 | 23959095 | 1.77E-15 |
| chr22 | 23935783 | 23959100 | 1.77E-15 |
| chr22 | 23936934 | 23959127 | 1.77E-15 |
| chr22 | 23937006 | 23959868 | 1.77E-15 |
| chr22 | 23940472 | 23960604 | 1.77E-15 |
| chr22 | 23941210 | 23960628 | 1.77E-15 |
| chr22 | 23953409 | 23973815 | 1.77E-15 |
| chr22 | 24029113 | 24047556 | 1.77E-15 |
| chr22 | 24051787 | 24079813 | 1.77E-15 |
| chr22 | 24086428 | 24113571 | 1.77E-15 |
| chr22 | 24246101 | 24268180 | 1.77E-15 |
| chr22 | 24246405 | 24269411 | 1.77E-15 |
| chr22 | 24248040 | 24274573 | 1.77E-15 |
| chr22 | 24249032 | 24274695 | 1.77E-15 |
| chr22 | 24471230 | 24497781 | 1.77E-15 |
| chr22 | 24475389 | 24500121 | 1.77E-15 |
| chr22 | 24475995 | 24501051 | 1.77E-15 |
| chr22 | 24476339 | 24501191 | 1.77E-15 |
| chr22 | 24477537 | 24501943 | 1.77E-15 |
| chr22 | 23867319 | 23892023 | 5.49E-14 |
| chr22 | 23868732 | 23894004 | 5.49E-14 |
| chr22 | 23869134 | 23896131 | 5.49E-14 |

| | | | |
|---|---|---|---|
| chr22 | 23869264 | 23896219 | 5.49E-14 |
| chr22 | 23869740 | 23898226 | 5.49E-14 |
| chr22 | 23870072 | 23898455 | 5.49E-14 |
| chr22 | 23870385 | 23898476 | 5.49E-14 |
| chr22 | 23926634 | 23955641 | 5.49E-14 |
| chr22 | 23926740 | 23955909 | 5.49E-14 |
| chr22 | 23930139 | 23956902 | 5.49E-14 |
| chr22 | 23931234 | 23957684 | 5.49E-14 |
| chr22 | 23933831 | 23958520 | 5.49E-14 |
| chr22 | 23941766 | 23960742 | 5.49E-14 |
| chr22 | 23941966 | 23960794 | 5.49E-14 |
| chr22 | 23942095 | 23961539 | 5.49E-14 |
| chr22 | 23950619 | 23970891 | 5.49E-14 |
| chr22 | 23952657 | 23971653 | 5.49E-14 |
| chr22 | 23952672 | 23971714 | 5.49E-14 |
| chr22 | 23952739 | 23972046 | 5.49E-14 |
| chr22 | 23955641 | 23974335 | 5.49E-14 |
| chr22 | 23955909 | 23976670 | 5.49E-14 |
| chr22 | 23956425 | 23977591 | 5.49E-14 |
| chr22 | 23956769 | 23978140 | 5.49E-14 |
| chr22 | 23956902 | 23987879 | 5.49E-14 |
| chr22 | 23957684 | 23988392 | 5.49E-14 |
| chr22 | 23958055 | 23988819 | 5.49E-14 |
| chr22 | 23958488 | 23990248 | 5.49E-14 |
| chr22 | 23958520 | 23991438 | 5.49E-14 |
| chr22 | 24028149 | 24047138 | 5.49E-14 |
| chr22 | 24052373 | 24080082 | 5.49E-14 |
| chr22 | 24086191 | 24113446 | 5.49E-14 |
| chr22 | 24249095 | 24274702 | 5.49E-14 |

| | | | |
|---|---|---|---|
| chr22 | 24249397 | 24274871 | 5.49E-14 |
| chr22 | 24470506 | 24497220 | 5.49E-14 |
| chr22 | 24472504 | 24498633 | 5.49E-14 |
| chr22 | 24473276 | 24499818 | 5.49E-14 |
| chr22 | 24478668 | 24501952 | 5.49E-14 |
| chr22 | 24480538 | 24503604 | 5.49E-14 |
| chr22 | 24481119 | 24503697 | 5.49E-14 |
| chr22 | 23635952 | 23683804 | 1.52E-12 |
| chr22 | 23636237 | 23685141 | 1.52E-12 |
| chr22 | 23636570 | 23685576 | 1.52E-12 |
| chr22 | 23640487 | 23696928 | 1.52E-12 |
| chr22 | 23640577 | 23697027 | 1.52E-12 |
| chr22 | 23642703 | 23698156 | 1.52E-12 |
| chr22 | 23643902 | 23698732 | 1.52E-12 |
| chr22 | 23644463 | 23699103 | 1.52E-12 |
| chr22 | 23861713 | 23885481 | 1.52E-12 |
| chr22 | 23862205 | 23887056 | 1.52E-12 |
| chr22 | 23863243 | 23887496 | 1.52E-12 |
| chr22 | 23863349 | 23887960 | 1.52E-12 |
| chr22 | 23866403 | 23891708 | 1.52E-12 |
| chr22 | 23866606 | 23891793 | 1.52E-12 |
| chr22 | 23870521 | 23898552 | 1.52E-12 |
| chr22 | 23870649 | 23898641 | 1.52E-12 |
| chr22 | 23871124 | 23898942 | 1.52E-12 |
| chr22 | 23871608 | 23899279 | 1.52E-12 |
| chr22 | 23926274 | 23953409 | 1.52E-12 |
| chr22 | 23932177 | 23958055 | 1.52E-12 |
| chr22 | 23933047 | 23958488 | 1.52E-12 |
| chr22 | 23942137 | 23962441 | 1.52E-12 |

| | | | |
|---|---|---|---|
| chr22 | 23942611 | 23962868 | 1.52E-12 |
| chr22 | 23947136 | 23965152 | 1.52E-12 |
| chr22 | 23947274 | 23965373 | 1.52E-12 |
| chr22 | 23948902 | 23966046 | 1.52E-12 |
| chr22 | 23949260 | 23966540 | 1.52E-12 |
| chr22 | 23949562 | 23967157 | 1.52E-12 |
| chr22 | 23949915 | 23969888 | 1.52E-12 |
| chr22 | 23950564 | 23970708 | 1.52E-12 |
| chr22 | 23959095 | 23992024 | 1.52E-12 |
| chr22 | 24027298 | 24045934 | 1.52E-12 |
| chr22 | 24027465 | 24046394 | 1.52E-12 |
| chr22 | 24027674 | 24046640 | 1.52E-12 |
| chr22 | 24027713 | 24046754 | 1.52E-12 |
| chr22 | 24027820 | 24047006 | 1.52E-12 |
| chr22 | 24052815 | 24081033 | 1.52E-12 |
| chr22 | 24054278 | 24081103 | 1.52E-12 |
| chr22 | 24054617 | 24081151 | 1.52E-12 |
| chr22 | 24054906 | 24082596 | 1.52E-12 |
| chr22 | 24056017 | 24083082 | 1.52E-12 |
| chr22 | 24056214 | 24083216 | 1.52E-12 |
| chr22 | 24056605 | 24083454 | 1.52E-12 |
| chr22 | 24083967 | 24108598 | 1.52E-12 |
| chr22 | 24085300 | 24112863 | 1.52E-12 |
| chr22 | 24250706 | 24275028 | 1.52E-12 |
| chr22 | 24250938 | 24276281 | 1.52E-12 |
| chr22 | 24380611 | 24412578 | 1.52E-12 |
| chr22 | 24383880 | 24412693 | 1.52E-12 |
| chr22 | 24384132 | 24417404 | 1.52E-12 |
| chr22 | 24469564 | 24497064 | 1.52E-12 |

| | | | |
|---|---|---|---|
| chr22 | 24479906 | 24502940 | 1.52E-12 |
| chr22 | 24481958 | 24504069 | 1.52E-12 |
| chr22 | 24482460 | 24504167 | 1.52E-12 |
| chr22 | 24482666 | 24504190 | 1.52E-12 |
| chr22 | 24486451 | 24507015 | 1.52E-12 |
| chr22 | 24486779 | 24508883 | 1.52E-12 |
| chr22 | 23631876 | 23646984 | 3.76E-11 |
| chr22 | 23632097 | 23647332 | 3.76E-11 |
| chr22 | 23632340 | 23650849 | 3.76E-11 |
| chr22 | 23632824 | 23658051 | 3.76E-11 |
| chr22 | 23632859 | 23664624 | 3.76E-11 |
| chr22 | 23634730 | 23666235 | 3.76E-11 |
| chr22 | 23635394 | 23667211 | 3.76E-11 |
| chr22 | 23635630 | 23675321 | 3.76E-11 |
| chr22 | 23635769 | 23680649 | 3.76E-11 |
| chr22 | 23636941 | 23688050 | 3.76E-11 |
| chr22 | 23637288 | 23690635 | 3.76E-11 |
| chr22 | 23640447 | 23695866 | 3.76E-11 |
| chr22 | 23640451 | 23695938 | 3.76E-11 |
| chr22 | 23644576 | 23699269 | 3.76E-11 |
| chr22 | 23644930 | 23699281 | 3.76E-11 |
| chr22 | 23861461 | 23883701 | 3.76E-11 |
| chr22 | 23861497 | 23883735 | 3.76E-11 |
| chr22 | 23863411 | 23888787 | 3.76E-11 |
| chr22 | 23863868 | 23891005 | 3.76E-11 |
| chr22 | 23865373 | 23891077 | 3.76E-11 |
| chr22 | 23865939 | 23891583 | 3.76E-11 |
| chr22 | 23871645 | 23899397 | 3.76E-11 |
| chr22 | 23913703 | 23945673 | 3.76E-11 |

| | | | |
|---|---|---|---|
| chr22 | 23914383 | 23945713 | 3.76E-11 |
| chr22 | 23914434 | 23945862 | 3.76E-11 |
| chr22 | 23917733 | 23945942 | 3.76E-11 |
| chr22 | 23920310 | 23946150 | 3.76E-11 |
| chr22 | 23920604 | 23947136 | 3.76E-11 |
| chr22 | 23921221 | 23947274 | 3.76E-11 |
| chr22 | 23921352 | 23948902 | 3.76E-11 |
| chr22 | 23922612 | 23949260 | 3.76E-11 |
| chr22 | 23923304 | 23949562 | 3.76E-11 |
| chr22 | 23924979 | 23949915 | 3.76E-11 |
| chr22 | 23925358 | 23950564 | 3.76E-11 |
| chr22 | 23925462 | 23950619 | 3.76E-11 |
| chr22 | 23925721 | 23952657 | 3.76E-11 |
| chr22 | 23925767 | 23952672 | 3.76E-11 |
| chr22 | 23925833 | 23952739 | 3.76E-11 |
| chr22 | 23943909 | 23963777 | 3.76E-11 |
| chr22 | 23945673 | 23964387 | 3.76E-11 |
| chr22 | 23946150 | 23965029 | 3.76E-11 |
| chr22 | 23959100 | 23992032 | 3.76E-11 |
| chr22 | 23959127 | 23993013 | 3.76E-11 |
| chr22 | 23959868 | 23993318 | 3.76E-11 |
| chr22 | 23992032 | 24025760 | 3.76E-11 |
| chr22 | 23993013 | 24025854 | 3.76E-11 |
| chr22 | 23993318 | 24026498 | 3.76E-11 |
| chr22 | 23993942 | 24026982 | 3.76E-11 |
| chr22 | 23994960 | 24027054 | 3.76E-11 |
| chr22 | 24026982 | 24043098 | 3.76E-11 |
| chr22 | 24027054 | 24043417 | 3.76E-11 |
| chr22 | 24027172 | 24044578 | 3.76E-11 |

| | | | |
|---|---|---|---|
| chr22 | 24056987 | 24083830 | 3.76E-11 |
| chr22 | 24057259 | 24083967 | 3.76E-11 |
| chr22 | 24083216 | 24107808 | 3.76E-11 |
| chr22 | 24083454 | 24108409 | 3.76E-11 |
| chr22 | 24083830 | 24108493 | 3.76E-11 |
| chr22 | 24251449 | 24276761 | 3.76E-11 |
| chr22 | 24338640 | 24368960 | 3.76E-11 |
| chr22 | 24359956 | 24393398 | 3.76E-11 |
| chr22 | 24360861 | 24393410 | 3.76E-11 |
| chr22 | 24376432 | 24410279 | 3.76E-11 |
| chr22 | 24376698 | 24410458 | 3.76E-11 |
| chr22 | 24378029 | 24410609 | 3.76E-11 |
| chr22 | 24378096 | 24410831 | 3.76E-11 |
| chr22 | 24380352 | 24411791 | 3.76E-11 |
| chr22 | 24383966 | 24415453 | 3.76E-11 |
| chr22 | 24384445 | 24417905 | 3.76E-11 |
| chr22 | 24385910 | 24418054 | 3.76E-11 |
| chr22 | 24387189 | 24420825 | 3.76E-11 |
| chr22 | 24387622 | 24420893 | 3.76E-11 |
| chr22 | 24387633 | 24421896 | 3.76E-11 |
| chr22 | 24387666 | 24422570 | 3.76E-11 |
| chr22 | 24468375 | 24496162 | 3.76E-11 |
| chr22 | 24468390 | 24496321 | 3.76E-11 |
| chr22 | 24469089 | 24496714 | 3.76E-11 |
| chr22 | 24482830 | 24504325 | 3.76E-11 |
| chr22 | 24482959 | 24504559 | 3.76E-11 |
| chr22 | 24484010 | 24504716 | 3.76E-11 |
| chr22 | 24484846 | 24505136 | 3.76E-11 |
| chr22 | 24485066 | 24506483 | 3.76E-11 |

| | | | |
|---|---|---|---|
| chr22 | 24485351 | 24506867 | 3.76E-11 |
| chr22 | 24487485 | 24509022 | 3.76E-11 |
| chr22 | 24487527 | 24509039 | 3.76E-11 |
| chr22 | 24487755 | 24509691 | 3.76E-11 |
| chr22 | 24488450 | 24510088 | 3.76E-11 |
| chr22 | 24489107 | 24510280 | 3.76E-11 |
| chr22 | 24495348 | 24512211 | 3.76E-11 |
| chr22 | 24495380 | 24512325 | 3.76E-11 |
| chr22 | 24495973 | 24512576 | 3.76E-11 |
| chr22 | 24496162 | 24512803 | 3.76E-11 |
| chr22 | 23568587 | 23589088 | 8.24E-10 |
| chr22 | 23569153 | 23589481 | 8.24E-10 |
| chr22 | 23569196 | 23589562 | 8.24E-10 |
| chr22 | 23569567 | 23589646 | 8.24E-10 |
| chr22 | 23569720 | 23589711 | 8.24E-10 |
| chr22 | 23569825 | 23589821 | 8.24E-10 |
| chr22 | 23569889 | 23591764 | 8.24E-10 |
| chr22 | 23570538 | 23591984 | 8.24E-10 |
| chr22 | 23571790 | 23592096 | 8.24E-10 |
| chr22 | 23571812 | 23592506 | 8.24E-10 |
| chr22 | 23571894 | 23593191 | 8.24E-10 |
| chr22 | 23571923 | 23593548 | 8.24E-10 |
| chr22 | 23630604 | 23645491 | 8.24E-10 |
| chr22 | 23631178 | 23645869 | 8.24E-10 |
| chr22 | 23631185 | 23646466 | 8.24E-10 |
| chr22 | 23632348 | 23655037 | 8.24E-10 |
| chr22 | 23632515 | 23657880 | 8.24E-10 |
| chr22 | 23634467 | 23665988 | 8.24E-10 |
| chr22 | 23637999 | 23690932 | 8.24E-10 |

| | | | |
|---|---|---|---|
| chr22 | 23638834 | 23694580 | 8.24E-10 |
| chr22 | 23639130 | 23694873 | 8.24E-10 |
| chr22 | 23639602 | 23695785 | 8.24E-10 |
| chr22 | 23645339 | 23699687 | 8.24E-10 |
| chr22 | 23645491 | 23700543 | 8.24E-10 |
| chr22 | 23756899 | 23785197 | 8.24E-10 |
| chr22 | 23757240 | 23785378 | 8.24E-10 |
| chr22 | 23767845 | 23792455 | 8.24E-10 |
| chr22 | 23770631 | 23792671 | 8.24E-10 |
| chr22 | 23770742 | 23794510 | 8.24E-10 |
| chr22 | 23770820 | 23794859 | 8.24E-10 |
| chr22 | 23771730 | 23795395 | 8.24E-10 |
| chr22 | 23772830 | 23795622 | 8.24E-10 |
| chr22 | 23773038 | 23796365 | 8.24E-10 |
| chr22 | 23773432 | 23797109 | 8.24E-10 |
| chr22 | 23773766 | 23797650 | 8.24E-10 |
| chr22 | 23860904 | 23880109 | 8.24E-10 |
| chr22 | 23860962 | 23881587 | 8.24E-10 |
| chr22 | 23861356 | 23883400 | 8.24E-10 |
| chr22 | 23872192 | 23899607 | 8.24E-10 |
| chr22 | 23872512 | 23900148 | 8.24E-10 |
| chr22 | 23872846 | 23900435 | 8.24E-10 |
| chr22 | 23873586 | 23901349 | 8.24E-10 |
| chr22 | 23876472 | 23902598 | 8.24E-10 |
| chr22 | 23876625 | 23903530 | 8.24E-10 |
| chr22 | 23876775 | 23904554 | 8.24E-10 |
| chr22 | 23879949 | 23905451 | 8.24E-10 |
| chr22 | 23880109 | 23906361 | 8.24E-10 |
| chr22 | 23913361 | 23943909 | 8.24E-10 |

| | | | |
|---|---|---|---|
| chr22 | 23945713 | 23964712 | 8.24E-10 |
| chr22 | 23945862 | 23964731 | 8.24E-10 |
| chr22 | 23945942 | 23964773 | 8.24E-10 |
| chr22 | 23960292 | 23993942 | 8.24E-10 |
| chr22 | 23960604 | 23994960 | 8.24E-10 |
| chr22 | 23960628 | 23995202 | 8.24E-10 |
| chr22 | 23960742 | 23995324 | 8.24E-10 |
| chr22 | 23960794 | 23995874 | 8.24E-10 |
| chr22 | 23961539 | 23996434 | 8.24E-10 |
| chr22 | 23964387 | 23998379 | 8.24E-10 |
| chr22 | 23970891 | 24016279 | 8.24E-10 |
| chr22 | 23971653 | 24016577 | 8.24E-10 |
| chr22 | 23971714 | 24016762 | 8.24E-10 |
| chr22 | 23972046 | 24016992 | 8.24E-10 |
| chr22 | 23973815 | 24017125 | 8.24E-10 |
| chr22 | 23974335 | 24018737 | 8.24E-10 |
| chr22 | 23976670 | 24018926 | 8.24E-10 |
| chr22 | 23991438 | 24023683 | 8.24E-10 |
| chr22 | 23992024 | 24025202 | 8.24E-10 |
| chr22 | 23995202 | 24027172 | 8.24E-10 |
| chr22 | 23995324 | 24027298 | 8.24E-10 |
| chr22 | 23995874 | 24027465 | 8.24E-10 |
| chr22 | 23996434 | 24027674 | 8.24E-10 |
| chr22 | 23997533 | 24027713 | 8.24E-10 |
| chr22 | 23998340 | 24027820 | 8.24E-10 |
| chr22 | 23998370 | 24028149 | 8.24E-10 |
| chr22 | 23998379 | 24029113 | 8.24E-10 |
| chr22 | 24002121 | 24032452 | 8.24E-10 |
| chr22 | 24022273 | 24038102 | 8.24E-10 |

| | | | |
|---|---|---|---|
| chr22 | 24023683 | 24039471 | 8.24E-10 |
| chr22 | 24025760 | 24041211 | 8.24E-10 |
| chr22 | 24025854 | 24041243 | 8.24E-10 |
| chr22 | 24026498 | 24041866 | 8.24E-10 |
| chr22 | 24057429 | 24085300 | 8.24E-10 |
| chr22 | 24058181 | 24086191 | 8.24E-10 |
| chr22 | 24059757 | 24086428 | 8.24E-10 |
| chr22 | 24083082 | 24107561 | 8.24E-10 |
| chr22 | 24251867 | 24276803 | 8.24E-10 |
| chr22 | 24251977 | 24276981 | 8.24E-10 |
| chr22 | 24338619 | 24366902 | 8.24E-10 |
| chr22 | 24338898 | 24369975 | 8.24E-10 |
| chr22 | 24356745 | 24391751 | 8.24E-10 |
| chr22 | 24361307 | 24394610 | 8.24E-10 |
| chr22 | 24362919 | 24394702 | 8.24E-10 |
| chr22 | 24364207 | 24395247 | 8.24E-10 |
| chr22 | 24364370 | 24395349 | 8.24E-10 |
| chr22 | 24364457 | 24395507 | 8.24E-10 |
| chr22 | 24364879 | 24395557 | 8.24E-10 |
| chr22 | 24375374 | 24410185 | 8.24E-10 |
| chr22 | 24388823 | 24423270 | 8.24E-10 |
| chr22 | 24390495 | 24423904 | 8.24E-10 |
| chr22 | 24391751 | 24427601 | 8.24E-10 |
| chr22 | 24467138 | 24495348 | 8.24E-10 |
| chr22 | 24467189 | 24495380 | 8.24E-10 |
| chr22 | 24468157 | 24495973 | 8.24E-10 |
| chr22 | 24489637 | 24511501 | 8.24E-10 |
| chr22 | 24489977 | 24511635 | 8.24E-10 |
| chr22 | 24491234 | 24511896 | 8.24E-10 |

| | | | |
|---|---|---|---|
| chr22 | 24493238 | 24511937 | 8.24E-10 |
| chr22 | 24496321 | 24513817 | 8.24E-10 |

**Supplementary Table 9: 4C primers**

| Locus | Forward primer name | Forward primer sequence | Reverse primer name | Reverse primer sequence | Assayed feature | Viewpoint (hg19) | | |
|---|---|---|---|---|---|---|---|---|
| Chr3 | LSM3_FW1 | CTACACGACGCTCTTCCGATCT CATCGACATATAAGAGATC | LSM3_RW2 | CAGACGTGTGCTCTTCCGATCGAT TAGGCCATGGTGACT | LSM3 Gene promoter | chr3 | 14218841 | 14220714 |
| Chr22 | MMP11_enh3_FW2 | CTACACGACGCTCTTCCGATCT CTCTGAGGAGAGCTGATC | MMP11_enh3_RW2 | CAGACGTGTGCTCTTCCGATCCTG AGGTGTGGGCATAGT | Enhancer Chr22 locus | chr22 | 24167718 | 24172058 |
